# Mapping-by-sequencing reveals genomic regions associated with seed quality parameters in *Brassica napus*

**DOI:** 10.1101/2022.06.01.494149

**Authors:** Hanna Marie Schilbert, Boas Pucker, David Ries, Prisca Viehöver, Zeljko Micic, Felix Dreyer, Katrin Beckmann, Benjamin Wittkop, Bernd Weisshaar, Daniela Holtgräwe

## Abstract

Rapeseed (*Brassica napus* L.) is an important oil crop and harbours the potential to serve as a highly productive source of protein. This protein exhibits an excellent amino acid composition and has a high nutritional value for humans. Seed protein content (SPC) and seed oil content (SOC) are two complex quantitative and polygenic traits which are negatively correlated and assumed to be controlled by additive and epistatic effects. A reduction of seed glucosinolate (GSL) content is desired as GSLs cause a stringent and bitter taste. The goal here was the identification of genomic intervals relevant for seed GSL content and SPC/SOC. Mapping-by-sequencing (MBS) revealed 30 and 15 new and known genomic intervals associated with seed GSL content and SPC/SOC, respectively. Within these intervals we identified known but also so far unknown putatively causal genes and sequence variants. A 4 bp insertion in the *MYB28* homolog on C09 shows a significant correlation with a reduction in seed GSL content. This study provides insights into the genetic architecture and potential mechanisms underlying seed quality traits, which will enhance future breeding approaches in *B. napus*.

## 1. Introduction

Rapeseed (*Brassica napus*, AACC, 2n = 38) is the second most important oil crop after soybean [1]. *B. napus* is a recent allopolyploid species formed by hybridization between *B. oleracea* and *B. rapa* followed by chromosome doubling around 7500 years ago [2]. In addition to its nutritionally beneficial seed oil composition, *B. napus* has the potential to serve as high quality protein source. Protein rich meals are a valuable “by-product” of oil extraction and can be used as viable source for plant protein due to its high quality and production volume [3]. The increasing demand for vegetable protein and oil requires breeding efforts in order to enhance the yield of protein and oil in *B. napus*. Dry mature seeds are composed of oil (45-50% w/w) and protein (20-25% w/w) [4]. Lipids are stored in the form of triacylglycerols (TAGs) in oil bodies, while seed storage proteins like cruciferin and napin are deposited in protein bodies or protein storage vacuoles [3]. *B. napus* protein isolates revealed a high bioavailability comparable to animal proteins like eggs or collagen and are rich in essential amino acids (EAAs) [5]. The high amounts of bioactive compounds in vegetable proteins have beneficial effects on human health by e.g. preventing hypertension, scavenging free radicals, and reducing cardiovascular risk factors [6].

The presence of off-taste components such as glucosinolates (GSLs) hinders the use of *B. napus* protein for food production [6]. GSLs are sulfur- and nitrogen rich secondary plant metabolites [7] and their degradation products have various functions, e.g. in protection against pathogens and herbivorous insects [8-11]. GSLs are classified into aliphatic, aromatic or indolic GSLs and their biosynthesis can be divided into three stages: I) chain elongation of selected precursor amino acids, II) synthesis of the core GSL structure, and III) secondary modifications of the amino acid side chain [12]. Transcriptional regulation of GSL biosynthesis is controlled by subgroup 12 of R2R3-MYB transcription factors [13-18]. In *A. thaliana, MYB28/HAG1* and *MYB29/HAG3* positively regulate aliphatic GSL biosynthetic genes [15,17], while *MYB34/ATR1, MYB51/HIG1*, and *MYB122/HIG2* control indolic GSL biosynthesis [14,16,18]. As aliphatic GSLs represent 91%-94% of total seed GSL content, while indolic GSLs contribute 5%-8% to the total seed GSL content in *B. napus* [19], the relevance of *MYB28/HAG1* and *MYB29/HAG3* homologs for controlling seed GSL content rises. In general, modern double low *B. napus* varieties display 10-15 μmol GSL per g seed instead of 60-100 μmol GSL per g seed in old varieties [4].

Three major loci controlling GSL content located on *B. napus* chromosome A09, C02, and C09 have been described [20-24]. These QTL are co-localized with the three homologs of *A. thaliana MYB28* and have thus been proposed as candidate genes responsible for the phenotypic variation (PV) of GSL content [2,20,23]. The two homologs on chromosome C02 and C09 are absent in the *B. napus* Darmor-bzh reference genome sequence [2]. The deletion of these two *MYB28* homologs was identified as cause for low GSL content [20]. Additional candidate genes for lowering GSL content were identified in an analysis of various *Brassica* genome sequences [23]. However, the link between base-pair level sequence variations and seed GSL content has not been confirmed or analysed in great detail yet.

Seed protein content (SPC) and seed oil content (SOC) are negatively correlated traits controlled by multiple genes assumingly involving epistatic and additive effects [25]. Consistent with the polygenic origin of SPC and SOC, several studies have reported different genomic intervals distributed across all linkage groups potentially involved in SPC/SOC control [26-35]. Major factors controlling SPC and SOC are additive effects ranging typically from 0.27-2.04% per individual QTL [25,30,31,33-35] and environmental conditions [26,35-37]. While the majority of previous studies reported frequently minor QTL, which explained individually ∼1.2%-19% of the PV in SPC and SOC [26,28-32,35], four studies reported less frequent major QTL explaining up to 20%-30.2% of the PV [26,33-35]. The chromosomal positions of SOC QTL differ between *B. napus* cultivars [31,34,35,38].

Due to the agronomic and economic importance of SPC and SOC, a better knowledge about the underlying regulatory network is of high relevance for future breeding strategies. The negative correlation of SPC and SOC challenges the breeding aim to simultaneously increase oil and protein content. Improvement of seed quality traits during breeding can be achieved by using the variability of naturally or induced mutations and interspecific hybridization among *B. napus* species [4]. However, breeding elite varieties e.g. by backcrossing techniques can take years. The rapid development of high-throughput sequencing technologies promote the application of approaches like mapping-by-sequencing (MBS) for the rapid identification of causal mutation underlying a phenotype of interest [39,40]. MBS is a fast and cost effective way to develop superior crop cultivars with desirable traits as demonstrated in various crops [41-46]. Reference genome sequences like the *B. napus* Darmor-bzh reference genome sequence [2] provide the basis for MBS approaches. With the rise of third generation sequencing technologies, long-read assemblies with a high continuity of *B. napus* cultivars like Zheyou7 became recently available [47-49].

In this study, genomic intervals associated with SPC, SOC, and GSL content in *B. napus* were analysed using a large segregating F2 population via MBS. This population was derived from a cross of the *B. napus* winter type cultivars Lorenz and Janetzkis Schlesischer. Furthermore, candidate genes were identified by incorporating transcriptomic data sets. Correlation and gene expression studies indicated that a 4 bp insertion located in a *MYB28* homolog on chromosome C09 is a major factor controlling seed GSL content. Sequence variants identified in here will facilitate the development of genetic markers for breeding programs in *B. napus*.

## 2. Materials and Methods

### 2.1 Plant material and trait measurement

The phenotypically segregating F2 population, designated L-x-JS, consists of 2323 individuals and was derived from a cross between the parental lines Lorenz (P1) and Janetzkis Schlesischer (P2), both are *B. napus* winter type rape varieties. Janetzkis Schlesischer (DOI: 10.25642/IPK/GBIS/288477) has a high seed GSL content of ∼90 μmol/g FW and contains erucic acid. Lorenz is listed as a variety for diversity with the accession number RAW 2152 (https://pgrdeu-preview.ble.de/tsorten/steckbrief/id/551533) and displays 00-quality. It has medium-high grain yields, high oil content and low GSL content (maintaining institute: Norddeutsche Pflanzenzucht Hans-Georg Lembke KG, DE005). 1373 F2 individuals were planted in Granskevitz alias growing area 2 (SPC_A2) (GE, GPS: 54.526908°, 13.21998°) and 948 in Asendorf alias growing area 1 (SPC_A1) (GE, GPS: 52.7724145°, 9.0044643°). The plants were grown in accordance with German legislation. In total, 1951 F2 individuals of the L-x-JS F2 population were used for the genotype and phenotype analyses for seed GSL content, while 2315 individuals were used for SPC due to the higher variance of SPC.

Seeds were collected for the GSL, SPC and SOC measurements via near-infrared reflectance spectroscopy (NIRS) and analysed in triplicates. The NIRS measurement was carried out with intact-seed samples. The measured trays were designed for high-throughput measurement of oil seed rape. Each tray requires a seed volume of about 2 cm^3^. The samples were scanned by a Polytec PSS-2121 diode array spectrometer (Polytec GmbH, Waldenbronn, Germany) with 256 pixels. Reflectance was measured in the range from 1,100 to 2,100 nm with a step size of 2 nm recorded with the software PSSHOP (Polytec) using DSV internal calibration. Calibration and validation procedures were carried out with several Software packages (Senso*Logic* GmbH, Norderstedt, Germany). Calibration performance was verified periodical with independent validation sets.

Individuals for sequencing were selected based on phenotypic data, DNA-quality, and cultivation location. For the GSL pools individuals grown in SPC_A1 and SPC_A2 were used to build the high and the low GSL pool. For the SPC pools, individuals grown in SPC_A2 were used to build the high and the low pool.

### 2.2 DNA extraction and pooling

Whole-genomic DNA was extracted from leaf disks using the CTAB method [50]. The low GSL pool consisted of 38 genotypes (GSL_L, <30.83 μmol/g dry weight), while the high GSL pool contained 52 genotypes (GSL_H, >70 μmol/g dry weight) (Figure 2). For growing area 2 (SPC_A2), 22 genotypes were used for the low protein pool (SPC_L_A2 low pool, <16.0% total dry mass, Figure 2) and 19 genotypes for the high protein pool (SPC_H_A2 high pool, >23.1% total dry mass) (Figure 2). Library preparation and pooling strategy was performed as described before [41]. The GSL pools were sequenced on a HiSeq1500 in highoutput mode using four lanes and the 2 × 100 PE scheme, while the SPC pools were sequenced on a HiSeq1500 in rapid mode using two lanes and the 2 × 150 PE scheme. Lorenz and Janetzkis Schlesischer were sequenced on a HiSeq1500 using the 2 × 150 PE scheme.

### 2.4 Mapping and variant calling

Read quality was checked with FastQC [51]. Reads were mapped via BWA-MEM v0.7 [52] to the *B. napus* Darmor-bzh v4.1 genome sequence [2] and the Zheyou7 assembly [47] (File S1). Default parameters were applied and the –M flag was set to avoid spurious mappings. Mapping statistics were calculated via the flagstat function of samtools [53] prior and past the following filtering step. Mappings were cleaned with samtools view -q 30 -b -F 0×900 -f 0×2 to remove low quality alignments and reads without a properly mapped mate. The filtered BAM files were passed to GATK v3.8 [54-56] for the identification of a variant set based on hard filtering. BWA-MEM and GATK were chosen due to excellent performances in previous studies [41,57].

### 2.5 Generation of the “gold standard”for SNV filtering

The workflow starts with the generation of a gold standard for SNV filtering, which contains SNVs which are homozygous in the parental genotypes and heterozygous in the reconstituted F1 (File S2, File S3). First the reads of the parents were mapped to the *B. napus* Darmor-bzh genome sequence v4.1 and variants were called as described above. Next, coverage filters based on the BAM and VCF files were applied. BAM-derived coverage files were constructed as described in Pucker *et al*. 2018 [58]. A minimum coverage of 10 and a maximum coverage of 60 were determined to yield high quality SNVs which are likely not affected by copy number variations (GitHub filter_parent_variants.py). The upper limit was chosen, because it represents twice the modal value of each file. Triallelic variants and variants present in both parents were excluded from further analyses as these are not contrasting between the pools (File S2) (GitHub combine_homo_VCFs_vs_Bn41.py).

The resulting set of homozygous SNVs of the parental genotypes was then screened for heterozygosity in a reconstituted F1 population (File S3) (GitHub filter_vcf_F1.py) to generate the final gold standard which contained 903,253 SNVs (File S2, File S4) (GitHub merge_vcfs.py). The reconstituted F1 variant set comprises variants derived from all analysed genomic sequencing data of our study. Only “heterozygous” variants, which showed an allele frequency between 0.2-0.8 against the *B. napus* Darmor-bzh genome sequence were used for the down-stream filtering (File S2).

### 2.6 Filter raw variants per pool for delta allele frequency (dAF) calculation

A sophisticated filtering approach was applied to select only highly reliable SNVs for the downstream analyses (File S2). High quality SNVs were extracted from the raw variants of each pool (GSL_L, GSL_H, SPC_H, SPC_L) by considering only SNV positions present in the gold standard (File S2) (GitHub filter_pools_vcfs_for_gold_standard.py). Only variants with a minimum 0.75 times the average median coverage and a maximum coverage of 1.5 times the average median coverage per pool were kept. This final set of variants of the high and low GSL and SPC_A2 pools, respectively, were used to calculate delta allele frequencies (dAFs) (File S2, File S5, File S6) (GitHub combine_single_VCFs_version3.py). The dAF is defined as the absolute difference between the allele frequency (AF) values from the two pools for a given variant position. Only biallelic variants are included for the calculation of dAFs to facilitate a reliable dAF estimation.

### 2.7 Interval detection

For interval detection, Fisher’s exact test was applied on the raw SNVs of the pools to yield variants with a significant dAF (Figure 1). A *p*-value cut-off of 0.05 was applied after correction for multiple testing (GitHub fisher_exact_test_corrects_for_multiples_testing.py). The passing SNVs are called “statistically meaningful differential Allele-specific Read Counts” (dARCs) (File S7, File S8).

**Figure 1:**
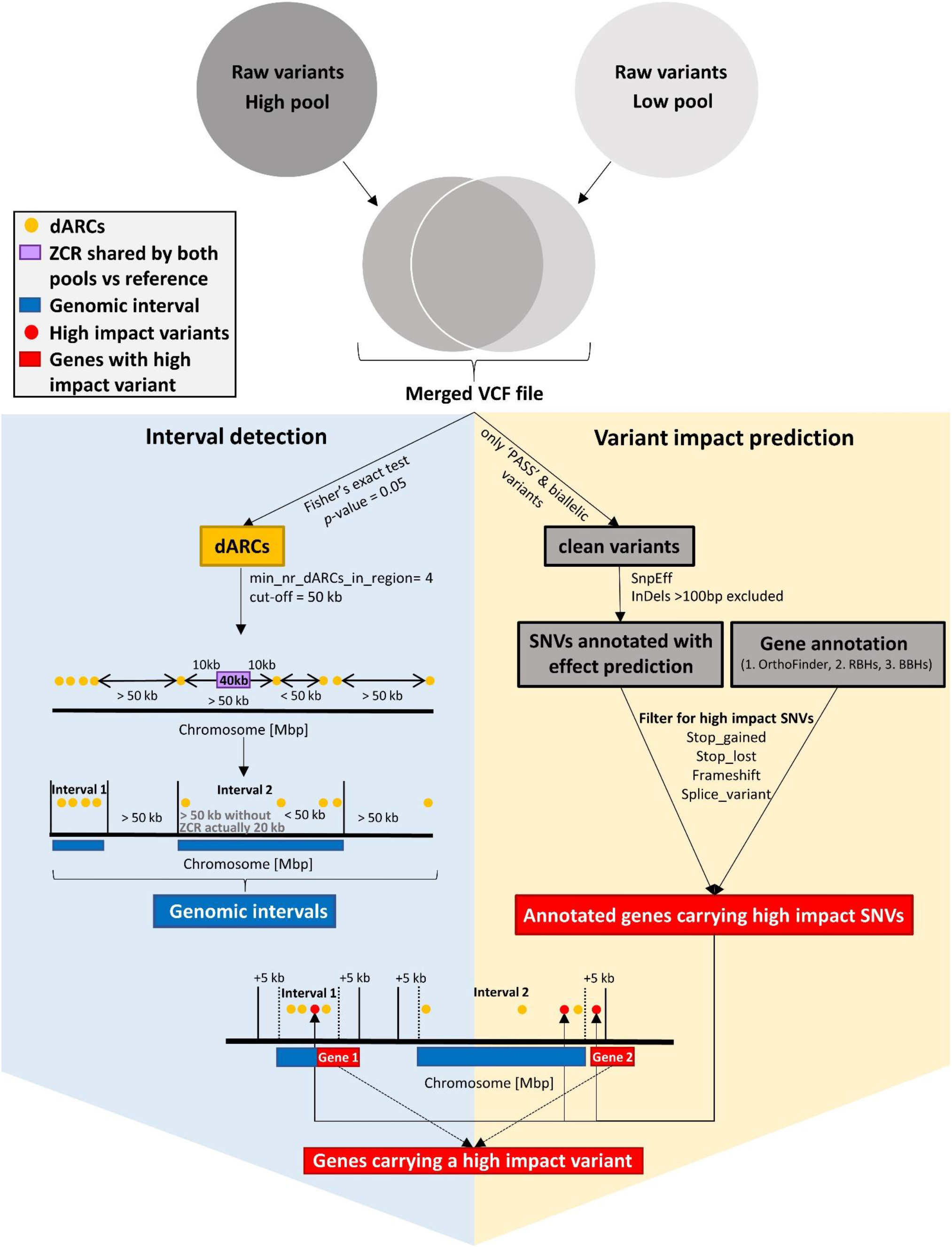
Schematic illustration of interval detection and extraction of genes carrying high impact variants. First both raw variants of each pool are combined into a merged VCF file. Interval detection was performed based on the density of dARCs (left). A set of clean variants were extracted from the merged VCF file and used for the detection of high impact SNVs (right). Finally, the results of both approaches are integrated (bottom). Raw variants (grey circles); dARCs (yellow points); ZCR (purple); genomic intervals (dark blue rectangles); high impact variants (red points); genes carrying high impact variants (red rectangles).

These dARCs were used to identify genomic intervals associated with the analysed traits (Figure 1) (Table 2, Table 5). The following criteria were applied: I) The minimum amount of dARCs in an interval was set to 4 (--min_nr_dARCs_in_reg), II) the distance between at least 3 dARCs of one interval needs to be greater than 1 kbp (--dis_in_reg), and III) distance between any two adjacent dARCs must be less than 50 kbp (--dis_out_reg) (GitHub get_intervals_based_on_dARCs_Bn41_v4.py). While a certain number of dARCs is required to seed an interval, it is also important that these are equally distributed. Numerous variants originating from the same sequenced DNA fragment could be due to an artifact and are excluded by requiring a minimal distance of the seed dARCs. To avoid extremely large intervals with low dARC frequencies between dARC rich intervals, the 50 kbp cut-off for the dARC distance is intended to split intervals without a constantly high dARC density.

Zero coverage regions (ZCRs) were identified by using the coverage information of both pools and applying a genome-wide screening with a window size of 200 bp per chromosome (GitHub PAV_finder.py). ZCRs are considered during the interval detection, as they are often responsible for splitting genomic intervals into parts (Figure 1) (File S9, File S10). The localisation of ZCRs at the same genomic position in both pools prevents the detection of variants and hence no dARCs can be detected.

Finally, detected genomic intervals were ranked according to their amount of dARCs. Initial candidate genomic intervals were manually inspected to find a suitable cut-off. For seed GSL content and SPC genomic intervals containing at least 100 dARCs and 65 dARCs were used for downstream analysis, respectively (Table 2, Table 5).

### 2.8 Generation of dAF plots

Noise in the genome-wide dAF plots (File S11, File S12) was reduced through the combination of adjacent dAFs (calculated as described in 2.6, GitHub sophisticated_cov_plot.py). Variants within a sliding window of 100 variants were represented by the median dAF of all variants in the window. Each step was a 5 variant shift of the window. The genome-wide distribution of “statistically meaningful differential Allele-specific Read Counts” (dARCs) (compare 2.7 for details) was visualised by the normalized density of dARCs. The normalized density of dARCs was calculated by combining the amount of dARCs in sliding windows of size 100 kbp with steps of size 30 kbp divided by the total amount of SNVs within this window. In addition, the mean mapping coverage of the pools using the same sliding window parameters were calculated and normalized to the maximum mean coverage per chromosome for visualisation (File S11, File S12).

### 2.9 Presence-absence variations (PAVs)

PAVs were identified based on the BAM derived coverage files by comparison of coverage information of both pools in a genome-wide window approach which considers annotated genes (GitHub PAV_finder.py). Genomic regions or genes with no or at least extremely low coverage in one pool, but substantially higher coverage in the other pool were considered as PAVs. The coverage was normalized to the overall coverage of the pool. Genes located on the genetically non-anchored random scaffolds were excluded from this analysis. For the identification of PAVs based on gene regions the following parameters were used: PAV_finder.py was used in gene mode and –mincov was set to 10 (File S13, File S14).

### 3.0 Functional annotation and candidate genes

Genes located within or spanning over the borders of the identified genomic intervals were extracted (Figure 1). Genes were functionally annotated by transferring the Araport11 functional annotation to the v5 gene models [2]. OrthoFinder v2.3.7 [59] was applied using default parameters to identify orthogroups between representative peptides of *A. thaliana* Araport11 as previously defined [60], and the *B. napus* representative peptide sequences derived from the *B. rapa, B. oleracea, B. napus* Express 617, Darmor-bzh, Lorenz, and Janetzkis Schlesischer (File S1, File S15, File S16, File S17, File S18, File S19). Remaining unannotated genes were functionally annotated by using reciprocal best blast hits (RBHs) and best blast hits (BBHs) as described previously [61] (Figure 1). We refer to the Bna genes that were annotated as homologs of the respective *A. thaliana* genes.

### 3.1 Variant impact prediction via SnpEff

Variants predicted to have an impact on the genes located within the genomic interval were extracted. First, SnpEff v4.1f [62] was applied on the merged variants of each pool (GitHub combine_single_VCFs_for_SnpEff.py), which passed GATK’s quality filters (‘PASS’) (Figure 1). The resulting VCF was subjected to SnpEff with default parameters using a custom database constructed from the *B. napus* Darmor-bzh v4.1 genome sequence and the v5 annotation, which were corrected for the used frame (File S20). SnpEff results were filtered for “high impact” variants as previously defined [61], which included predictions of loss or gain of a stop codon mutations, frameshift mutations, and splice site variants (GitHub get_intervals_based_on_sig_snps_Bn41_v4.py) (Figure 1). Finally, genes located within +/-5 kbp of the borders of a genomic interval were analysed for predicted high impact variants (Figure 1) (File S21, File S22).

### 3.2 Generation and analysis of RNA-Seq data

Seeds and leaves (28 days after flowering (DAF)) RNA-Seq samples of Janetzkis Schlesischer (P2) and seeds and leaves (23 and 35 DAF) RNA-Seq samples of *B. napus* SGDH14 (medium-high seed GSL content) [63] were prepared and sequenced according to Schilbert *et al*. 2021 [64]. Additionally, RNA-Seq reads derived from seeds and leaves of *B. napus* Express 617 [64] and public RNA-Seq data sets (File S23) were mapped to the *B. napus* Darmor-bzh v4.1 and Zheyou7 assemblies using STAR v.2.7.1a [65]. STAR was run in basic mode allowing maximal 5% mismatches and requiring an alignment spanning at least 90% of the read length. Mapping statistics were calculated based on STAR.log files via a customized python script (GitHub parse_STAR_log_file_create_mapping_statistic.py) (File S24). We used featureCounts v1.5.0-p3 [66] for the generation of count tables. The mean fragments per kilobase exon per million reads (FPKM) or mean counts per million (CPM) expression values per organ were used for downstream analysis (GitHub generate_figures_only_mean_expression_calc.py and map_mean_exp_to_cand_genes_in_reg.py). For example, mean CPM expression values of Janetzkis Schlesischer, as well as average coverage information per pool were assigned to the genes to infer PAVs between pools (File S25, File S26) (GitHub fetch_gene_IDs_from_gff3_file.py, map_mean_exp_to_cand_genes_in_reg.py, map_PAVs_to_genes_in_regs.py).

### 3.3 Identification of MYB homologs

MYB homologs were identified with KIPEs as described previously [67]. KIPEs was run with a minimal BLAST hit similarity of 40% to reduce the number of fragmented peptides derived from possible mis-annotations. As bait peptide sequences, all *A. thaliana* MYBs were used [13]. As subject species, the proteomes of several *Brassica* species were used (File S1, File S17, File S19, File S27, File S28). The alignment was constructed with MAFFT v.7 [68] and trimmed to minimal alignment column occupancy of 10%. Next, a phylogenetic tree was build (https://GitHub.com/bpucker/script_collection/tree.py) with FastTree v2.1.10 [69] using 10,000 rounds of bootstrapping, including the identified MYB homologs from several *Brassica* species and well described MYB sequences from literature (File S29, File S30, File S31, File S32, File S33). The phylogenetic trees were visualized with FigTree v1.4.3 (http://tree.bio.ed.ac.uk/software/figtree/) (File S34, File S35). Additional previously identified MYB sequences derived from Darmor-bzh [70] were added manually to the phylogenetic tree to ensure completeness of MYB homologs.

By analysing mappings of genomic sequence reads from the parental genotypes against the Darmorbzh and Zheyou7 assembly, the copy numbers of *BnaMYB* genes/alleles involved in GSL biosynthesis were manually inspected via IGV [71]. Generally, the numbering of GSL MYBs is based on Seo *et al*. 2017 [70] with small modifications. Finally, *BnaMYB28_2* alleles from the *B. napus* cultivars of a subset of 100 lines from the BnASSYST diversity panel [72] were validated via PCR and Sanger sequencing (File S36).

## 3 Results

### 3.1 Phenotyping of the segregating F2 population

A large F2 population segregating for seed quality traits and consisting of over 2000 individuals derived from a cross between the *B. napus* cultivars Lorenz (P1) and Janetzkis Schlesischer (P2) was used for MBS (Figure 2). The traits studied were seed GSL content and SPC/SOC. The seed GSL content ranged from 11.8-88.1 μmol/g, while the SPC and SOC ranged between 9.7-28.0% and 24.3-56.3%, respectively (Figure 2).

**Figure 2:**
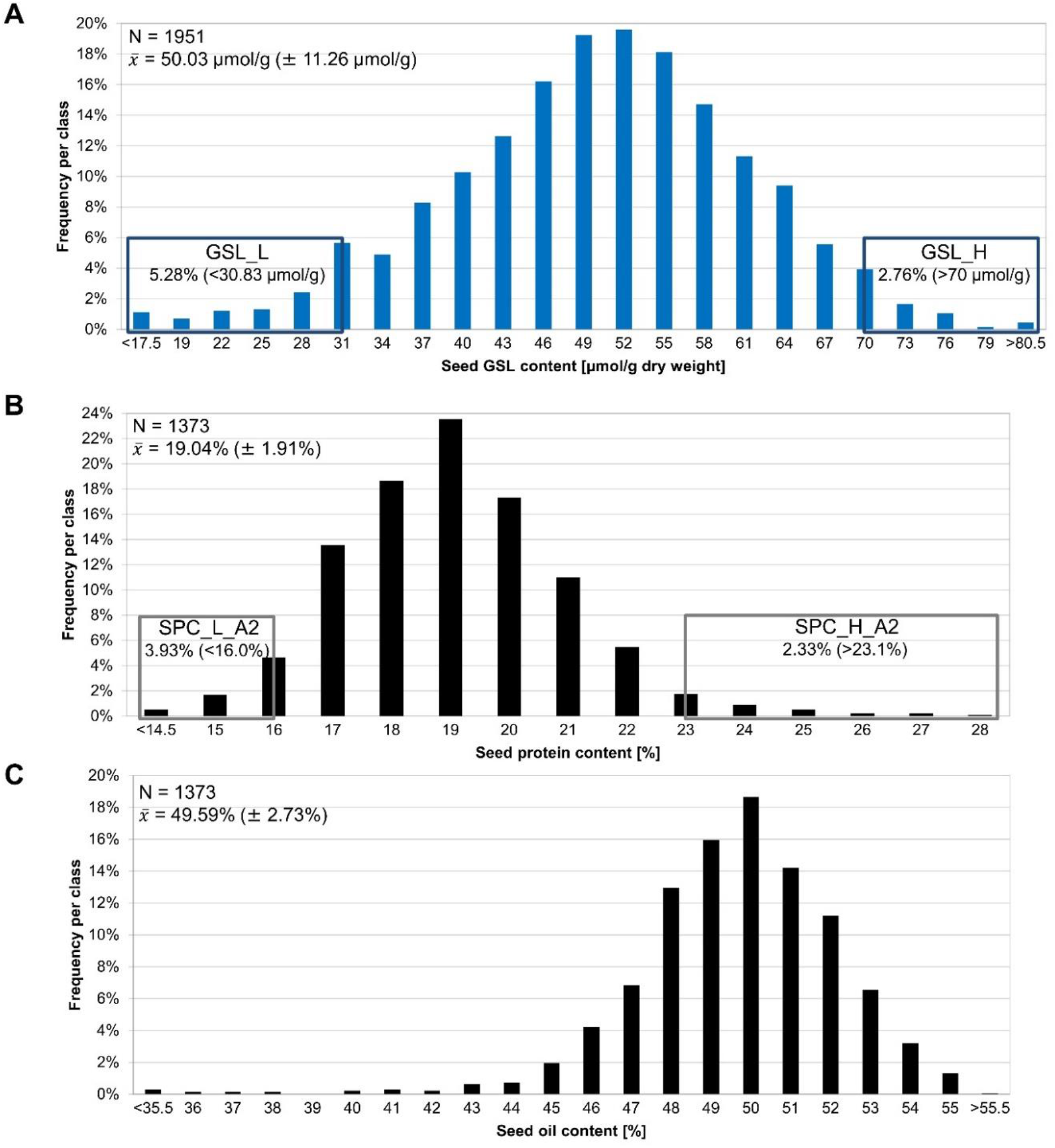
Distribution of traits in the F2 population. (A) Distribution of seed GSL content, (B) seed protein content, and (C) seed oil content of the segregating F2 population. Seeds of individual F2 plants were harvested and phenotyped via near-infrared resonance spectroscopy (NIRS). The sample size and mean of the distribution are given by N and 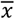, respectively. The rectangles in (A) and (B) mark the tails of the distributions used for pool building, e.g. the GSL low-(GSL_L), GSL high-(GSL_H), SPC low-(SPC_L_A2) and SPC high-(SPC_H_A2) pool. As SPC and SOC are negatively correlated the SPC and SOC pools are largely congruent and thus only the SPC pools were marked. The tails relevant for building the contrasting GSL pools account for 5.28% (<30.83 μmol/g) for the GSL_L pool and 2.76% (>70 μmol/g) for the GSL_H pool of the whole F2 population. The tails from the SPC distribution used to build the pools comprise 3.93% (<16.0%) for SPC_L_A2 and 2.33% (>23.1%) for SPC_H_A2 of the whole F2 population.

#### MBS predicted candidate genomic intervals controlling SPC, SOC, and seed GSL content

### 3.2 Mapping

To map candidate genes, pools from the F2 population were subjected to MBS analysis. Both parental genotypes P1 and P2 and the two pools representing individuals with extreme phenotypes were sequenced, both for seed GSL content and SPC/SOC. After read mapping to the *B. napus* Darmor-bzh genome sequence, low quality alignments and reads without a properly mapped mate were removed. Among these data sets at least 52% to 62% of the reads per data set were confidently mapped (File S37). See supplementary file S37 for mapped read depth values (File S37).

### 3.3 Variant calling

Variant calling revealed between 3,580,759 to 5,215,492 high quality variants (InDels and single-nucleotide variants (SNVs)) for the respective samples (Table 1). Of these, 2,632,505 (73.5 %) to 3,987,788 (76.5 %) variants were distributed on the 19 pseudochromosomes. The remaining variants were distributed on the genetically non-anchored random scaffolds and were excluded from further analysis. The raw variants of each pool were filtered for the gold standard SNVs, resulting in high quality SNVs sets of 889,280 SNVs for the SPC pools and 880,842 for the GSL pools (∼1,036-1,053 SNVs per Mbp) (Table 1). These SNVs were used to generate delta allele frequency (dAF) plots of the high and the corresponding low pool (File S11, File S12). We noticed that the usage of statistically meaningful differential Allele specific Read Counts (dARCs; see M&M for details) dARCs resulted in less noisy interval detection when compared against dAF approaches (File S11, File S12).

**Table 1:** Variant calling and dARCs statistics. For each data set the amount of called raw variants is given. Moreover, the number of variants in the gold standard is listed, as well as the number of SNVs left after filtering for the gold standard SNVs. Finally, the number of dARCs is stated.

| Samples | Raw variants | Gold standard SNVs |  |
| --- | --- | --- | --- |
| P1 | 3,580,759 | 903,253 (File S4) |  |
| P2 | 4,905,445 |  |  |
|  |  | SNVs left after filtering for gold standard | Statistically meaningful differential allele-specific read counts (dARCs) |
| SPC_A2 High-pool | 5,215,492 | 889,280 (File S6) | 8,407 |
| SPC_A2 Low-pool | 4,848,100 |  |  |
| GSL High-pool | 5,105,239 | 880,842 (File S5) | 20,726 |
| GSL Low-pool | 5,003,187 |  |  |

### 3.4 Genomic intervals and candidate genes associated with seed glucosinolate content

Evaluation of the dARC distribution among the pseudochromosomes allowed identification of 30 genomic intervals associated with seed GSL content (see M&M). These intervals were detected on six chromosomes, namely A02, A06, A09, C02, C07, and C09. Their sizes range from 73 kbp to 1.32 Mbp (Figure 3, Table 2). Out of the 30 intervals, 18 intervals are located on A09, five on C09, three on C02, two on C07, and one on A02 and A06 (Table 2). Several intervals in close proximity on one chromosome emerged due to the lack of dARCs located between these intervals. This can be caused by e.g. I) regions with low numbers of SNVs, II) low quality variants that do not qualify as dARCs, and III) a combination of the two mentioned causes.

**Figure 3:**
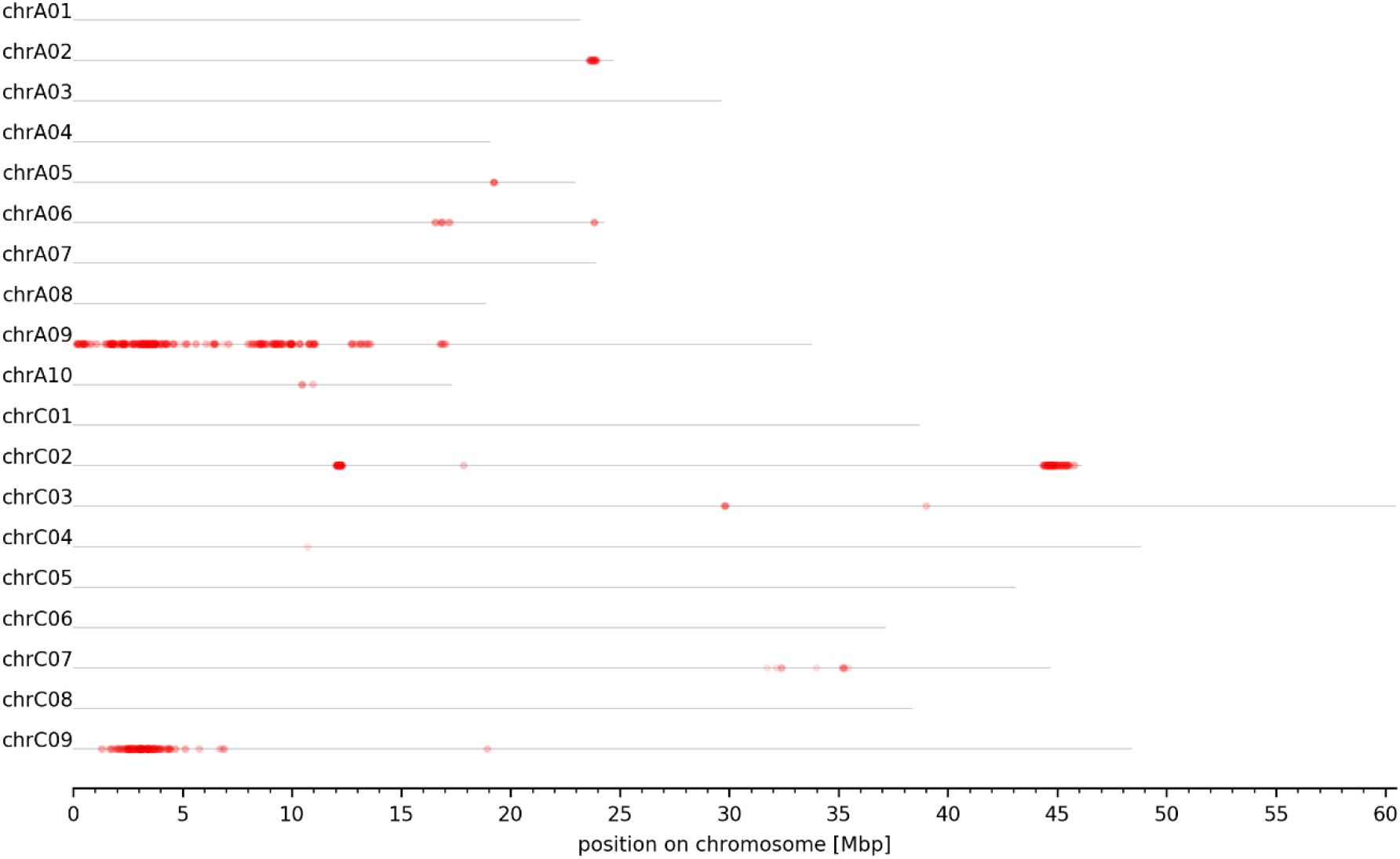
Genome-wide plot of normalized dARC density for seed GSL content. The normalized density of dARCs is plotted across all pseudochromosomes of the *B. napus* Darmor-bzh genome sequence. A heatmap ranging from white to red represents the normalized density of dARCs, where a red colour represents a high amount of dARCs.

**Table 2:** Genomic intervals of seed GSL content. The chromosomal position, size, start and end per genomic interval are listed.

| Interval ID | Chromosome | Size [bp] | Start [bp] | End [bp] |
| --- | --- | --- | --- | --- |
| A02_GSL_1 | chrA02 | 326909 | 23675288 | 24002197 |
| A06_GSL_1 | chrA06 | 90191 | 16818388 | 16908579 |
| A09_GSL_1 | chrA09 | 1007698 | 28668 | 1036366 |
| A09_GSL_2 | chrA09 | 479317 | 1465924 | 1945241 |
| A09_GSL_3 | chrA09 | 73214 | 2071845 | 2145059 |
| A09_GSL_4 | chrA09 | 317624 | 2303455 | 2621079 |
| A09_GSL_5 | chrA09 | 248198 | 3036680 | 3284878 |
| A09_GSL_6 | chrA09 | 395239 | 3440941 | 3836180 |
| A09_GSL_7 | chrA09 | 308180 | 4056024 | 4364204 |
| A09_GSL_8 | chrA09 | 79540 | 4580338 | 4659878 |
| A09_GSL_9 | chrA09 | 270234 | 6924362 | 7194596 |
| A09_GSL_10 | chrA09 | 934210 | 8018931 | 8953141 |
| A09_GSL_11 | chrA09 | 215209 | 9150682 | 9365891 |
| A09_GSL_12 | chrA09 | 173036 | 9514676 | 9687712 |
| A09_GSL_13 | chrA09 | 247692 | 9859341 | 10107033 |
| A09_GSL_14 | chrA09 | 78703 | 10831700 | 10910403 |
| A09_GSL_15 | chrA09 | 117588 | 10993988 | 11111576 |
| A09_GSL_16 | chrA09 | 374266 | 12597247 | 12971513 |
| A09_GSL_17 | chrA09 | 710703 | 13041084 | 13751787 |
| A09_GSL_18 | chrA09 | 378004 | 16764142 | 17142146 |
| C02_GSL_1 | chrC02 | 247718 | 12063724 | 12311442 |
| C02_GSL_2 | chrC02 | 724820 | 44411662 | 45136482 |
| C02_GSL_3 | chrC02 | 431332 | 45205897 | 45637229 |
| C07_GSL_1 | chrC07 | 562637 | 32063223 | 32625860 |
| C07_GSL_2 | chrC07 | 204488 | 35236183 | 35440671 |
| C09_GSL_1 | chrC09 | 671097 | 1152540 | 1823637 |
| C09_GSL_2 | chrC09 | 1320909 | 1911629 | 3232538 |
| C09_GSL_3 | chrC09 | 195766 | 3410309 | 3606075 |
| C09_GSL_4 | chrC09 | 425478 | 3699696 | 4125174 |
| C09_GSL_5 | chrC09 | 199414 | 4285280 | 4484694 |

In total, 1,807 genes were found within the genomic intervals associated with seed GSL content (File S25). Some of these genes have a well-known function in GSL biosynthesis or breakdown. A homolog of *A. thaliana* methylthioalkylmalate synthase 1 (*AthMAM1*), *BnaC02g41790D*, is involved in GSL biosynthesis and is part of the genomic interval C02_GSL_2. Moreover, homologs of APS kinase (*AthAPK*) and n-(methylsulfinyl)alkyl-glucosinolate hydroxylase (*AthAOP3*), *BnaA09g08410D* and *BnaA09g01260D*, involved in GSL biosynthesis were identified in the genomic intervals A09_GSL_1 and A09_GSL_7, respectively. The thioglucoside glucohydrolase homolog of *AthTGG, BnaA09g08470D*, involved in GSL breakdown was found in the genomic interval A09_GSL_7. Moreover, key transcription factors involved in the regulation of GSL content were located within the genomic intervals. Two close homologs of *AthMYB28* (*AthHAG1*), *BnaC09g05300D* and *BnaC09g05290D*, were located within the genomic interval C09_GSL_2 (File S25). In addition, homologs of *AthMYB34* (*AthATR1*), *BnaC09g05060D* and *BnaC02g41860D*, were identified in the genomic intervals C09_GSL_2 and C02_GSL_2, respectively (File S25).

#### 3.4.1 Glucosinolate associated MYB genes contributed by P1 and P2

To determine which parental genotype brings in which of the key transcription factor genes, we set out to identify all *BnaMYB* genes/alleles involved in GSL biosynthesis (*MYB28, MYB29, MYB34, MYB51, MYB122*, see introduction; collectively referred to here as ‘*B. napus* GSL MYBs’) in P1 and P2. Since it turned out that not all *BnaMYB* homologs are resolved in the *B. napus* Darmor-bzh genome sequence, we used in addition the long-read assembly of the *B. napus* cultivar Zheyou7 which covers more *BnaMYB* homologs. *BnaMYB* sequences of various genotypes including both parental genotypes were subjected to a phylogenetic analysis (File S29, File S35). By analysing mappings of genomic sequence reads from the parental genotypes against both assemblies, the copy numbers of *B. napus* GSL MYBs were identified (Table 3, File S29, File 15).

**Table 3:** *B. napus* GSL MYB gene copies identified in the parental genotypes. The number of the GSL MYB genes identified in the parental genotypes *B. napus* Janetzkis Schlesischer (P2) and *B. napus* Lorenz (P1) is listed.

|  | Lorenz (P1) | Janetzki Schlesiischer (P2) |
| --- | --- | --- |
| *BnaMYB28* | 4 | 5 |
| *BnaMYB29* | 4 | 4 |
| *BnaMYB34* | 8 | 7 |
| *BnaMYB51* | 7 | 7 |
| *BnaMYB122* | 6 | 5 |

A tandem gene duplication event of *BnaMYB122_2* in P1 resulted in a higher number of *BnaMYB122* genes compared to P2 (File S35). The additional *BnaMYB28* copy in P2, *BnaMYB28_5*, is likely to be derived from the loss of this copy in P1 as indicated by the fractionated and extremely low coverage of the C02 *BnaMYB28_5* by genomic read mappings of P1 reads to the Zheyou7 assembly (File S29, File S15, Table 3, Table 4). This was also supported by the analysis of GSL pools, where *BnaMYB28_5* revealed a ∼ 3 higher genomic coverage in the high GSL pool compared to the low GSL pool, indicating that this locus is only inherited by the high GSL parent P2.

**Table 4:** *BnaMYB28* homologs. The *BnaMYB28* homologs identified in the *B. napus* cultivars Zheyou7, Darmor-bzh, Janetzkis Schlesischer (P2), and Lorenz (P1) are listed.

| Name | Zheyu7 | Darmor-bzh | Janetzki Schlesiſcher (P2) | Lorenz (P1) |
| --- | --- | --- | --- | --- |
| *BnaMYB28_1* | *BnaC07T0355800ZY* | *BnaCnng43220D* | Present in genomic mapping | Present in genomic mapping |
| *BnaMYB28_2* | *BnaC09T0054800ZY* | *BnaC09g05300D*<br>+ *BnaC09g05290D* | Present in genomic mapping | Present in genomic mapping |
| *BnaMYB28_3* | *BnaA03T0422000ZY* | *BnaA03g40190D* | Present in genomic mapping | Present in genomic mapping |
| *BnaMYB28_4* | *BnaA09T0074900ZY* | Deleted | Absent | Absent |
| *BnaMYB28_5* | *BnaC02T0362400ZY* | Deleted | Present in genomic mapping | Absent |
| *BnaMYB28_6* | *BnaA02T0409000ZY* | Non-functional copy | Present in genomic mapping | Present in genomic mapping |

Large deletions were detected on A09 in both, P1 and P2. Both deletions affect the presence or absence of *B. napus* GSL MYBs. The ∼920 kbp deletion of P2 ranges from ∼4.06 to 4.98 Mbp, while the ∼50 kbp deletion of P1 ranges from 4.46 to 4.51 Mbp (pseudochromosome positions taken from the Zheyou7 assembly). The P2 deletion A09_P2_920 harbours 163 genes, while only 1 gene (*BnaA09g05680D*) is located inside the shared deletion of P1 (File S38). Fractionated and extremely low coverage of the A09 *BnaMYB28_4* homolog was observed in genomic read mappings of P1 and P2 reads to the Zheyou7 assembly, indicating its deletion in both parental genotypes (Table 4, File 15). In addition to *BnaMYB28_4*, additional genes associated with GSL biosynthesis were identified in the A09_P2_920 deletion (File S38) which overlap with high ranked genes affected by PAVs (File S13). The *BnaMYB34_7* homolog is located within the A09_P2_920 deletion, but outside of the one of P1 (Table 3, Table 4, File 15). This was also supported by the analysis of GSL pools, where *BnaMYB34_7* revealed a ∼3 times higher genomic coverage in the low GSL pool compared to the high GSL pool, indicating that this locus is inherited by the low GSL parent P1. The A09_P2_920 deletion overlaps with the genomic intervals A09_GSL_4 and A09_GSL_5 and might also be the reason for additional genomic intervals detected in its proximity.

In order to identify *B. napus* GSL MYB genes expressed in seeds which could influence seed GSL content and thus explain the phenotypic variation in the high and low GSL pool, we analysed their expression in seeds and leaves of P2. Most *BnaMYB28, BnaMYB29, BnaMYB34, BnaMYB51*, and *BnaMYB122* homologs were not or very low expressed in leaves and seeds (Figure 4). Only five homologs are expressed in seeds: *BnaMYB28_2, BnaMYB28_5, BnaMYB34_1, BnaMYB51_2*, and *BnaMYB51_6*. As *BnaMYB28_5* is absent in the low GSL parent P1 (Table 4) but expressed in P2, this homolog might explain the genomic intervals identified in the south of chromosome C02 (Figure 3, Table 2). Supporting these genomic intervals, *BnaMYB34_1* is also located in the south of C02 (File S29). *BnaMYB51_2* and *BnaMYB51_6* are located on C08 and A08 respectively and have homologs in both parental genotypes which showed no genomic coverage differences between the high and low GSL pool (File S15). However, *BnaMYB28_2* on C09 exceeds the expression of all *B. napus* GSL MYB homologs by a factor of at least 2-3 fold in leaves and seeds (Figure 4). The analysis of over 650 public available *B. napus* RNA-Seq data sets (File S23) supports the high expression of *BnaMYB28_2* compared to all other GSL MYBs across various tissues and environmental conditions (File S39).

**Figure 4:**
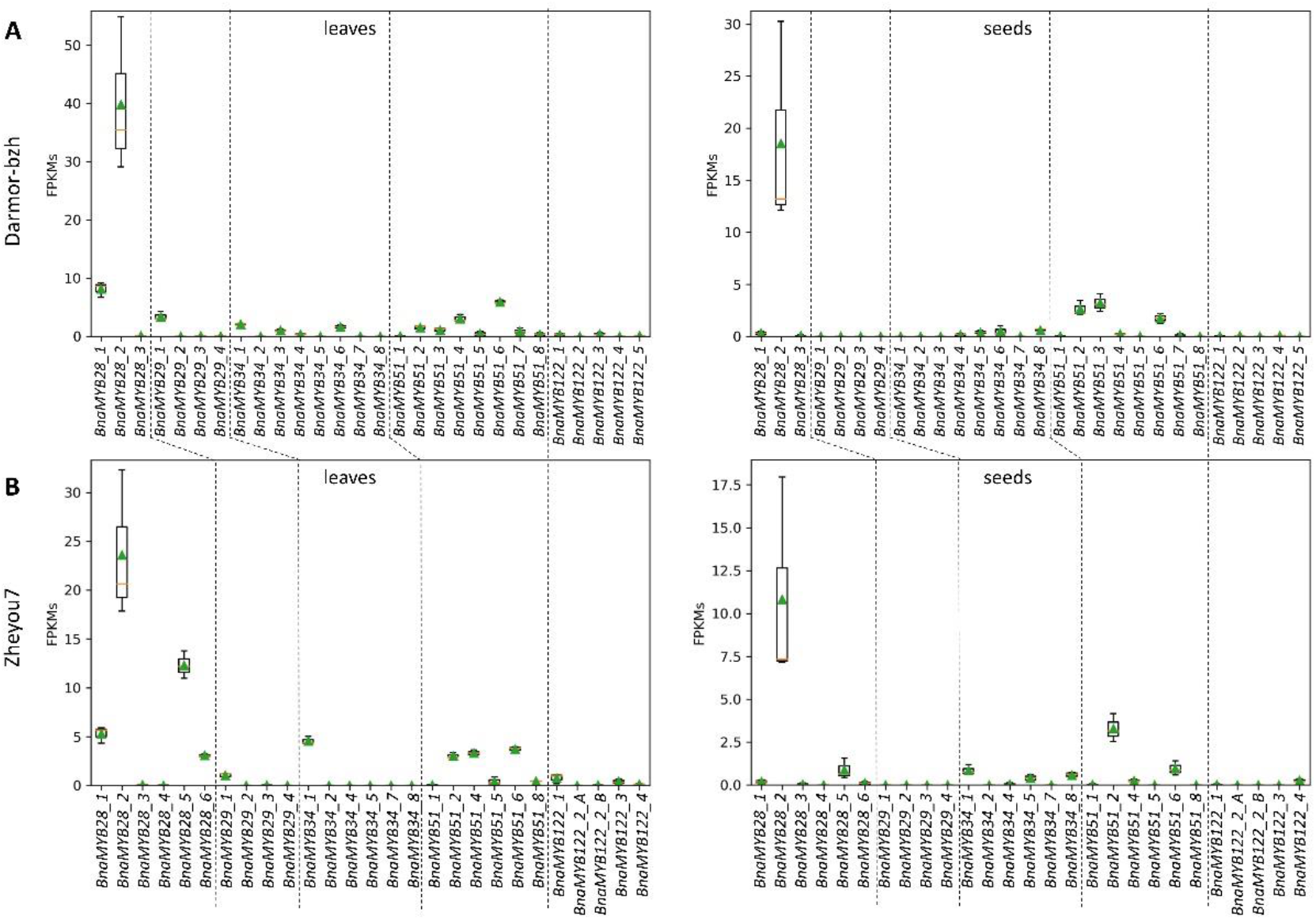
Expression of *B. napus* GSL MYB homologs. The expression of *BnaMYB28, BnaMYB29, BnaMYB34, BnaMYB51*, and *BnaMYB122* homologs in leaves and seeds of P2 on the basis of the (A) Darmor-bzh and (B) Zheyou7 genome sequence, as well as their annotation is displayed. The value displayed for *BnaMYB28_2* on the basis of Darmor-bzh is the average FPKM of both annotated fragments at this locus, *BnaC09g05290D* and *BnaC09g05300D*. The analysis was performed on the basis of the Darmor-bzh and Zheyou7 assembly to ensure that all *B. napus* GSL MYB homologs are represented. For example, although the *BnaMYB28_5* homolog on C02 is missing in the Darmor-bzh genome sequence, the allele of *BnaMYB28_5* in P2 could be assigned based on the corresponding Zheyou7 sequence. FPKMs = fragments per kilobase million; n=3.

#### 3.4.2 Variation effects in genes involved in seed glucosinolate biosynthesis

As sequence variants can influence the function of gene products, the impact of sequence variants on genes located within or near the genomic intervals (+/-5 kbp) was predicted. Interestingly, the highly expressed *BnaMYB28_2* located on C09 is affected by a 4 bp insertion (GCTA) near the end of the annotated third exon (Figure 5A, File S15, File S40). The phylogeny of *MYB28* homologs across several Brassicaceae species revealed that the ancestral allele did not contain this 4 bp insertion (Figure 5A and B, File S35), i.e. the ancestral allele encodes a functional MYB transcription factor. The Darmor-bzh genome sequence contains the insertion. The 4 bp insertion results in a premature stop codon of the *MYB28* homolog *BnaC09g05300D* leading to a truncated protein (Figure 5B). The second fragment of this locus is annotated as *BnaC09g05290D* which encodes only a *MYB28* C-terminal fragment (Figure 5A).

**Figure 5:**
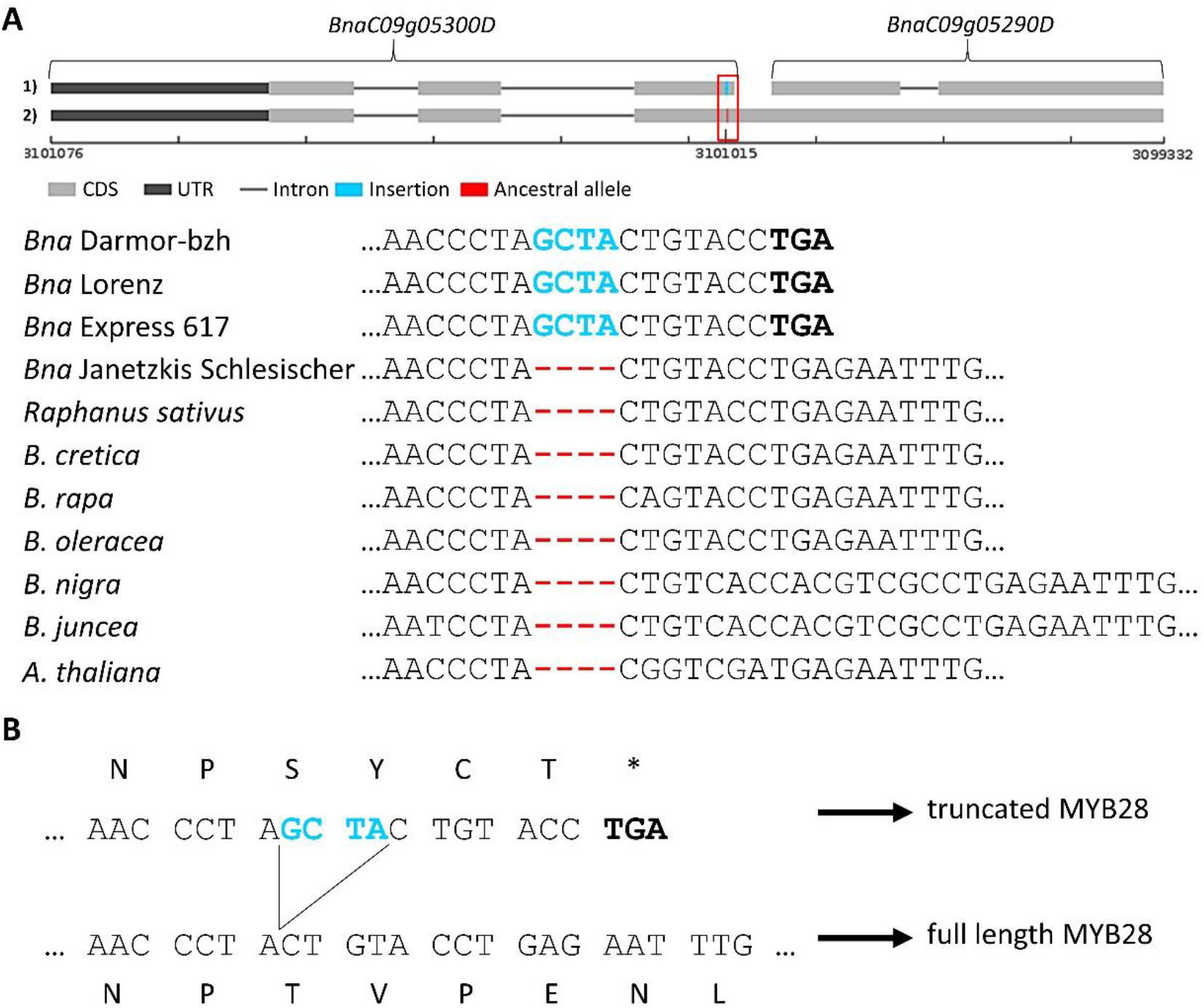
A 4 bp insertion is associated with the inactivation of *BnaMYB28_2* which is strongly expressed in seeds. (A) Genomic structure of *BnaC09g05300D* and *BnaC09g05290D* and alignments of several Brassicaceae *MYB28* homologs. The stop codon is shown in bold. (B) Translated reading frame stressing the stop codon derived from the 4 bp insertion. The 4 bp insertion is shown in blue, while the ancestral allele is marked in red.

The ancestral allele is present in 73 % of the GSL high pool reads and 29 % of the GSL low pool reads, resulting in a dAF of 0.44 (File S21, File S40). The genomic reads of the high GSL parent P2 showed the ancestral allele, while those of the low GSL parent P1 carried the insertion (Figure 5A, File S40). RNA-Seq data from leaves and seeds of P2 support the presence of the ancestral allele on transcript level (File S40).

The BnASSYST diversity panel was screened to investigate the correlation of the 4 bp insertion with low seed GSL content. The two alleles of the *BnaMYB28_2* C09 homolog, namely *BnaMYB28_2_1\** describing the ancestral allele and *BnaMYB28_2_2\** describing the insertion allele, were validated by this analysis and confirmed by sequencing (Figure 6A). The insertion allele *BnaMYB28_2_2\** was identified to be significantly correlated with low seed GSL content (Figure 6B). Moreover, co-segregation of the insertion with the C09 homolog was detected (Figure 6B). This finding is in accordance with our phylogenetic analysis, supporting the assumption that the allele without the 4 bp insertion is the ancestral allele.

**Figure 6:**
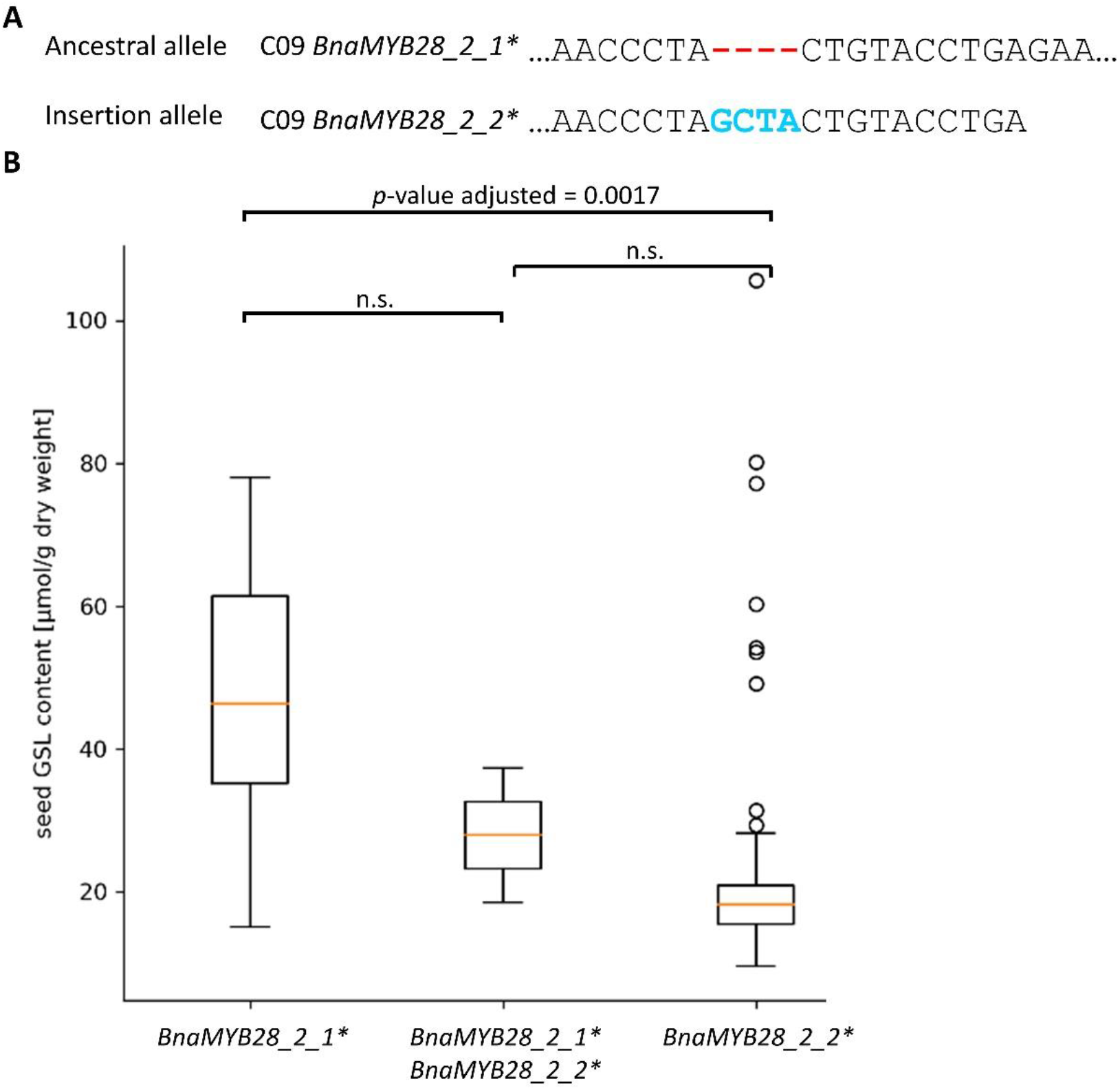
The 4 bp insertion is associated with low seed GSL content. (A) The two C09 *BnaMYB28_2* alleles showing the ancestral allele and the 4 bp insertion allele. (B) Boxplots for seed GSL content based on the genotypes derived from the BnASSYST diversity panel (n=100): n=90 for *BnaMYB28_2_2\**; n=2 for *BnaMYB28_2_2\** + *BnaMYB28_2_1\**; n=8 for *BnaMYB28_2_1\**. Differences between genotypes were analysed by Mann-Whitney-U-test and corrected for multiple testing. n.s. represents not significant.

In order to evaluate whether the 4 bp insertion might be ubiquitously associated with low seed GSL content in *B. napus*, we analysed its presence across several *B. napus* lines (File S41). In accordance with our previous findings, all high GSL lines contain at least one functional *BnaMYB28* homolog harbouring the ancestral allele as it has been observed for example for the *B. napus* genotype SGDH14. All low GSL lines revealed the presence of the insertion allele of the C09 *BnaMYB28* homolog, while the A09 *BnaMYB28* homolog was absent (File S41).

#### 3.4.3 PAVs

Additional candidate genes were identified via PAV analysis, which revealed 316 genes affected by PAVs (File S13). As seed GSL content is a polygenic trait, it is not expected to identify genes with no read coverage in one pool compared to full coverage in the other pool. In this study, genes likely to be deleted in one parent but present in the other resulted in a 1/3 read coverage ratio. Genes predicted to be present in the low GSL parent P1, but absent in the high GSL parent P2 are described. By analysis of the chromosomal positions of high ranked PAVs, two deletions on A09 were identified. The first major ∼900 kbp deletion on A09 (A09_P2_920) and its associated candidate genes have already been described above. A second ∼25 kbp deletion is located within A09_GSL_13 ranging from ∼10.028-10.053 Mbp, namely A09_P2_25 (File S42). Three of four genes located within this deletion are involved in abscisic acid (ABA) signaling in *A. thaliana*. Namely, a homolog of the calcium-dependent protein kinase 32 *AthCPK32, BnaA09g16660D*, as well as two homologs of *BURNOUT1* (*AthBNT1*), *BnaA09g16680D* and *BnaA09g16690D*, were identified to be absent in P2 compared to P1. For the fourth gene no functional annotation was available.

### 3.5 Genomic intervals, candidate genes and variation effects associated with seed protein and oil content

In total 15 genomic intervals associated with SPC and SOC were identified on chromosome A01, A06, A09, C03, C04, C08, and C09 and their sizes range from 10.5 kbp to 2.07 Mbp (Figure 7, Table 5). Out of these 15 intervals, five intervals are located on C08, three on A06, two on C04 and C09 and one on A01, A09 and C03 (Table 5). 351 genes were located within the genomic intervals (File S26), of which some have a well-known function in lipid and/or protein biosynthesis. In addition, several SNVs affecting genes associated with SPC and SOC were investigated.

**Figure 7:**
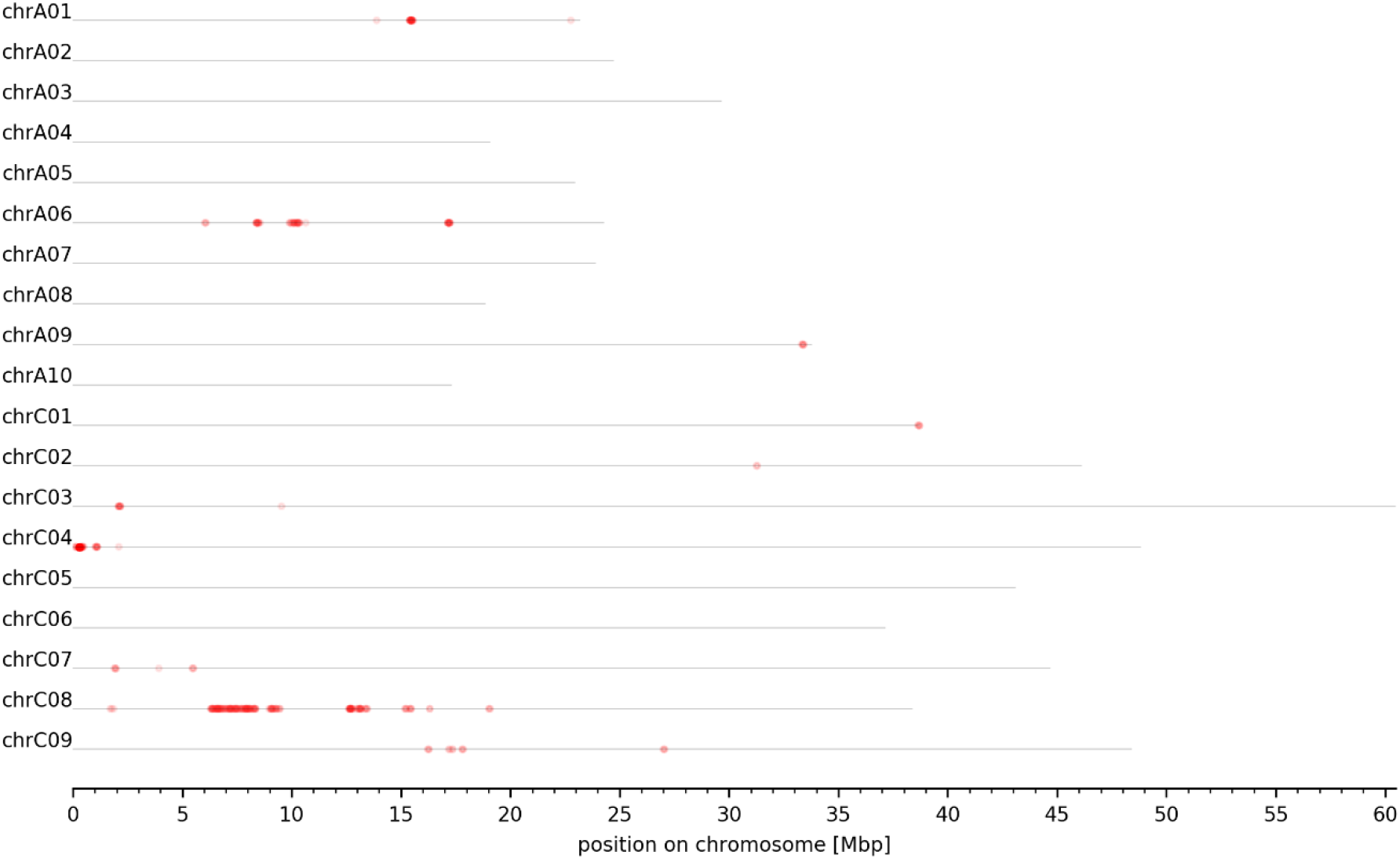
Genome-wide plot of normalized dARC density for SPC and SOC. The normalized density of dARCs is plotted across all pseudochromosomes of the *B. napus* Darmor-bzh genome sequence. A heatmap ranging from white to red represents the normalized density of dARCs, where a red colour represents a high amount of dARCs.

**Table 5:** Genomic intervals of SPC and SOC content. The chromosomal position, size, start and end per genomic interval are listed.

| Interval ID | Chromosome | Size [bp] | Start [bp] | End [bp] |
| --- | --- | --- | --- | --- |
| A01_SPC_1 | chrA01 | 143410 | 15427093 | 15570503 |
| A06_SPC_1 | chrA06 | 247164 | 8434596 | 8681760 |
| A06_SPC_2 | chrA06 | 72097 | 9931989 | 10004086 |
| A06_SPC_3 | chrA06 | 410672 | 10177027 | 10587699 |
| A09_SPC_1 | chrA09 | 41585 | 33415524 | 33457109 |
| C03_SPC_1 | chrC03 | 86028 | 2142025 | 2228053 |
| C04_SPC_1 | chrC04 | 53519 | 364385 | 417904 |
| C04_SPC_2 | chrC04 | 10677 | 1104377 | 1115054 |
| C08_SPC_1 | chrC08 | 252499 | 1743278 | 1995777 |
| C08_SPC_2 | chrC08 | 2072658 | 6332262 | 8404920 |
| C08_SPC_3 | chrC08 | 472053 | 9033205 | 9505258 |
| C08_SPC_4 | chrC08 | 464034 | 12719350 | 13183384 |
| C08_SPC_5 | chrC08 | 138554 | 13409507 | 13548061 |
| C09_SPC_1 | chrC09 | 92091 | 16262887 | 16354978 |
| C09_SPC_2 | chrC09 | 381823 | 17275714 | 17657537 |

A homolog of altered seed germination 2 (*AthASG2*), *BnaC03g04570D*, is located in C03_SPC_1 and affected by a frameshift mutation which is present in 86% of the low SPC pool reads and in 21% of the high SPC pool reads (File S22). Within C04_SPC_1 a homolog of ATP binding cassette subfamily B4 (*AthABCB4*), *BnaC04g00490D*, was predicted to carry a frameshift mutation with a prevalence of 91% of the low SPC pool reads and 15% of the high SPC pool reads. In C04_SPC_2 a homolog of a kinase family protein with an ARM repeat domain (*AthCTEXP*), *BnaC04g01520D*, was predicted to gain a stop codon which is present in 93% of the low SPC pool reads and in 21% of the high SPC pool reads (File S22). In C08_SPC_2, two major candidate genes were detected. First, a homolog of the well-known regulator of seed oil content phospholipase D delta (*AthPLDδ*), *BnaC08g05680D*, is affected by a mutation of the splice site of the first to the second exon (File S22). This mutation is present in 67% of the low SPC pool reads and in 18% of the high SPC pool reads. Second, a homolog of the serine carboxypeptidase-like 41 (*AthSCPL41*), *BnaC08g05590D*, carries a frameshift mutation in 59 % of the low SPC pool reads and was not detected in the high SPC pool reads (File S22). *BnaC08g05680D* and *BnaC08g05590D* are located next to each other in the *B. napus* Darmor-bzh genome sequence. In addition, a frameshift mutation in a homolog of an amino acid transporter, *BnaC08g02490D*, (C08_SPC_1) and mutations in homologs responsible for post-translational protein modifications, e.g. ubiquitylation (*BnaA06g15510D, BnaA06g18030D, BnaA06g18370D, BnaA06g18380D*) (A06_SPC_1 and A06_SPC_3) or myristoylation (*BnaA06g15490D*) (A06_SPC_1) were identified in genomic intervals on chromosome A06 (File S22). A homolog of the candidate gene F-box protein 7 (*AthFBP7*), *BnaA06g15510D*, (A06_SPC_1) was predicted to gain a stop codon due to a SNV in 78% of the low SPC pool reads and 14% of the high SPC pool reads (File S22). Another candidate gene is the homolog of RAB GTPase homolog 8A (*AthRABE1c*), *BnaA06g18220D* (A06_SPC_3), which is affected by a frameshift and stop gained mutation in the high SPC pool and are not present in the low SPC pool (File S22). PAVs genes located near or in the genomic intervals were not associated with SPC/SOC based on the functional annotation of their corresponding *A. thaliana* homolog (File S14). In summary, these candidate genes are proposed to contribute to the variations of SPC and SOC in *B. napus*.

## 4. Discussion

We investigated a segregating F2 population to identify genomic intervals and candidate genes associated with protein, oil, and glucosinolate content of *B. napus* seeds. The genomic intervals and candidate genes identified in this study should provide deeper insights into the genetic architecture of the three complex traits. We envision that the results of this study will be used for genetic improvement of seed quality in *B. napus*.

### 4.1 Seed oil and protein content

Control of the multigenic traits SPC and SOC is complex and previous studies have reported various SPC and SOC QTL with the majority being minor QTL distributed across all linkage groups [28-35]. As expected for the multigenic traits SPC and SOC, we identified several genomic intervals distributed across 7 chromosomes: A01, A06, A09, C03, C04, C08, and C09. A large proportion of the genomic intervals overlap with loci associated with SOC from previous studies, such as A01_SPC_1, A06_SPC_1-3, C08_SPC_4-5 [27,30,35] indicating the high reliability of these loci. The genomic interval A01_SPC_1 overlaps with a significant region associated with amount of eicosenoic acid [27]. All intervals on chromosome A06 are in line with significant regions associated with oleic acid and linoleic acid [27]. Chao *et al*. identified several QTL for SPC and SOC, of which two QTL for SOC are in proximity to the genomic interval C03_SPC_1 and one QTL for SPC overlaps with C08_SPC_3-5 [26]. Moreover, C08_SPC_4 and C03_SPC_1 are located in proximity to SNVs significantly associated with linolenic acid [27,38], while C08_SPC_5 is close to SNVs significantly associated with oleic acid, erucic acid [27,38], and eicosenoic acid [27].

Numerous candidate genes and sequence variants associated with SPC/SOC have been detected. For example, a homolog of the candidate gene phospholipase D delta, *BnaC08g05680D* is located in C08_SPC_2. Phospholipases are involved in lipid degradation, membrane reconstruction and signal transduction [73]. PLD*δ*, one of the most abundant PLDs, hydrolyses phospholipids to phosphatidic acid (PA) [73]. Devaiah *et al*. showed significant reduced seed germination for the *pldδ A. thaliana* and attenuation of *PLDα1* expression might improve oil stability, seed quality and seed aging [74]. In leaves of *pldδ A. thaliana* mutants the suppression of *PLDδ* results in the attenuation of PA formation, which blocks the degradation of membrane lipids retarding ABA-promoted senescence [75]. Another candidate gene is serine carboxypeptidase-like 41 (*SCPL41*), whose *B. napus* homolog is located in C08_SPC_2. *SCPL41* was identified as negative regulator of membrane lipid metabolism and is proposed to be required for phospholipid metabolism or PA-dependent signaling in *A. thaliana* [76]. Deletion of *SCPL41* increased total leaf lipid content and phosphatidylcholine, phosphatidylethanolamine, and phosphatidylglycerol contents, which are substrates of phospholipid hydrolysis via PLD [76]. Interestingly, *PLDδ* and *SCPL41* are located next to each other in the *B. napus* genome sequence indicating they might be functionally related or act in the same network as it has been observed for e.g. biosynthetic gene clusters in *A. thaliana* [77]. In the low SPC pool the *B. napus PLDδ* and *SCPL41* homologs are affected by high impact variants. Therefore, the most likely non-functional *SCPL41* and *PLDδ* homologs might result in an increase of total lipid content in the low SPC pool. Due to the negative correlation of SPC and SOC this would in turn lead to a low SPC.

On chromosome C04 *CTEXP* and *ABCB4* homologs have been identified as candidate genes affected by nonsense and frameshift mutations, respectively, with a high prevalence in the low SPC pool. Homologs of *CTEXP* are known to play a role in intracellular protein trafficking [78], while *ABCB4* was identified as an auxin efflux transporter [79]. However, members of the same enzyme family are known as intracellular sterol transporters in mice [80].

*A. thaliana* mutants of the candidate gene *ASG2* located in C03_SPC_1 show seeds with an increased oil body density, fatty acid content, and weight [81]. The authors hypothesize that ASG2 modulates the gene expression or activity of ω-6-fatty acid desaturase (FAD2) and/or ω-3-fatty acid desaturase (FAD3), which are involved in the production of unsaturated FAs [81]. Thus *ASG2* might be a novel candidate gene contributing to an increased SOC in the low SPC pool. Moreover, the candidate genes involved in posttranslational protein modifications might influence SPC/SOC content as e.g. myristoylation enables protein-lipid interactions and controls the transport and localization of proteins [82].

The candidate gene F-box protein 7 located in A06_SPC_1 is affected by a nonsense mutation in the low SPC pool, and *A. thaliana fbp7* mutants display a defect in protein biosynthesis after cold and heat stress [83]. *FBP7* is proposed to regulate translation through ubiquitylation and thereby inactivates a translation repressor under temperature stress [83]. The nonsense mutation in *BnaA06g15510D* might results in a non-functional *FBP7* leading to activation of the translational repressor and thus in a reduction of protein content.

Located in A06_SPC_3 a *RABE1c* homolog was affected by several mutations in the high SPC pool. Peroxisomal fatty acid-oxidation is the main pathway for seed lipids catabolism [84]. RABE1c is responsible for peroxin 7 (PEX7) dislocation/degradation on the peroxisome membrane and mutation of *RABE1c* restored peroxisomal β-oxidation activity and *PEX7* expression [84]. Treatment with proteasome inhibitors also restored endogenous PEX7 protein levels in GFP-PEX7-expressing seedlings [84]. Thus, a mutated *RABE1c* in the high SPC pool might decrease SOC and in parallel increase SPC by increased peroxisomal β-oxidation activity and proteasome inhibitory-like characteristics, respectively.

Of the genes located in the genomic interval on chromosome A01 no association with SPC or SOC was detected based on the functional annotation. However, Liu *et al*. identified a significant SNV located within *BnaA01g22680D*, which is in proximity to the genomic interval A01_SPC_1 [85]. The *A. thaliana* homolog is mildew resistance locus O 6 (*AthMLO6*). Besides the in this study identified candidate genes, unknown genes or genomic components might be involved in trans-regulatory or epistatic interactions of SPC/SOC which may be responsible for the indicated genomic intervals.

### 4.2 Seed glucosinolate content

Seed GSL content is influenced by several major and minor QTL. In this study, loci controlling seed GSL content were identified on chromosome A09, C09, C02, A02, A06, and C07 being in accordance with previous findings [20-24]. All significant SNPs located in the regions on A09, C02, C07 and C09 explained 56.7% of the cumulative phenotypic variance [22].

We identified several genomic intervals on chromosome A09. In this case a large interval is subdivided into several intervals because of I) regions with low numbers of SNVs, or II) low quality variants or III) a combination of both. Regions with low numbers of SNVs can be caused by deletion in one or both parental genotypes. Therefore, dARCs cannot be detected in these regions which results in a subdivision of intervals. It is possible that such intervals are flanking loci associated with the trait of interest e.g. in the case of large deletions. Two deletions in the high GSL parent P2 may have a major influence on seed GSL content. The first ∼900 kbp deletion (A09_P2_920) causes the loss of *BnaMYB28_4* and *BnaMYB34_7*, the *A. thaliana* homologs of these two genes are known positive regulators of GSL biosynthesis. However, because *BnaMYB28_4* is also absent in the low GSL parent P1 and *BnaMYB34_7* is not expressed in the high GSL parent P2 seeds, these genes are likely not responsible for the observed variation in seed GSL content between the parents. The A09_P2_920 deletion also causes the loss of *BnaA09g05810D* and *BnaA09g05510D* annotated as *CALNEXIN1* (*At5g61790*) and *COBRA* (*At5g60920*), respectively. CALNEXIN1 and COBRA were identified as interaction partner of the aliphatic GSL pathway specific enzyme CYP83A1 [86]. *A. thaliana calnexin1* and *cobra* homozygous T-DNA insertion mutants revealed an increased total aliphatic and indolic GSL content [86], indicating that they have a negative influence on GSL biosynthesis. Thus, the deletion of both genes in P2 might contribute to its high GSL content.

The second ∼25 kbp deletion of P2 (A09_P2_2) was found to cause the loss of genes which are involved in abscisic acid (ABA) signaling in *A. thaliana*. Plant hormones like ABA, jasmonic acid (JA) and salicylic acid (SA) impact indolic and aliphatic GSL biosynthesis by increasing the expression of GSL transcription factors like *MYB28* and *MYB29* and *vice versa* [18,87,88]. The deletion of the two homologs (*BnaA09g16680D, BnaA09g16690D*) of *AthBURNOUT1* could result in increased levels of plant stress hormones, as *A. thaliana burnout1* loss of function mutants overproduce stress hormones such as JA, SA, ABA, and ethylene [89]. Moreover, the *A. thaliana* homolog of the deleted *BnaCPK32* (*BnaA09g16660D*) is a positive regulator of *AthABF4*, which positively regulates the expression of ABA-responsive genes to increase stress tolerance [90]. Thus, lacking *BnaCPK32* might impair ABA signaling and stress tolerance, which might be compensated by a high GSL content. Taken together, the possible increase in plant stress hormones and reduced stress tolerance might boost GSL production in P2.

In addition to the detection of trait-associated genomic intervals and large deletions, our approach enables the identification of single candidate genes and pin-points sequence variants and domains at the base-pair level which are associated with seed GSL content. Seed GSL content is influenced by multiple genes involved in the biosynthesis of aliphatic and indolic glucosinolates, as well as GSL breakdown and transport [12,23,91]. However, seed GSLs can be largely decreased by reducing aliphatic GSLs as they represent 91%-94% of total seed GSL content [19]. According to Kittipol *et al*. the results of some studies lead to the assumption that inhibition of GSL transport processes cause the low seed GSL trait in *B. napus* as no significant correlation between leaf and seed GSL could be found. However, Kittipol *et al*. showed that seed and leave aliphatic GSL content is most likely regulated by a master regulator affecting all plant tissues rather than long-distance transport, because no accession with high leaf and low seed GSL content was identified [92]. Thus the positive regulator of the aliphatic GSL biosynthesis, MYB28, was proposed as master regulator [92] and we could indeed identify specifically the *BnaMYB28_2* homolog on chromosome C09 as major regulator of seed GSL content. *B. napus* lines with a low GSL content carry a 4 bp insertion in this gene which causes a premature stop codon thus leading to a most likely non-functional MYB transcription factor. Consequently, structural genes in the GSL biosynthesis are no longer activated due to the lack of this central transcriptional regulator. In contrast, lines with a high GSL content carry a functional *BnaMYB28_2* allele. A correlation between this 4 bp insertion and low seed GSL content was observed before [93], but not explained mechanistically. The Damor-bzh genome sequence harbours the 4 bp insertion which caused the prediction of two gene models at this locus, namely *BnaC09g05300D* and *BnaC09g05290D*. While *BnaC09g05300D* contains a R2R3-MYB DNA-binding domain on sequence level, *BnaC09g05290D* does not [70]. This could explain why previous studies [92,93] have not described the molecular consequences of this insertion. Our findings on the genomic level are supported by RNA-Seq analyses which show a strong expression of *BnaMYB28_2* in seeds. The A09 *BnaMYB28* homolog might contribute to a high seed GSL content in some high GSL lines, but was absent in all low GSL lines investigated in this study (File S41) marking the 4 bp insertion of the C09 homolog as key determinant for seed GSL content phenotype. Interestingly, the 4 bp insertion is located in the middle of a QTL for seed GSL content which explained 48% of the phenotypic variation [94]. Furthermore, the importance of the *MYB28* C09 homolog as positive regulator of GSL biosynthesis was demonstrated in *B. oleracea* varieties [95,96]. Yi *et al*. analysed the expression of 81 genes involved in GSL biosynthesis in 12 genotypes of four *B. oleracea* subspecies across leaves, stems, and florets [95]. Interestingly, out of five aliphatic transcription factors-related genes only the C09 *MYB28* homolog (*Bol036286*) was expressed in all genotypes, again stressing its essential role in aliphatic GSL biosynthesis. The data also confirm that not all GSL MYB transcription factors need to be expressed to produce GSLs [95].

Additional genomic differences controlling seed GSL content are copy number variations. As the *BnaMYB28* homolog on C02 (*BnaMYB28_5*) is absent in the low GSL parent P1, while being expressed in the seeds of in the high GSL parent P2, *BnaMYB28_5* could explain some of the phenotypic variation in GSL content. The role of the *MYB28* homologs on C02 and C09 in GSL biosynthesis was analysed by double knockout lines of *B. oleracea* [97]. The remaining functional MYB28 homolog on C07 and the two MYB29 homologs did not compensate the low GSL phenotype, indicating that these homologs play an inferior role in GSL production [97]. In line with our expression studies showing that the C02 homolog *BnaMYB28_5* is >12-fold higher expressed in leaves compared to seeds, the expression of the C02 *MYB28* homolog is assumed to be of most importance in regulating aliphatic glucosinolate biosynthesis in aerial organs [92]. Finally, *BnaMYB28_5* is much lower expressed in seeds compared to *BnaMYB28_2* supporting our hypothesis that the *BnaMYB28* C09 homolog is the major regulator of seed GSL content. Although these transcription factors appear responsible for a huge proportion of the GSL difference between *B. napus* lines, we have identified additional candidate genes associated with seed GSL content like *BnaC02g41790D* (homolog of *AthMAM1*) and *BnaA09g08410D* (homolog of *AthAPK*). In accordance with our findings, *BnaA09g01260D* (homolog of *AthAOP3*) and *BnaA09g08470D* (homolog of *AthTGG1*) were recently identified as novel candidate genes for seed GSL content [93].

## 5 Conclusions

We identified and described the molecular consequences of a 4 bp insertion located in the third exon of *BnaMYB28_2* on chromosome C09 as the most likely causative variant explaining the majority of the phenotypic variance in seed GSL content. *B. napus* lines with a low GSL content carry a 4 bp insertion in this gene which causes a premature stop codon leading to a most likely non-functional MYB. *BnaMYB28_2* is the only GSL transcription factor highly expressed in seeds as demonstrated in the high GSL parent P2 and other *B. napus* genotypes. Moreover, we identified several new candidate genes controlling SPC and SOC. The new insight into the molecular mechanisms of SPC, SOC, as well as seed GSL content can serve as useful target for the genetic improvement of *B. napus* seed quality traits.

## Supplementary Materials

The following files are available online at https://doi.org/10.4119/unibi/2963492

File S1: Data sets used for phylogenetic and genomic analysis.

File S2: Schematic illustration of the workflow for the generation of the gold standard and follow up delta allele frequency calculation.

File S3: Variants of the reconstituted F1.

File S4: Variants of the gold standard.

File S5: Seed GSL content SNVs left after filtered for the gold standard.

File S6: SPC SNVs left after filtered for the gold standard.

File S7: dARCs of seed GSL content pools.

File S8: dARCs of SPC pools.

File S9: ZCRs of seed GSL content pools.

File S10: ZCRs of SPC pools.

File S11: Genome-wide delta allele frequency plots of seed GSL content pools.

File S12: Genome-wide delta allele frequency plots of SPC pools.

File S13: PAVs of GSL pool analysis.

File S14: PAVs of SPC pool analysis.

File S15: IGV screenshot of GSL MYB homologs expression and genomic coverage in the GSL pools and the parental genotypes, as well as transcriptomic coverage of Janetzkis Schlesischer (JS) leaves and seeds RNA-Seq data.

File S16: CDS sequences from *B. napus* Lorenz.

File S17: Peptide sequences from *B. napus* Lorenz.

File S18: CDS sequences from *B. napus* Janetzkis Schlesischer.

File S19: Peptide sequences from *B. napus* Janetzkis Schlesischer.

File S20: Frame-corrected gff3 file for SnpEff analysis.

File S21: Predicted high impact variants of genes located within +/-5 kbp of the borders of the genomic intervals associated with seed GSL content.

File S22: Predicted high impact variants of genes located within +/-5 kbp of the borders of the genomic intervals associated with SPC content.

File S23: SRA IDs of the used public available RNA-Seq data sets from *B. napus*.

File S24: RNA-Seq mapping statistics.

File S25: Genes located in genomic intervals associated with seed GSL content.

File S26: Genes located in genomic intervals associated with SPC content.

File S27: CDS sequences from *B. napus* SGDH14.

File S28: Peptide sequences from *B. napus* SGDH14.

File S29: GSL MYB transcription factors homologs.

File S30: MYB amino acid sequences used for phylogenetic analysis.

File S31: GSL transcription factors’coding sequences of *B. napus* Janetzkis Schlesischer.

File S32: GSL transcription factors’coding sequences of *B. napus* Lorenz.

File S33: GSL transcription factors’coding sequences of *B. napus* SGDH14.

File S34: Phylogenetic tree based on the amino acid sequences of all MYB homologs identified in Brassica species marking the key GSL MYB transcription factor clade.

File S35: Phylogenetic tree based on the amino acid sequences of the key GSL transcription factors identified in Brassica species.

File S36: Oligonucleotide sequences.

File S37: Genomic read mapping statistics.

File S38: Gene located in the deleted region of *B. napus* Janetzkis Schlesischer.

File S39: Expression of GSL transcription factors based on public available *B. napus* RNA-Seq data sets.

File S40: Genomic and transcriptomic read mappings of GSL pools, parental genotypes, and Janetzkis Schlesischer (JS) leaves and seeds RNA-Seq data vs the Darmor-bzh genome sequence. (A) Whole gene view is presented, while (B) shows a close up of the position of the 4 bp InDel.

File S41: The C09 *BnaMYB28_2* and A09 *BnaMYB28_4* alleles association with seed GSL phenotype across various *B. napus* genotypes.

File S42: IGV screenshot of the ∼25 kbp deletion located within A09_GSL_13 in *B. napus* Janetzkis Schlesischer.

## Author Contributions

B.We. and D.H. conceived the project. H.M.S., B.P., D.R. and D.H. conducted data analysis. H.M.S. and B.P. wrote the initial draft manuscript. B.We. and D.H. supervised the project. Z.M., F.D., and K.B. performed the plant material and genotype selection, initiated and propagated the F2 crossing population and quantified GSL, oil and protein content in seeds. B.Wi. provided additional genotypes (the ERANET-Assyst *B. napus* diversity set) and related phenotypic data. P.V. prepared all sequencing libraries and generated all sequence data analysed in this study. All authors have read and agreed to the final version of this manuscript.

## Funding

The project was funded by the Federal Ministry of Education and Research of Germany (BMBF) under the grant numbers 0315957D (NuGGET) and 031B0198D (RaPEQ). We acknowledge support for the publication costs by the Deutsche Forschungsgemeinschaft and the Open Access Publication Fund of Bielefeld University.

## Institutional Review Board Statement

Not applicable.

## Informed Consent Statement

Not applicable.

## Data Availability Statement

All data generated in this study can be found under the ENA/NCBI Bioproject ID PRJEB36483. In detail, the Janetzkis Schlesischer and SGDH14 RNA-Seq data sets generated for this study can be found with the ENA/NCBI IDs: ERS11936124-ERS11936129 and ERS11936130-ERS11936138, respectively. The genomic data from the pools can be accessed with the ENA/NCBI IDs: ERS4275842 (GSL high pool), ERS4275843 (GSL low pool), ERS4275846 (SPC high pool) and ERS4275847 (SPC low pool). The genomic reads of the parents, Lorenz (P1) and Janetzkis Schlesischer (P2), can be found with the ENA/NCBI IDs ERS4368530 and ERS4368529, respectively. The applied scripts in this study are freely available on GitHub: https://GitHub.com/hschilbert/BnaMBS (DOI: 10.5281/zenodo.6578120).

## Acknowledgments

We are thankful to the whole NuGGET and RaPEQ team for the great support. Many thanks to Helene Schellenberg as well as to Willy Keller for excellent technical assistance. We thank Christian Möllers for providing the *B. napus* genotype SGDH14 and related phenotypic data.

## Conflicts of Interest

The authors declare no conflict of interest.

## Notes

https://doi.org/10.4119/unibi/2963492

https://zenodo.org/record/6578120

### Competing Interest Statement

The authors have declared no competing interest.

