## Supplementary figures and images for "Mapping-by-sequencing reveals genomic regions associated with seed quality parameters in *Brassica napus*"

### File S2

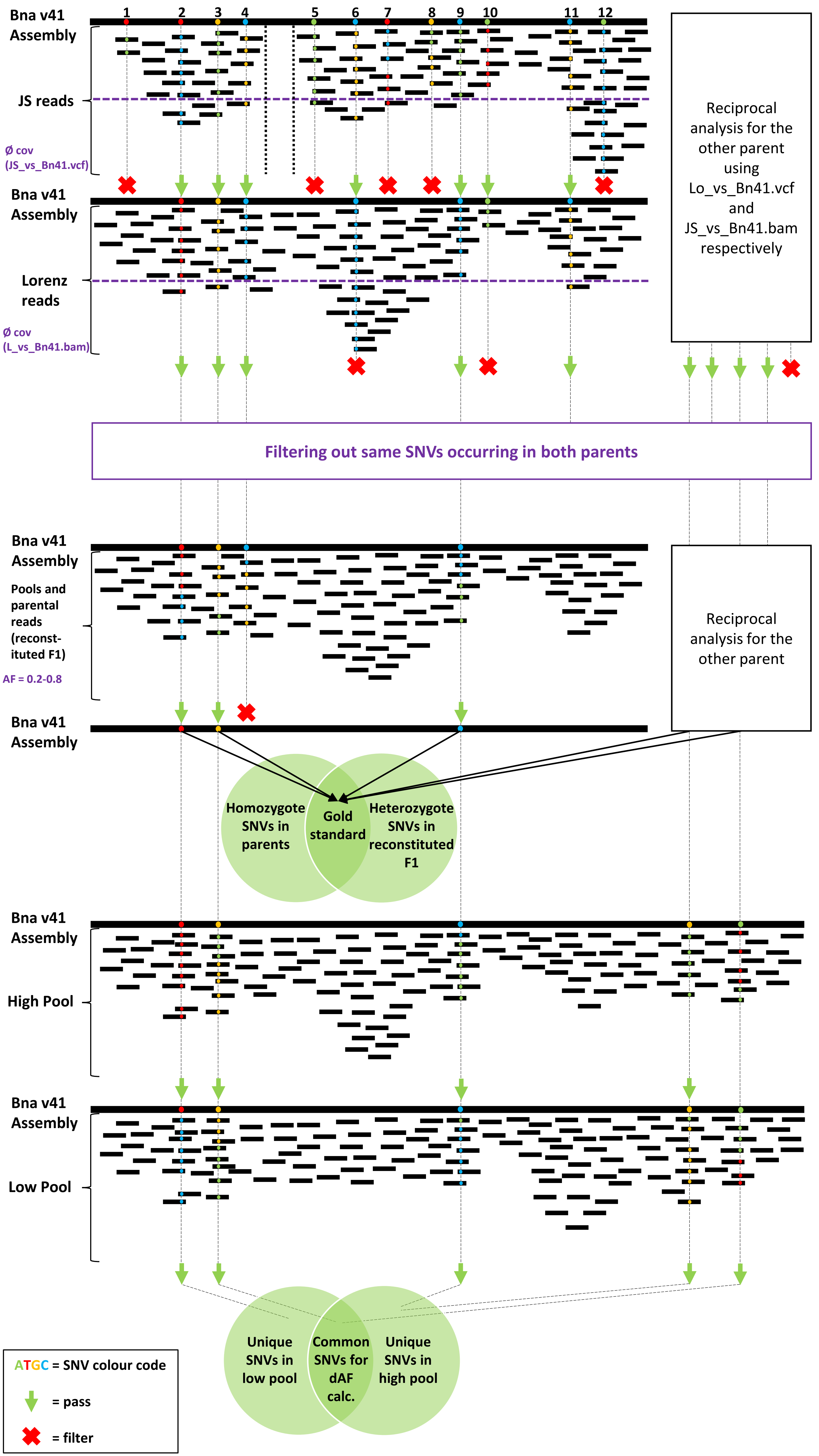

### File S34

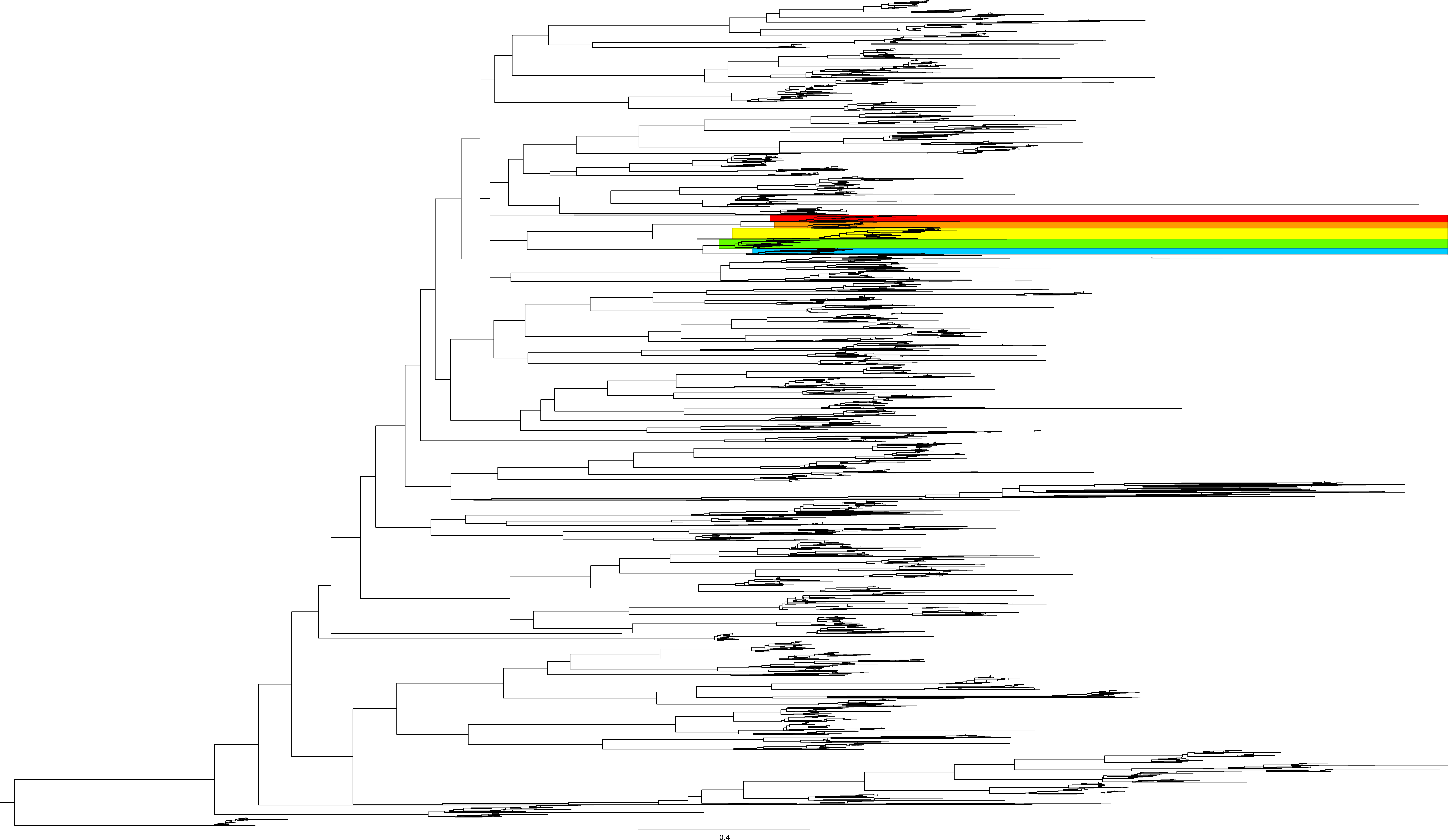

### File S35

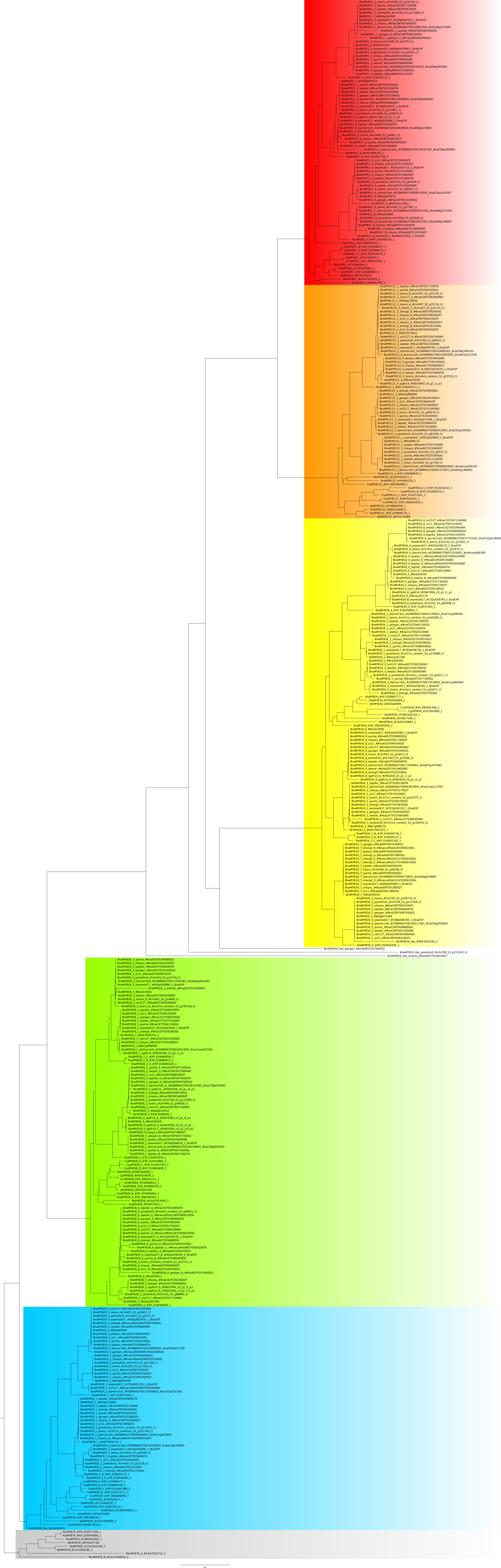

### File S39

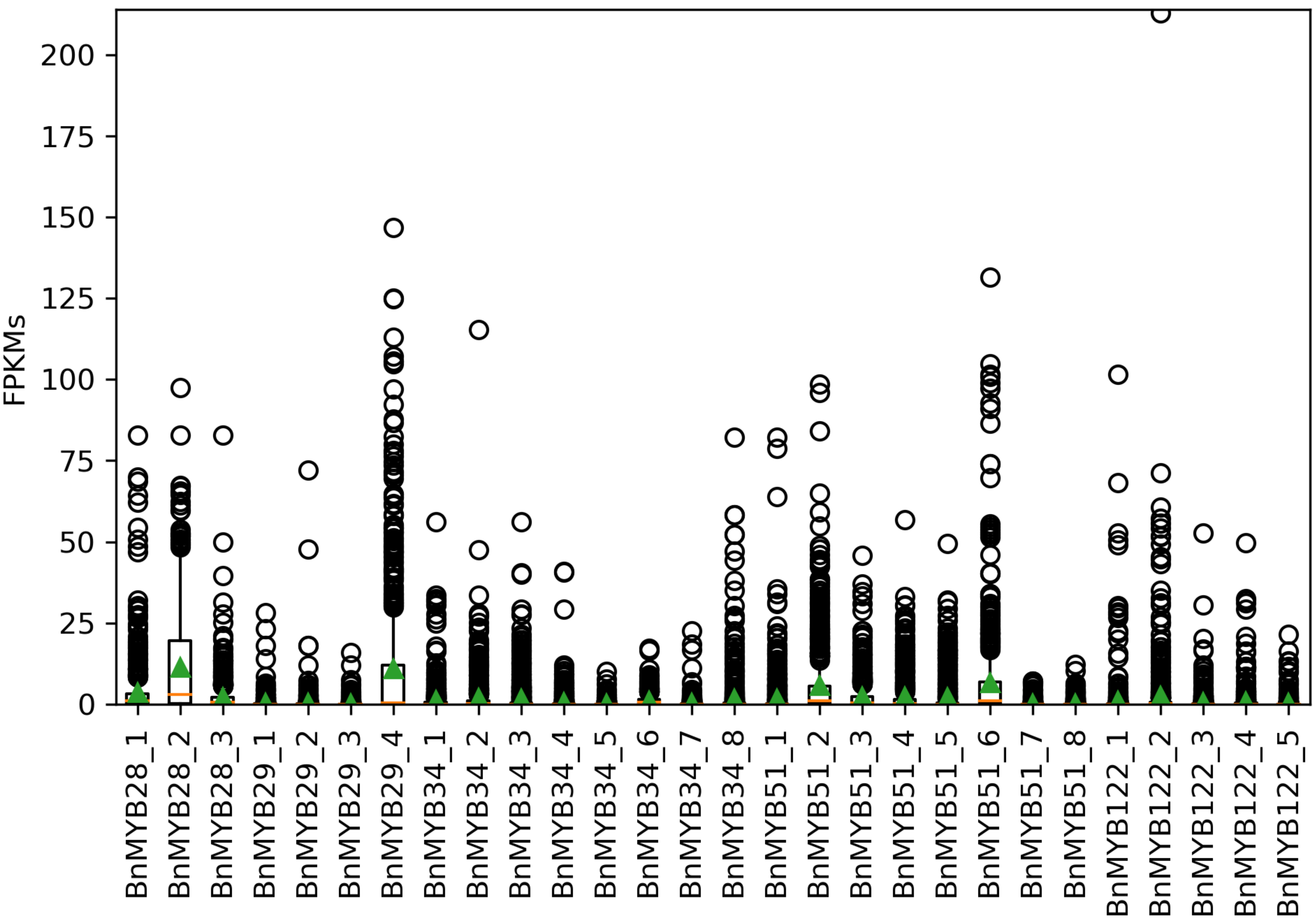

### File S40

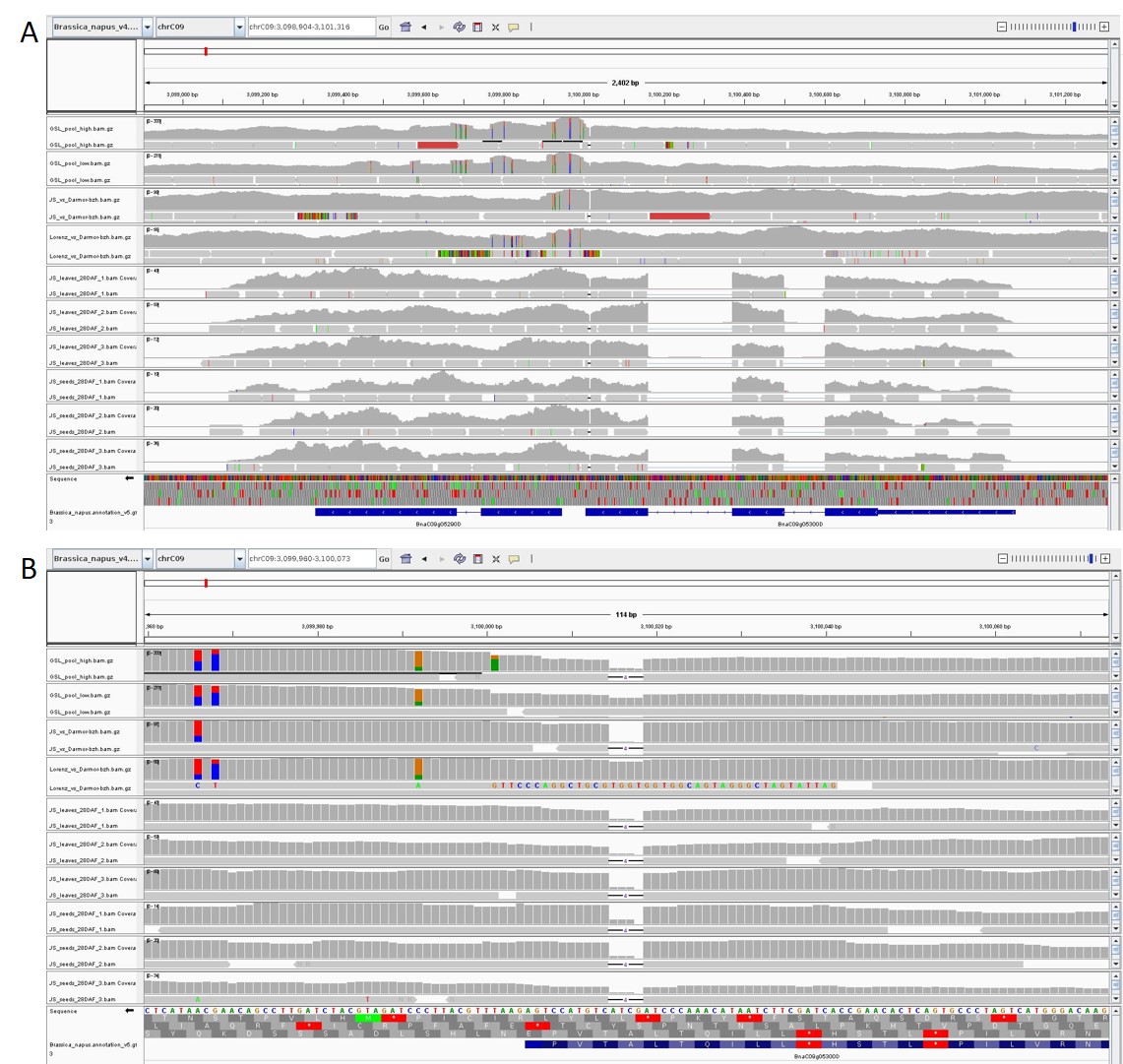

### File S42

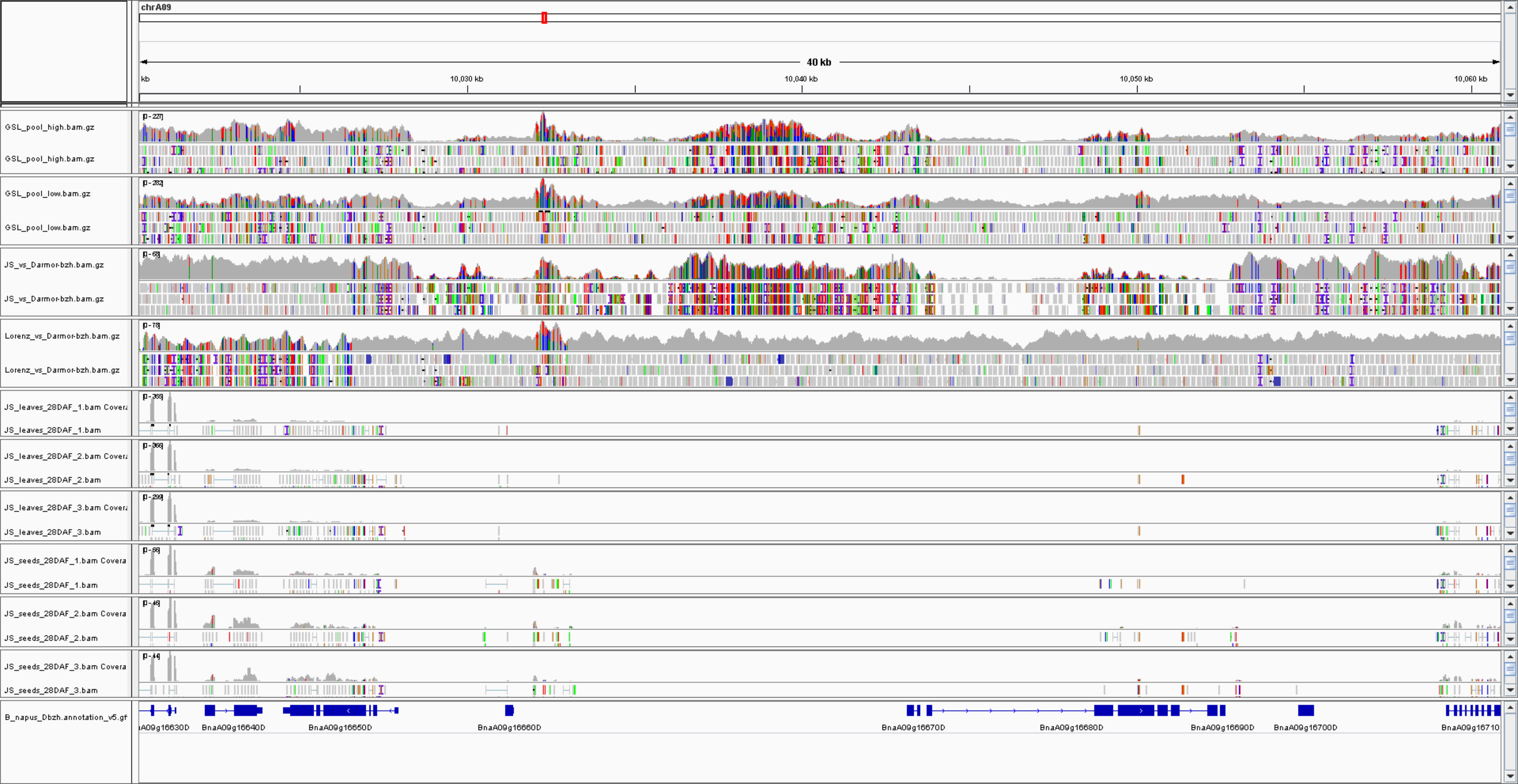
