## Supplementary material for "Mapping-by-sequencing reveals genomic regions associated with seed quality parameters in *Brassica napus*": File S11

**File S11: Genome-wide genomic intervals associated with seed GSL content.** The delta allele frequency between the high and the low pool, respectively, is displayed for all analysed positions in the *B. napus* genome sequence. A color code ranging from light blue over to magenta indicates the size of delta per chromosomal position. The normalized coverage is plotted in green and red corresponding to the low and high pool respectively. The density of normalized dARCs is showed in orange.

A01

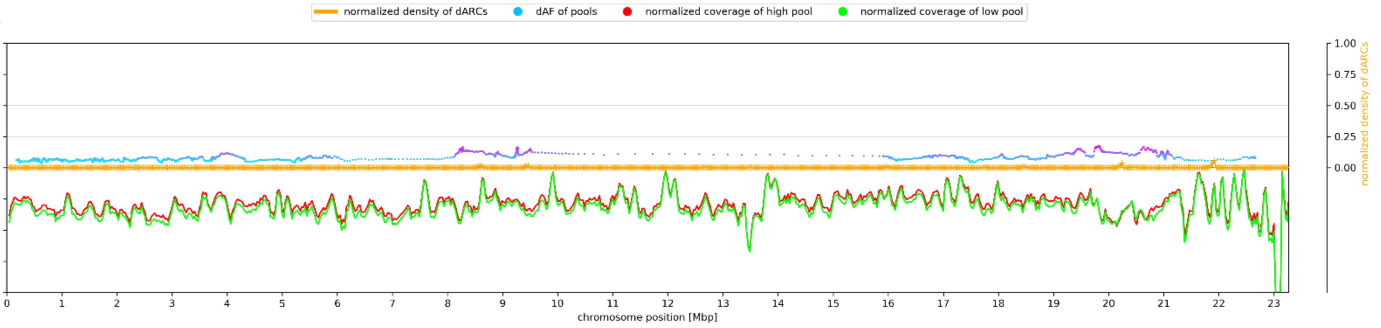


A02

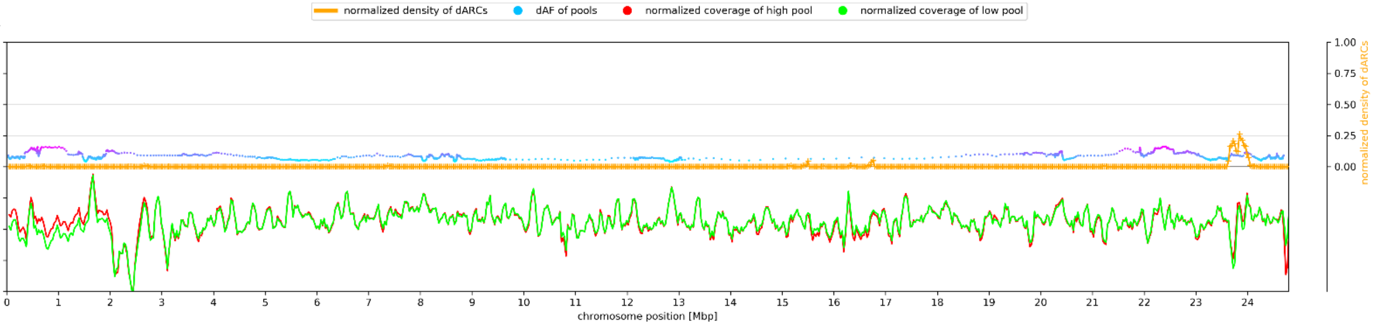


A03

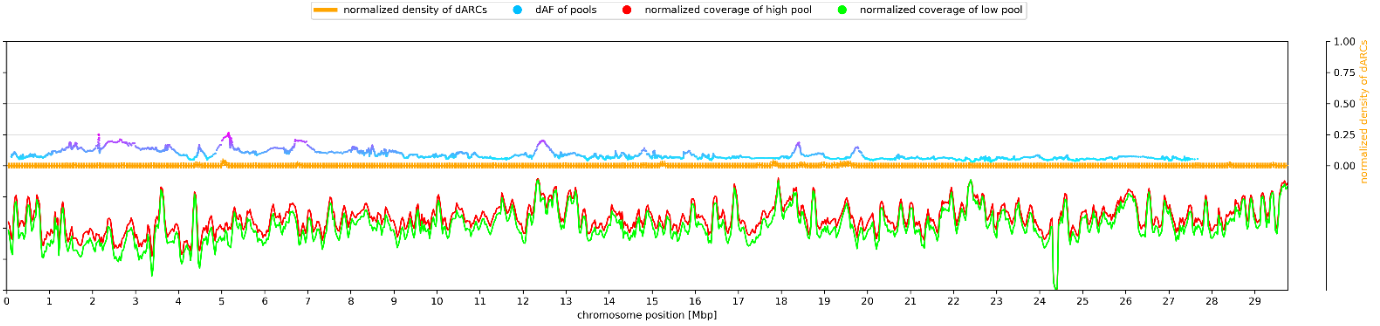


A04

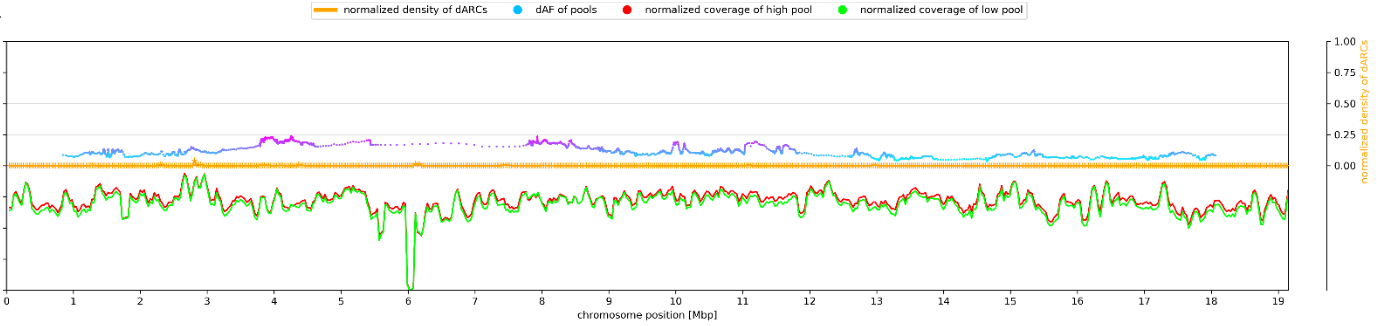


A05

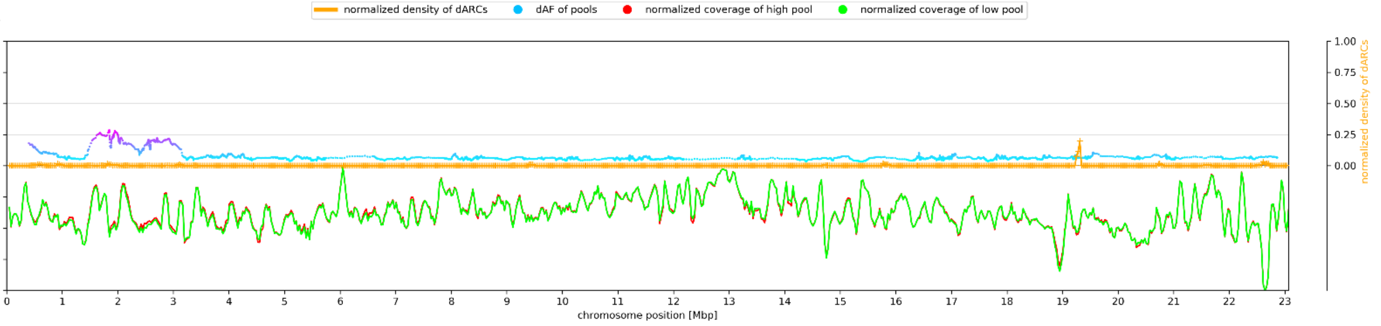


A06

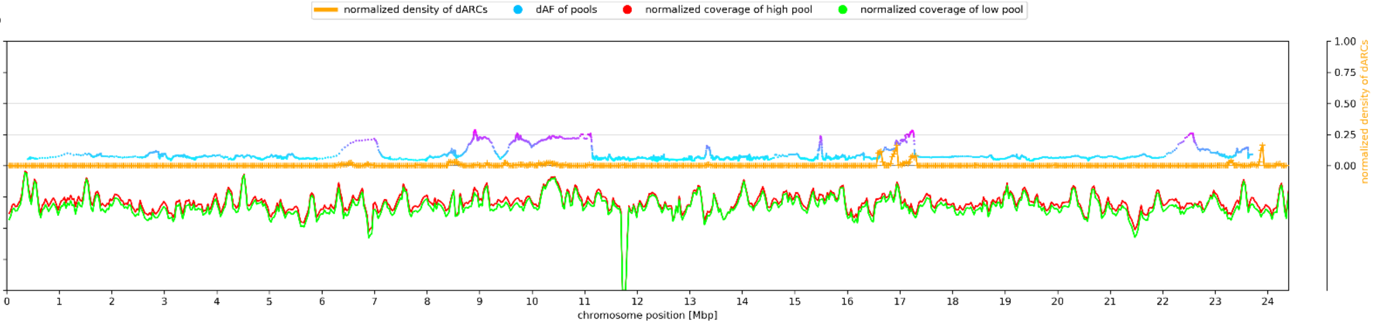

A07

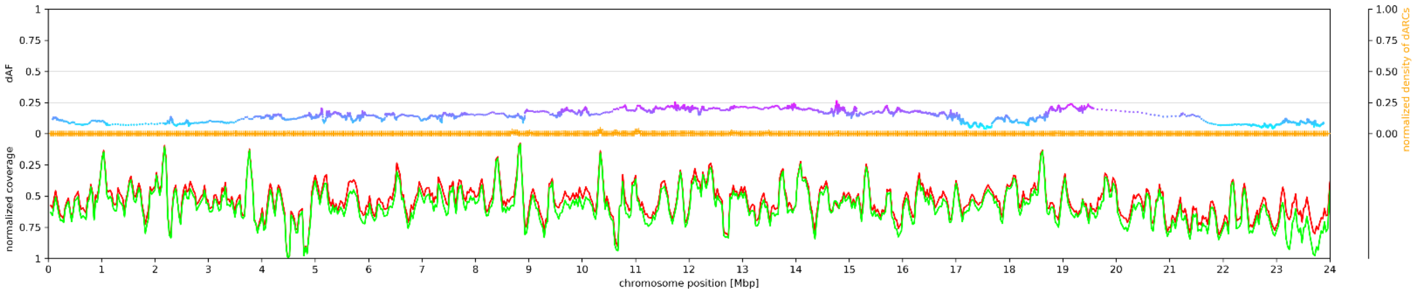


A08

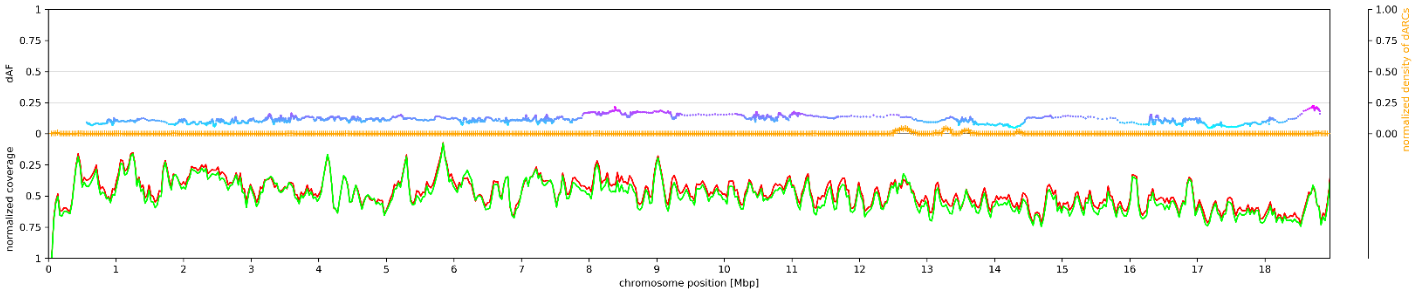


A09

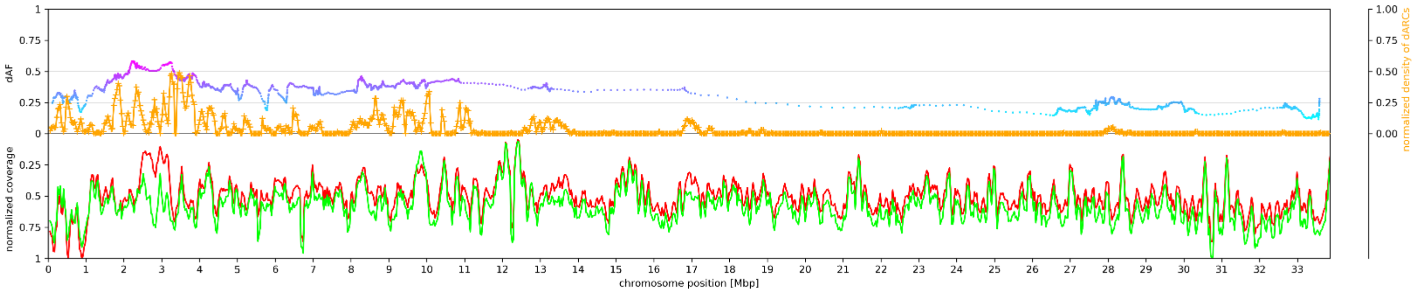


A10

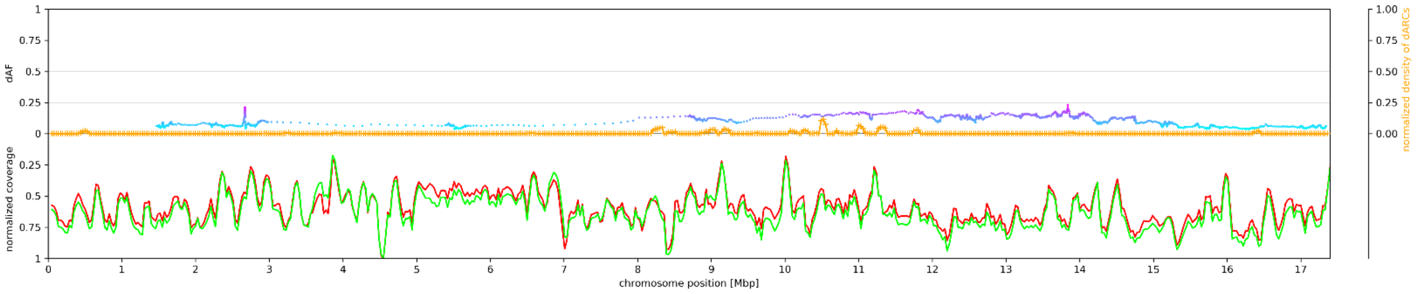


C01

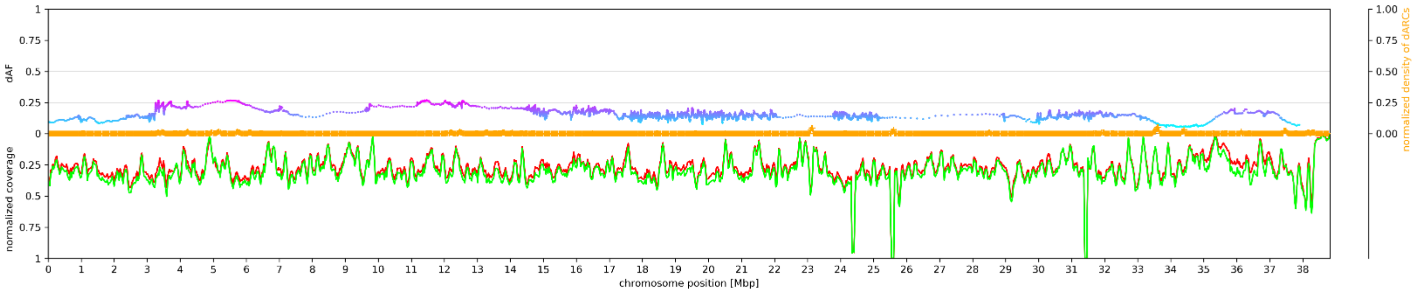


C02

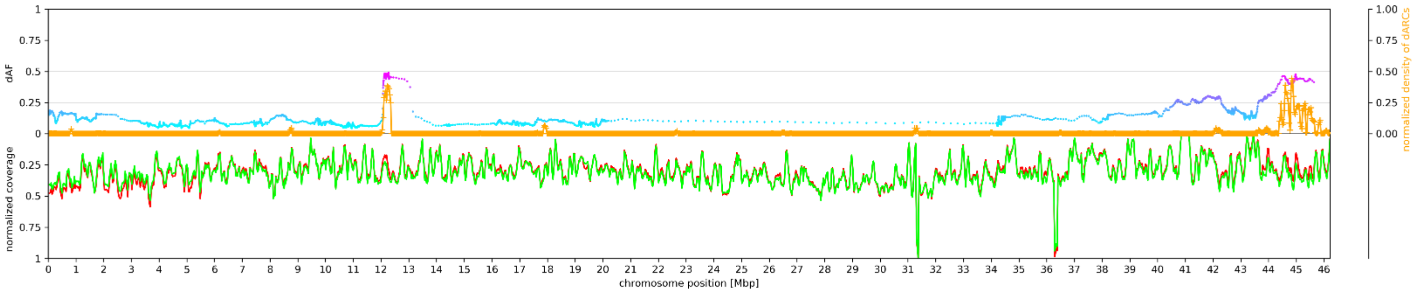


C03

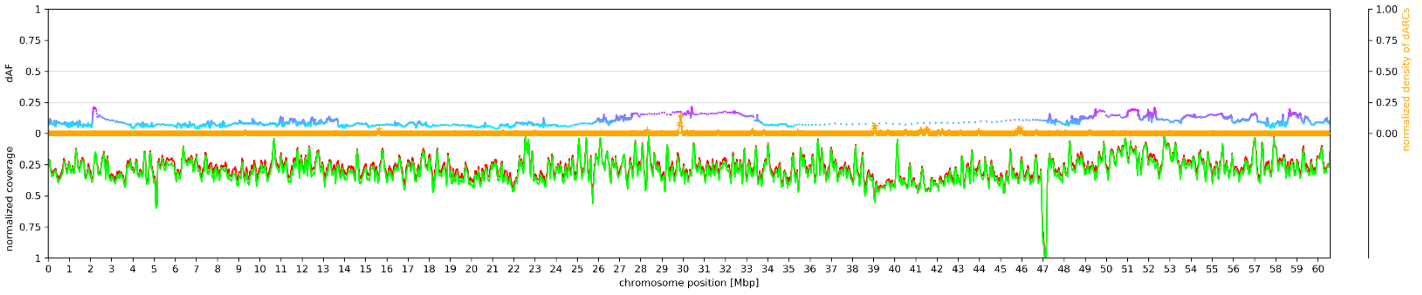

C04

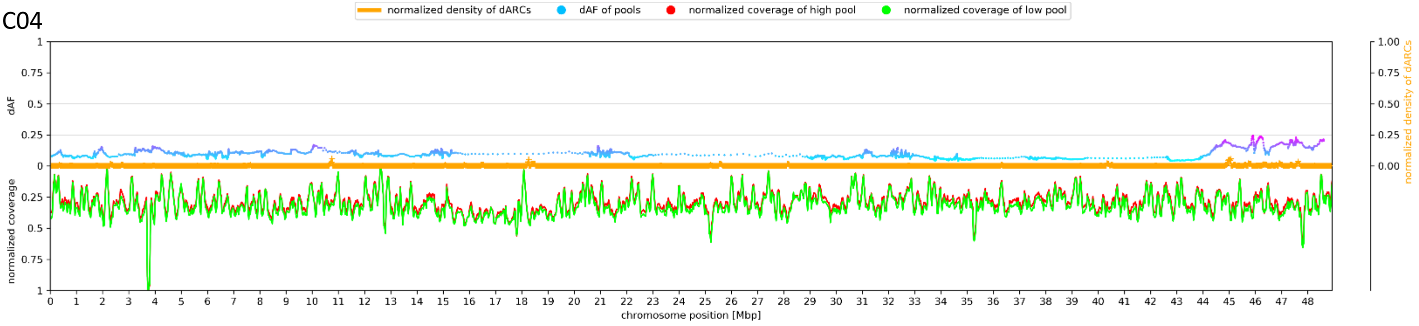


C05

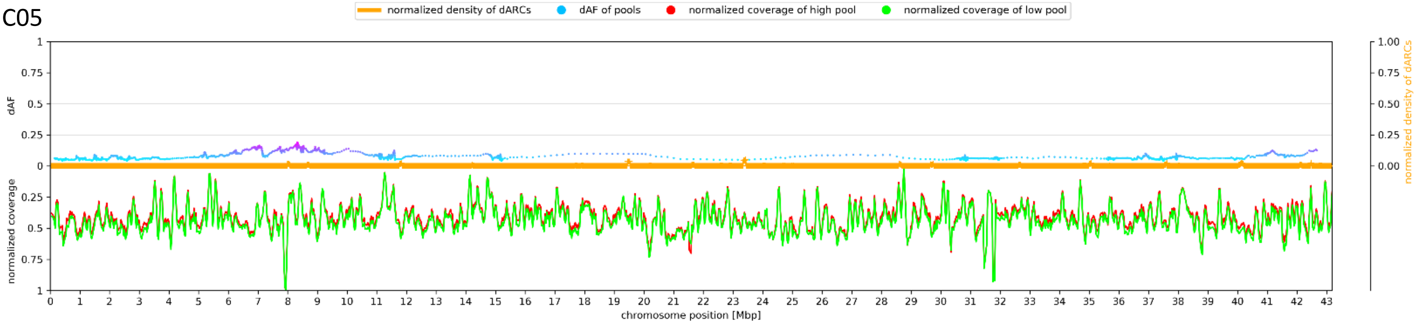


C06

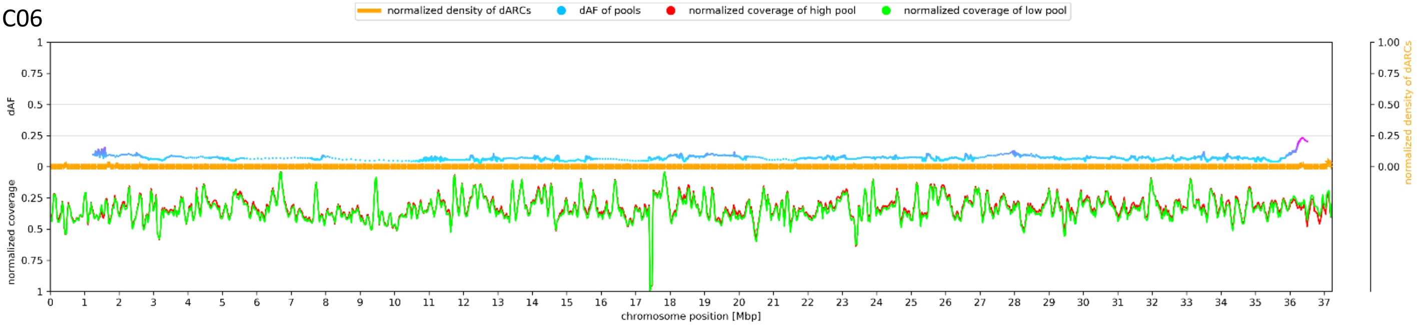


C07

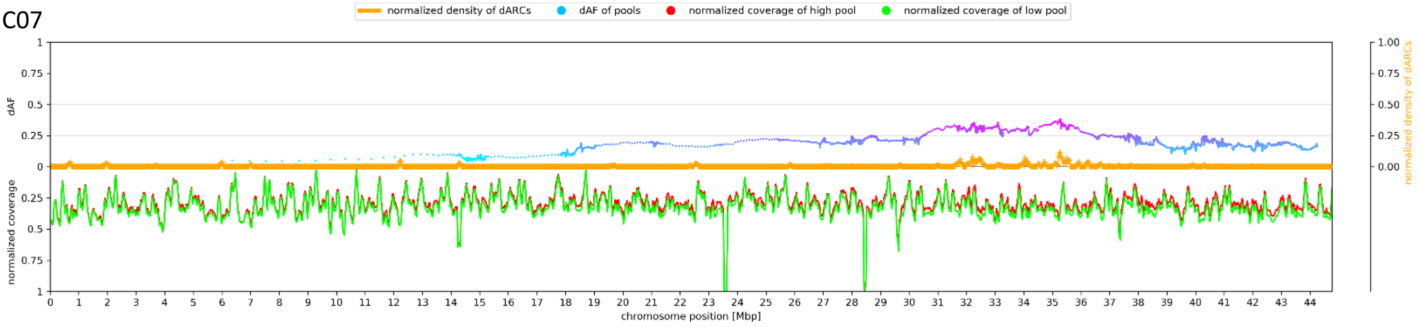


C08

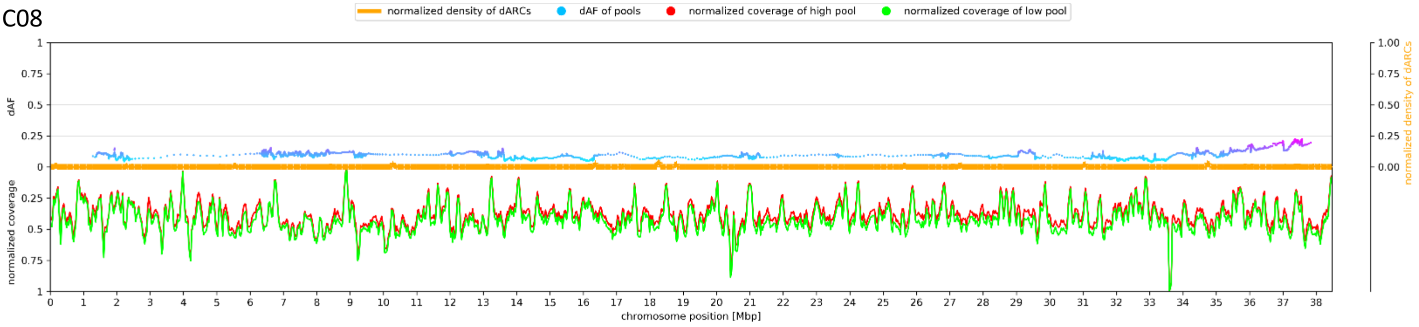


C09

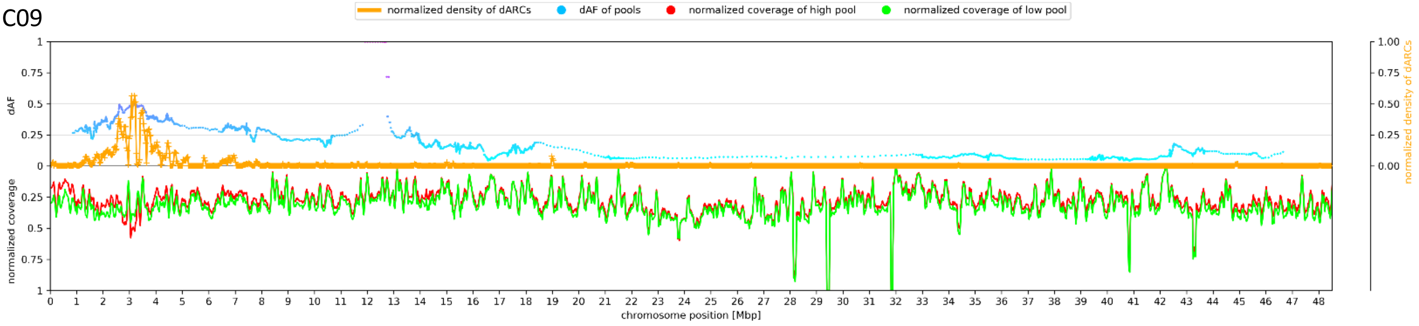
