## Supplementary material for "Mapping-by-sequencing reveals genomic regions associated with seed quality parameters in *Brassica napus*": File S15

### MYB28

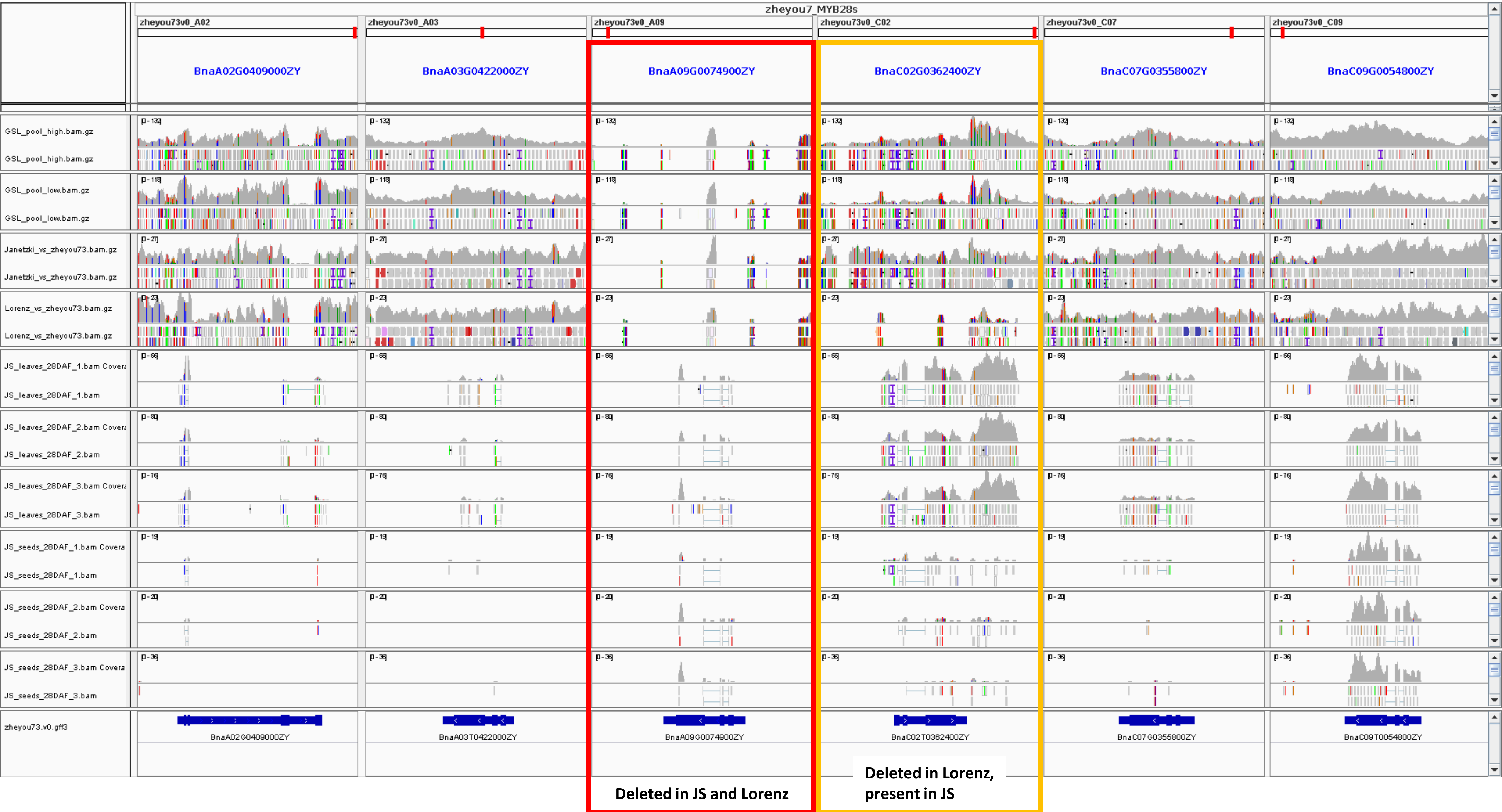
zheyou7_MYB28s
zheyou73v0_A02
zheyou73v0_A03
zheyou73v0_A09
zheyou73v0_C02
zheyou73v0_C07
zheyou73v0_C09
BnaA02G0409000ZY
BnaA03G0422000ZY
BnaA09G0074900ZY
BnaC02G0362400ZY
BnaC07G0355800ZY
BnaC09G0054800ZY
GSL_pool_high.bam.gz
GSL_pool_low.bam.gz
Janetzki_vs_zheyou73.bam.gz
Lorenz_vs_zheyou73.bam.gz
JS_leaves_28DAF_1.bam
JS_leaves_28DAF_2.bam
JS_leaves_28DAF_3.bam
JS_seeds_28DAF_1.bam
JS_seeds_28DAF_2.bam
JS_seeds_28DAF_3.bam
zheyou73.v0.gff3
BnaA02G0409000ZY
BnaA03T0422000ZY
BnaA09G0074900ZY
BnaC02T0362400ZY
BnaC07G0355800ZY
BnaC09T0054800ZY
Deleted in JS and Lorenz
Deleted in Lorenz, present in JS

### MYB28

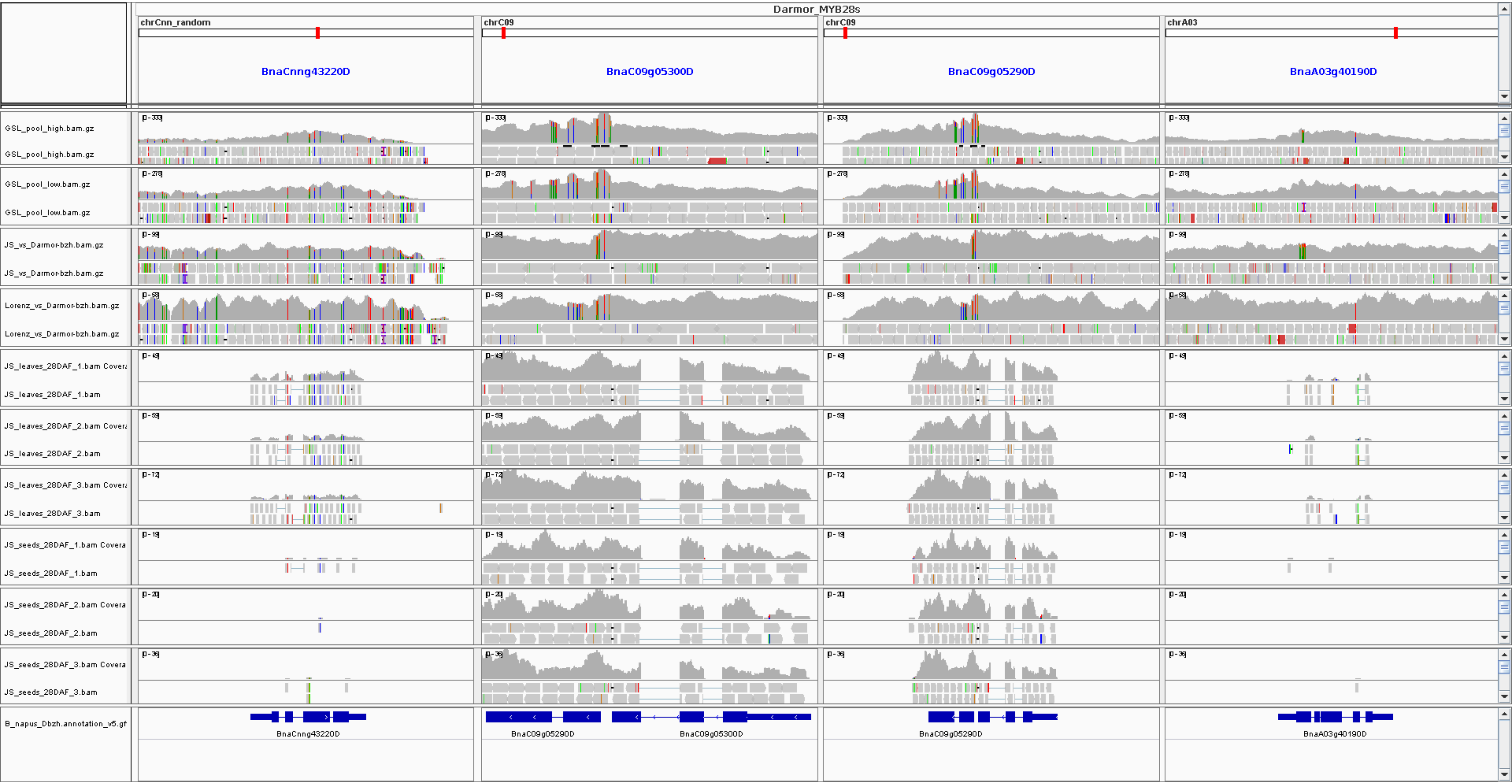

Darmor_MYB28s
chrCnn_random
chrC09
chrC09
chrA03
BnaCnng43220D
BnaC09g05300D
BnaC09g05290D
BnaA03g40190D
GSL_pool_high.bam.gz
GSL_pool_high.bam.gz
GSL_pool_low.bam.gz
GSL_pool_low.bam.gz
JS_vs_Darmor-bzh.bam.gz
JS_vs_Darmor-bzh.bam.gz
Lorenz_vs_Darmor-bzh.bam.gz
Lorenz_vs_Darmor-bzh.bam.gz
JS_leaves_28DAF_1.bam Covera
JS_leaves_28DAF_1.bam
JS_leaves_28DAF_2.bam Covera
JS_leaves_28DAF_2.bam
JS_leaves_28DAF_3.bam Covera
JS_leaves_28DAF_3.bam
JS_seeds_28DAF_1.bam Covera
JS_seeds_28DAF_1.bam
JS_seeds_28DAF_2.bam Covera
JS_seeds_28DAF_2.bam
JS_seeds_28DAF_3.bam Covera
JS_seeds_28DAF_3.bam
B_napus_Dbzh.annotation_v5.gf
BnaCnng43220D
BnaC09g05290D
BnaC09g05300D
BnaC09g05290D
BnaA03g40190D

### MYB28

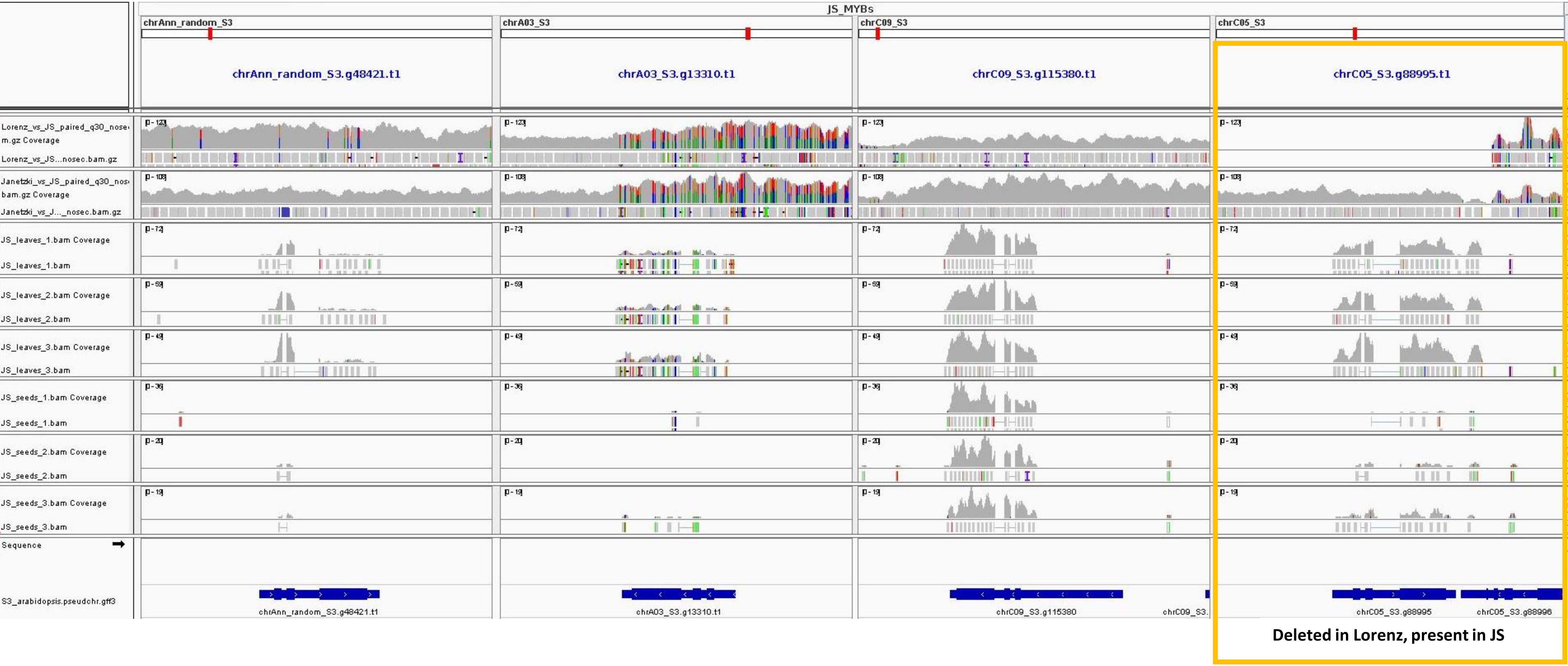

JS_MYBs
chrAnn_random_S3
chrA03_S3
chrC09_S3
chrC05_S3
chrAnn_random_S3.g48421.t1
chrA03_S3.g13310.t1
chrC09_S3.g115380.t1
chrC05_S3.g88995.t1
Lorenz_vs_JS_paired_q30_nosem.gz Coverage
Lorenz_vs_JS...nosec.bam.gz
Janetzki_vs_JS_paired_q30_nosbam.gz Coverage
Janetzki_vs_J..._nosec.bam.gz
JS_leaves_1.bam Coverage
JS_leaves_1.bam
JS_leaves_2.bam Coverage
JS_leaves_2.bam
JS_leaves_3.bam Coverage
JS_leaves_3.bam
JS_seeds_1.bam Coverage
JS_seeds_1.bam
JS_seeds_2.bam Coverage
JS_seeds_2.bam
JS_seeds_3.bam Coverage
JS_seeds_3.bam
Sequence
S3_arabidopsis.pseudchr.gff3
chrAnn_random_S3.g48421.t1
chrA03_S3.g13310.t1
chrC09_S3.g115380
chrC05_S3.g88995
chrC05_S3.g88996
Deleted in Lorenz, present in JS

### MYB28

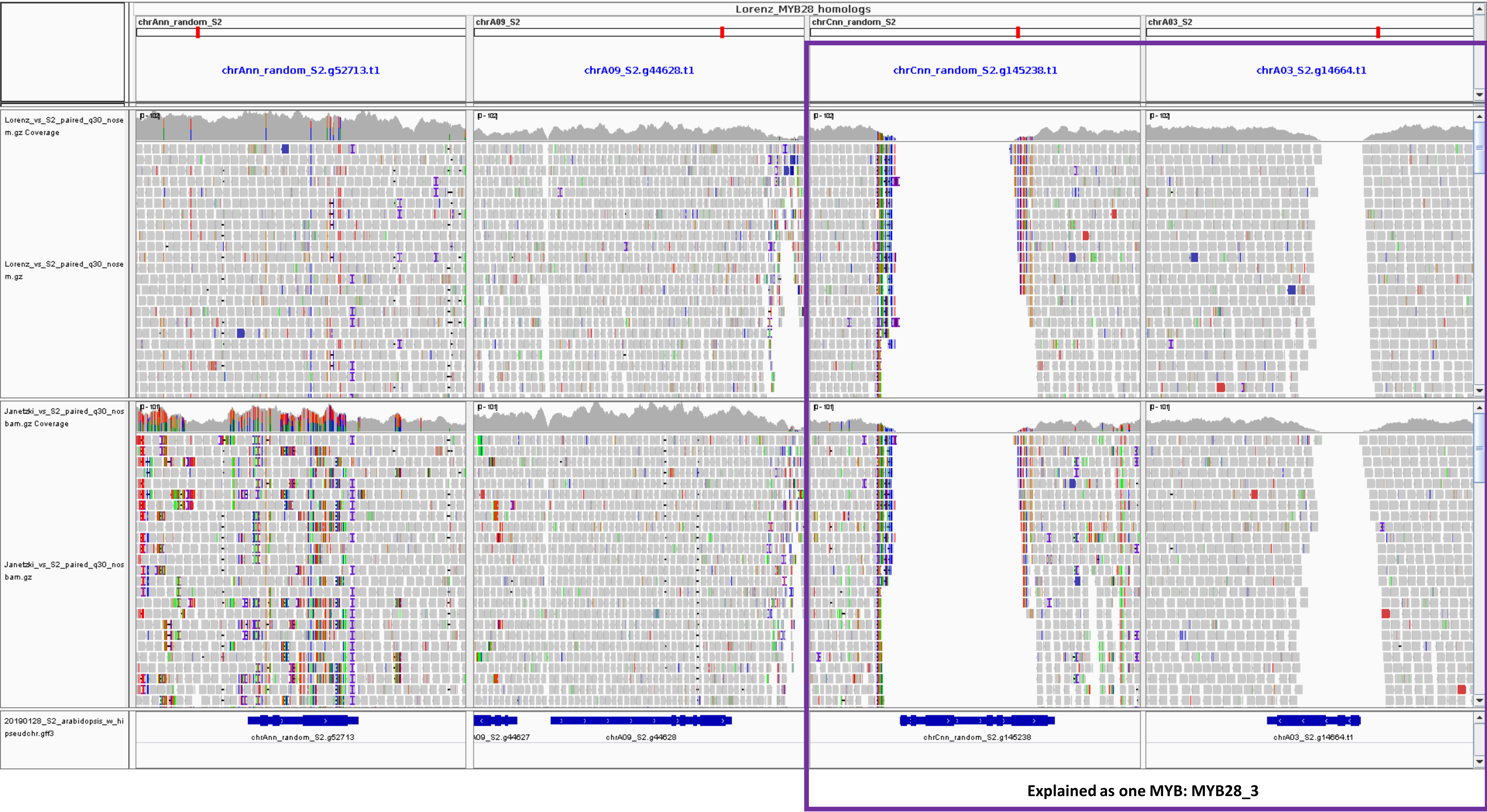
Lorenz_MYB28_homologs
chrAnn_random_S2
chrA09_S2
chrCnn_random_S2
chrA03_S2
chrAnn_random_S2.g52713.t1
chrA09_S2.g44628.t1
chrCnn_random_S2.g145238.t1
chrA03_S2.g14664.t1
Lorenz_vs_S2_paired_q30_nosem.gz Coverage
Lorenz_vs_S2_paired_q30_nosem.gz
Janetzki_vs_S2_paired_q30_nosbam.gz Coverage
Janetzki_vs_S2_paired_q30_nosbam.gz
20190128_S2_arabidopsis_w_hi pseudchr.gff3
chrAnn_random_S2.g52713
A09_S2.g44627
chrA09_S2.g44628
chrCnn_random_S2.g145238
chrA03_S2.g14664.t1
Explained as one MYB: MYB28_3

### MYB29

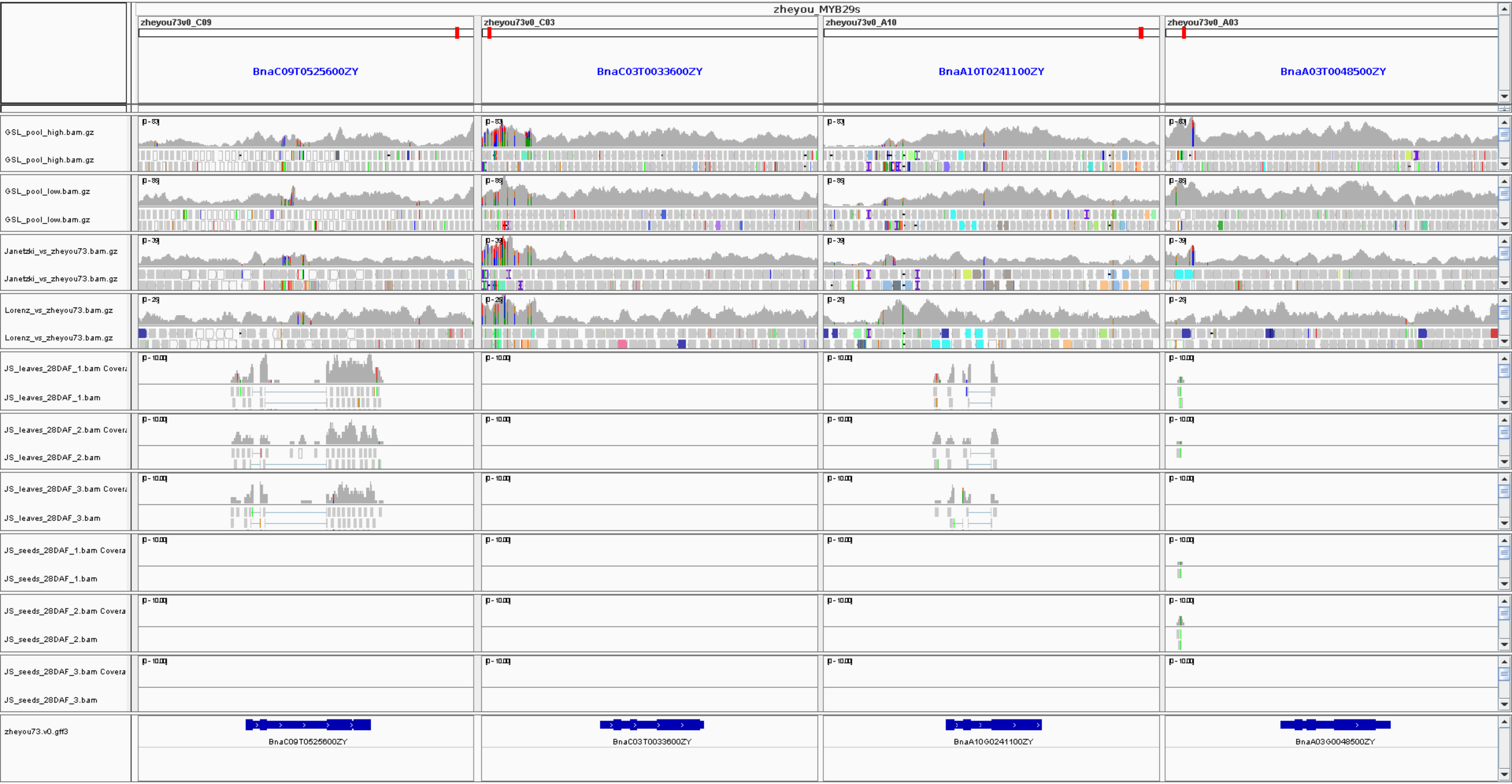

zheyou_MYB29s
zheyou73v0_C09
zheyou73v0_C03
zheyou73v0_A10
zheyou73v0_A03
BnaC09T0525600ZY
BnaC03T0033600ZY
BnaA10T0241100ZY
BnaA03T0048500ZY
GSL_pool_high.bam.gz
GSL_pool_low.bam.gz
Janetzki_vs_zheyou73.bam.gz
Lorenz_vs_zheyou73.bam.gz
JS_leaves_28DAF_1.bam
JS_leaves_28DAF_2.bam
JS_leaves_28DAF_3.bam
JS_seeds_28DAF_1.bam
JS_seeds_28DAF_2.bam
JS_seeds_28DAF_3.bam
zheyou73.v0.gff3
BnaC09T0525600ZY
BnaC03T0033600ZY
BnaA10G0241100ZY
BnaA03G0048500ZY

### MYB29

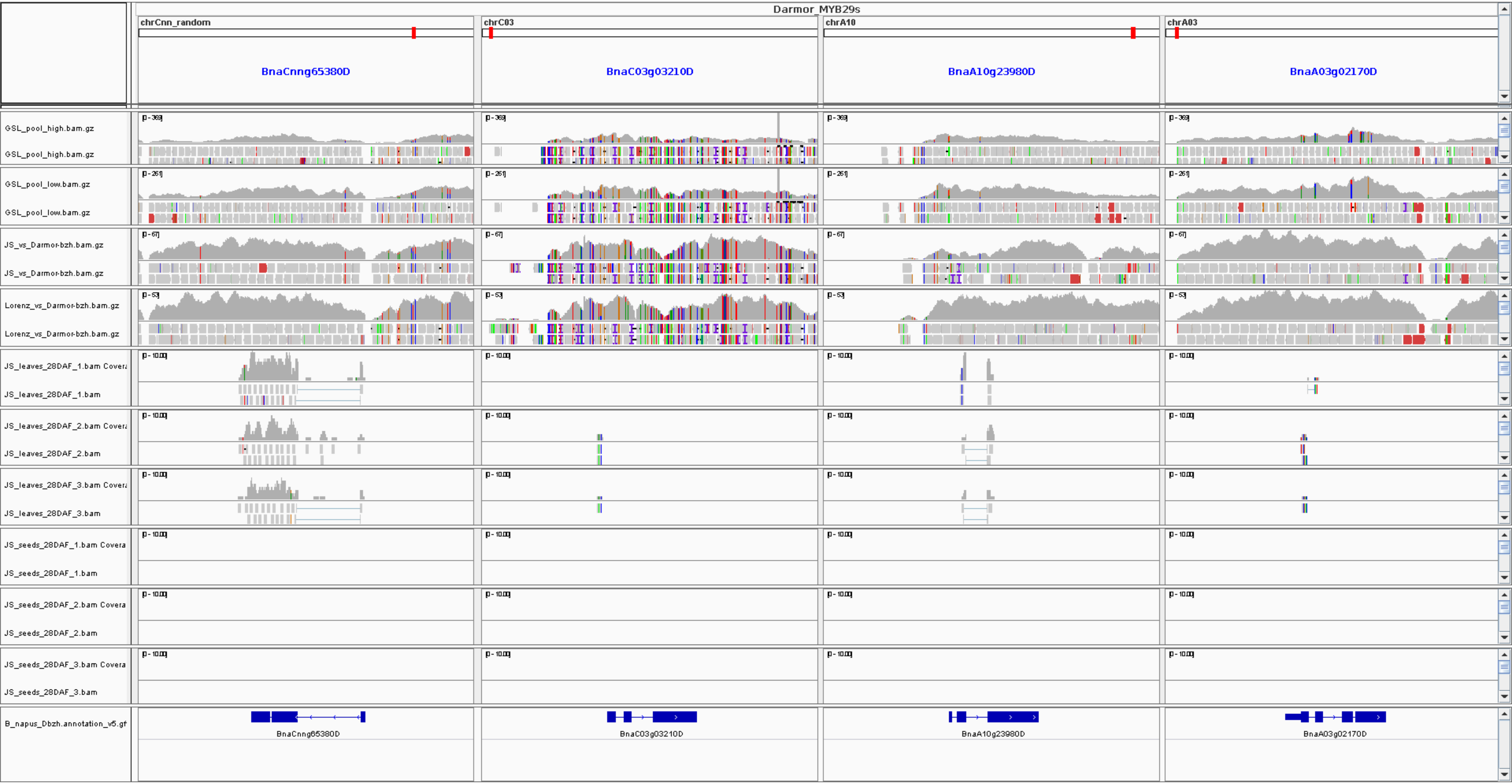

Darmor_MYB29s
chrCnn_random
chrC03
chrA10
chrA03
BnaCnng65380D
BnaC03g03210D
BnaA10g23980D
BnaA03g02170D
GSL_pool_high.bam.gz
GSL_pool_high.bam.gz
GSL_pool_low.bam.gz
GSL_pool_low.bam.gz
JS_vs_Darmor-bzh.bam.gz
JS_vs_Darmor-bzh.bam.gz
Lorenz_vs_Darmor-bzh.bam.gz
Lorenz_vs_Darmor-bzh.bam.gz
JS_leaves_28DAF_1.bam Covera
JS_leaves_28DAF_1.bam
JS_leaves_28DAF_2.bam Covera
JS_leaves_28DAF_2.bam
JS_leaves_28DAF_3.bam Covera
JS_leaves_28DAF_3.bam
JS_seeds_28DAF_1.bam Covera
JS_seeds_28DAF_1.bam
JS_seeds_28DAF_2.bam Covera
JS_seeds_28DAF_2.bam
JS_seeds_28DAF_3.bam Covera
JS_seeds_28DAF_3.bam
B_napus_Dbzh.annotation_v5.gf
BnaCnng65380D
BnaC03g03210D
BnaA10g23980D
BnaA03g02170D

### MYB34_1

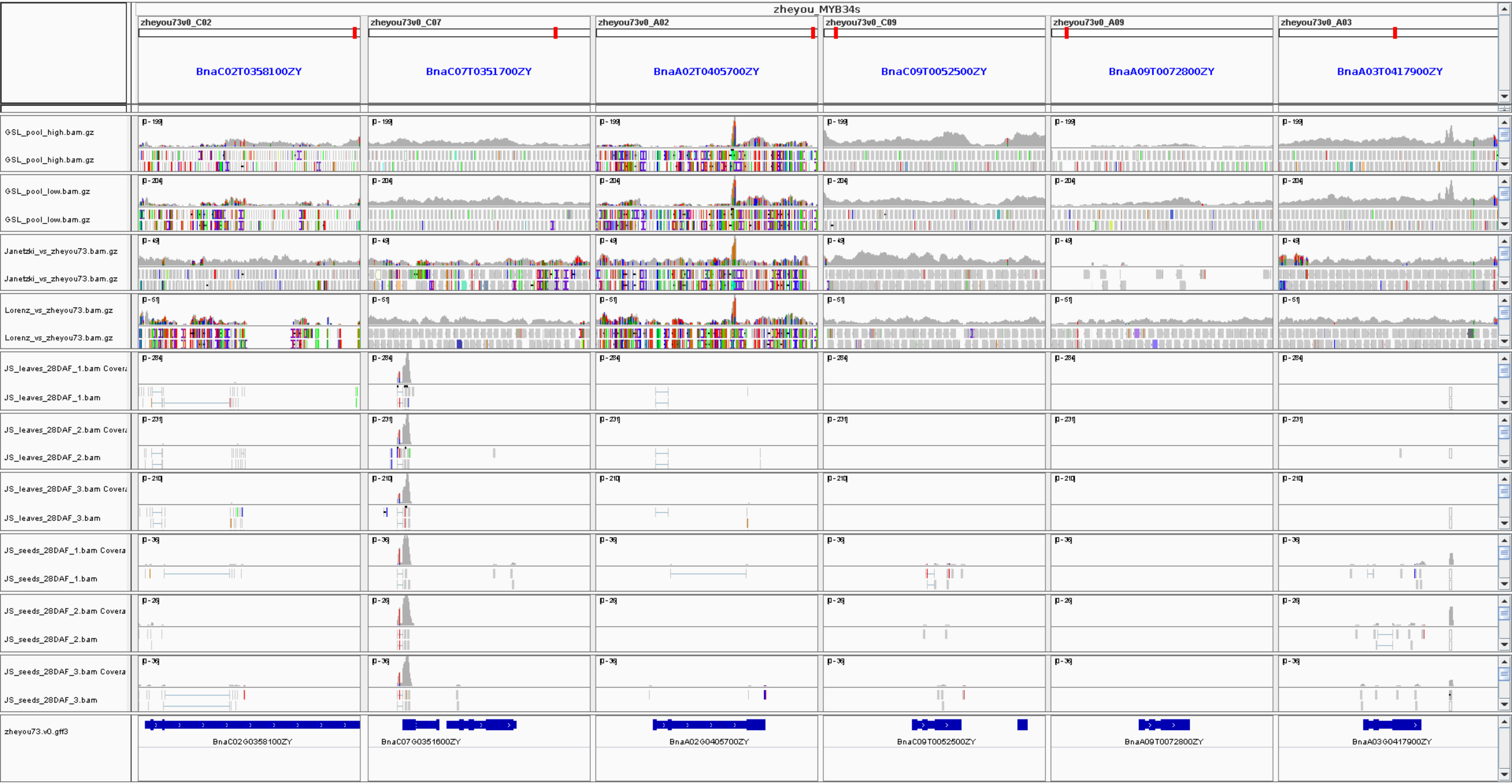
zheyou_MYB34s
zheyou73v0_C02
zheyou73v0_C07
zheyou73v0_A02
zheyou73v0_C09
zheyou73v0_A09
zheyou73v0_A03
BnaC02T0358100ZY
BnaC07T0351700ZY
BnaA02T0405700ZY
BnaC09T0052500ZY
BnaA09T0072800ZY
BnaA03T0417900ZY
GSL_pool_high.bam.gz
GSL_pool_low.bam.gz
Janetzki_vs_zheyou73.bam.gz
Lorenz_vs_zheyou73.bam.gz
JS_leaves_28DAF_1.bam
JS_leaves_28DAF_2.bam
JS_leaves_28DAF_3.bam
JS_seeds_28DAF_1.bam
JS_seeds_28DAF_2.bam
JS_seeds_28DAF_3.bam
zheyou73.v0.gff3
BnaC02G0358100ZY
BnaC07G0351600ZY
BnaA02G0405700ZY
BnaC09T0052500ZY
BnaA09T0072800ZY
BnaA03G0417900ZY

### MYB34_2

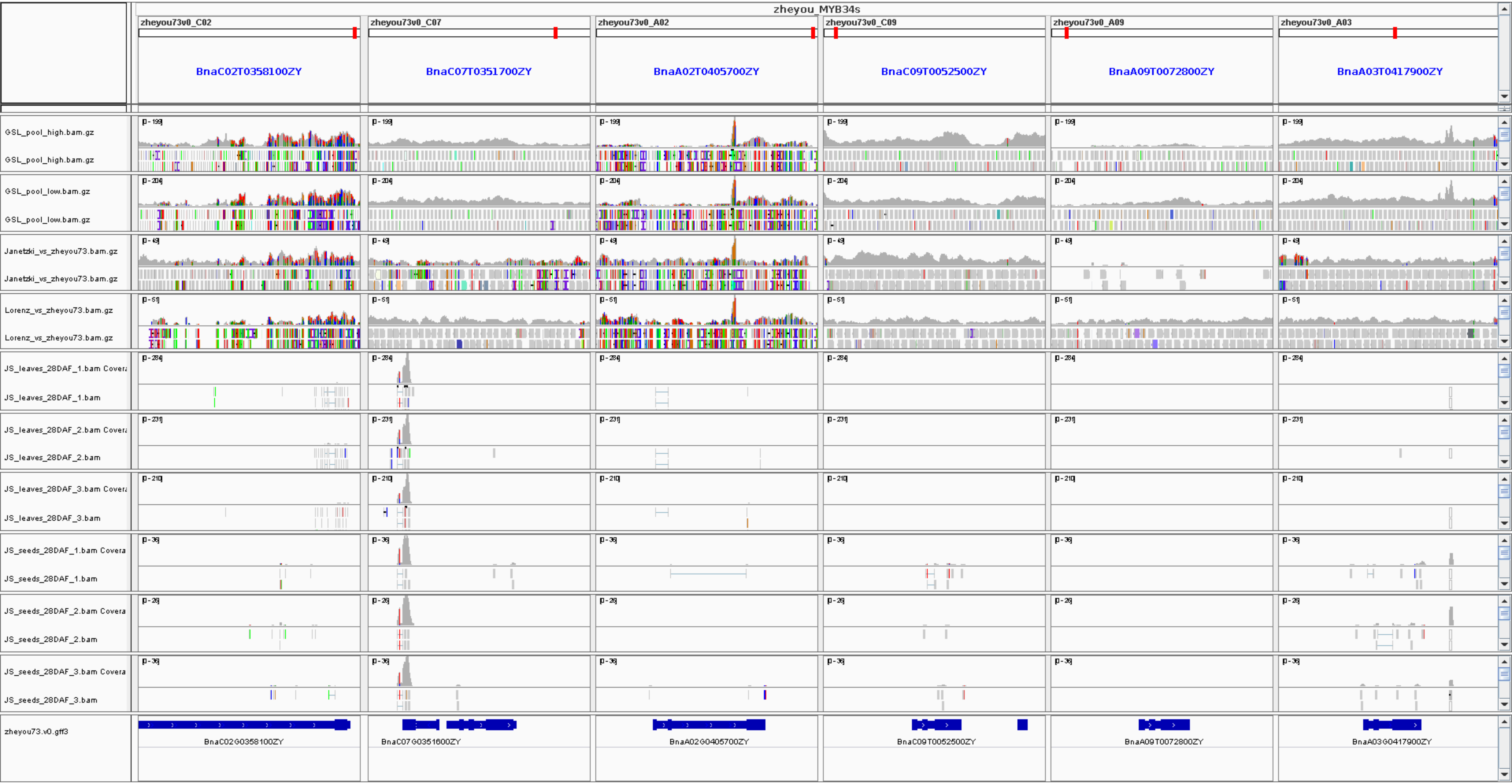

zheyou_MYB34s
zheyou73v0_C02
zheyou73v0_C07
zheyou73v0_A02
zheyou73v0_C09
zheyou73v0_A09
zheyou73v0_A03
BnaC02T0358100ZY
BnaC07T0351700ZY
BnaA02T0405700ZY
BnaC09T0052500ZY
BnaA09T0072800ZY
BnaA03T0417900ZY
GSL_pool_high.bam.gz
GSL_pool_low.bam.gz
Janetzki_vs_zheyou73.bam.gz
Lorenz_vs_zheyou73.bam.gz
JS_leaves_28DAF_1.bam
JS_leaves_28DAF_2.bam
JS_leaves_28DAF_3.bam
JS_seeds_28DAF_1.bam
JS_seeds_28DAF_2.bam
JS_seeds_28DAF_3.bam
zheyou73.v0.gff3
BnaC02G0358100ZY
BnaC07G0351600ZY
BnaA02G0405700ZY
BnaC09T0052500ZY
BnaA09T0072800ZY
BnaA03G0417900ZY

### MYB34

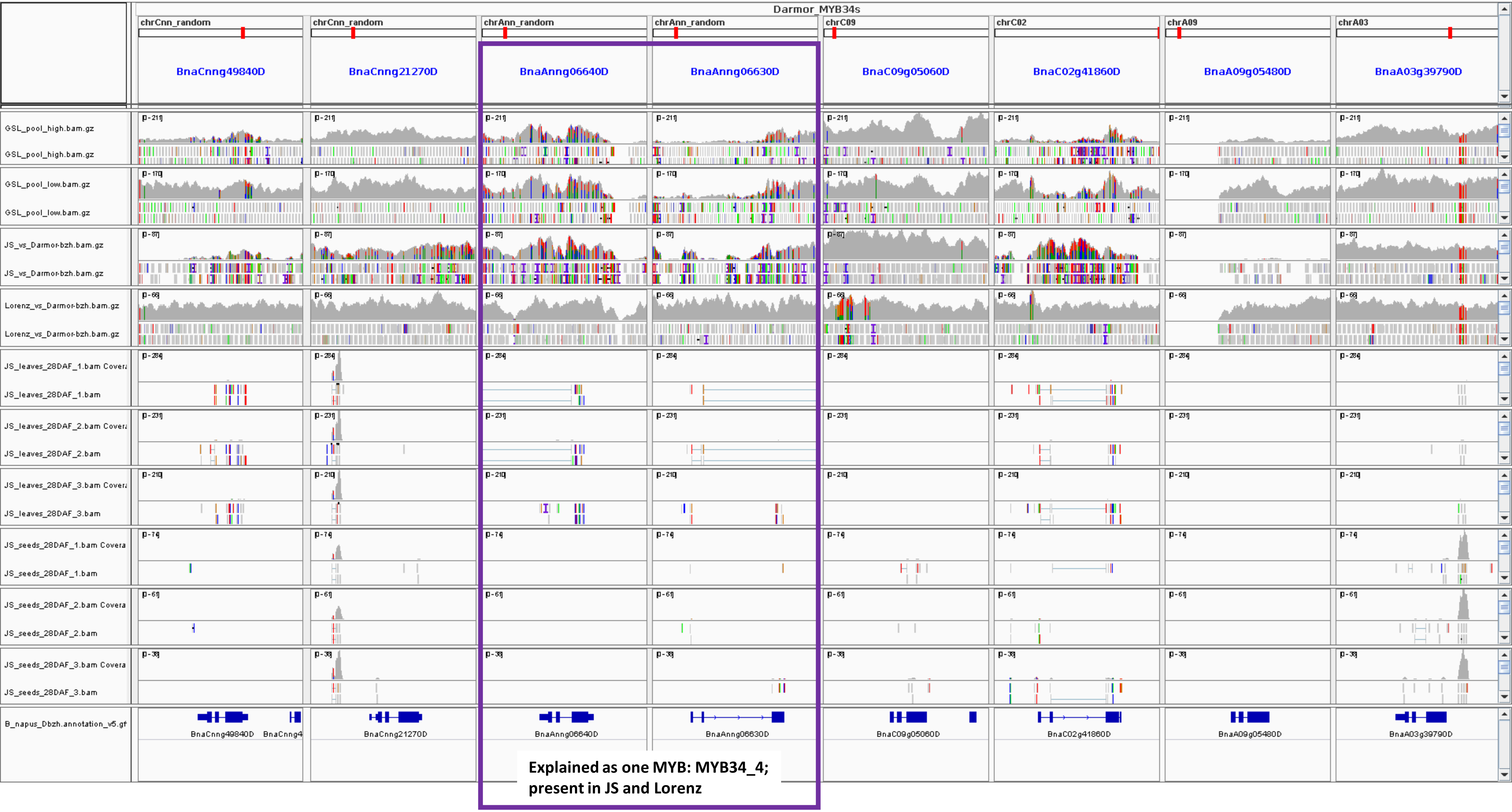
Darmor_MYB34s
chrCnn_random
chrCnn_random
chrAnn_random
chrAnn_random
chrC09
chrC02
chrA09
chrA03
BnaCnng49840D
BnaCnng21270D
BnaAnng06640D
BnaAnng06630D
BnaC09g05060D
BnaC02g41860D
BnaA09g05480D
BnaA03g39790D
GSL_pool_high.bam.gz
GSL_pool_high.bam.gz
GSL_pool_low.bam.gz
GSL_pool_low.bam.gz
JS_vs_Darmor-bzh.bam.gz
JS_vs_Darmor-bzh.bam.gz
Lorenz_vs_Darmor-bzh.bam.gz
Lorenz_vs_Darmor-bzh.bam.gz
JS_leaves_28DAF_1.bam Covera
JS_leaves_28DAF_1.bam
JS_leaves_28DAF_2.bam Covera
JS_leaves_28DAF_2.bam
JS_leaves_28DAF_3.bam Covera
JS_leaves_28DAF_3.bam
JS_seeds_28DAF_1.bam Covera
JS_seeds_28DAF_1.bam
JS_seeds_28DAF_2.bam Covera
JS_seeds_28DAF_2.bam
JS_seeds_28DAF_3.bam Covera
JS_seeds_28DAF_3.bam
B_napus_Dbzh.annotation_v5.gf
BnaCnng49840D
BnaCnng4
BnaCnng21270D
BnaAnng06640D
BnaAnng06630D
BnaC09g05060D
BnaC02g41860D
BnaA09g05480D
BnaA03g39790D
Explained as one MYB: MYB34_4;
present in JS and Lorenz

### MYB34_3 and MYB34_4

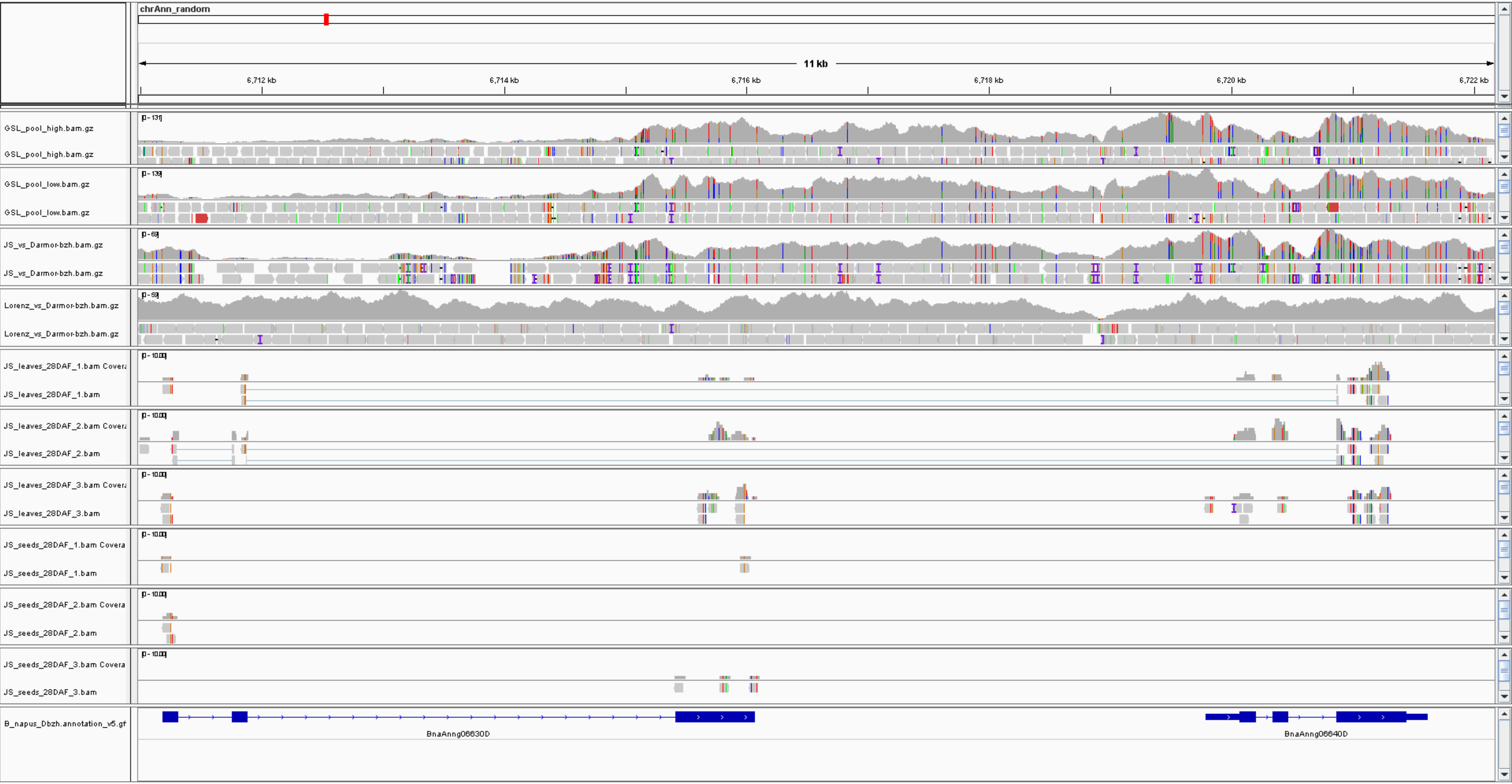


**Shows that the two annotated gene models in Darmor-bzh BnaAnng06630D and BnaAnng06640D likely explain one MYB, as supported by the RNA-Seq data**

### MYB51

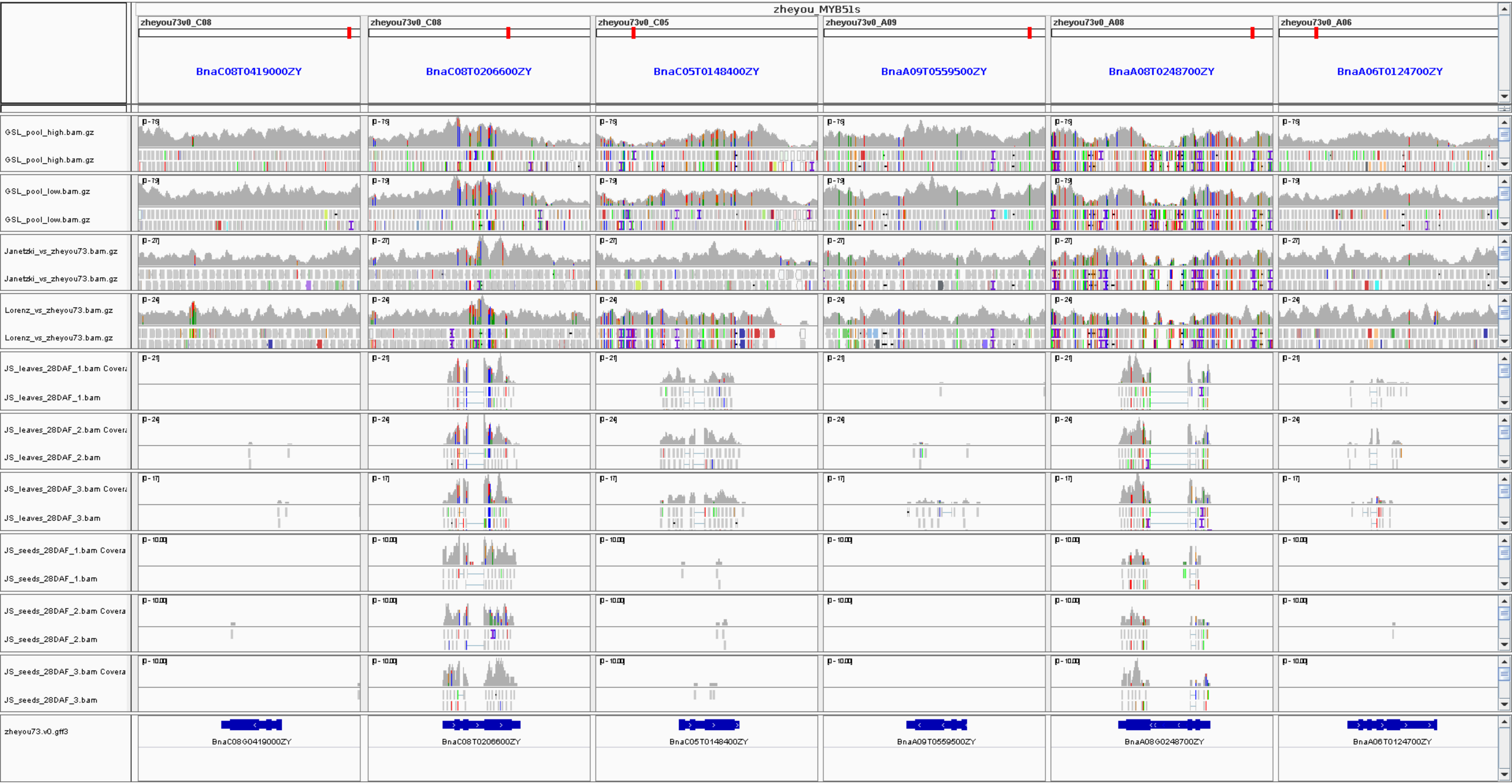
zheyou_MYB51s
zheyou73v0_C08
zheyou73v0_C08
zheyou73v0_C05
zheyou73v0_A09
zheyou73v0_A08
zheyou73v0_A06
BnaC08T0419000ZY
BnaC08T0206600ZY
BnaC05T0148400ZY
BnaA09T0559500ZY
BnaA08T0248700ZY
BnaA06T0124700ZY
GSL_pool_high.bam.gz
GSL_pool_low.bam.gz
Janetzki_vs_zheyou73.bam.gz
Lorenz_vs_zheyou73.bam.gz
JS_leaves_28DAF_1.bam
JS_leaves_28DAF_2.bam
JS_leaves_28DAF_3.bam
JS_seeds_28DAF_1.bam
JS_seeds_28DAF_2.bam
JS_seeds_28DAF_3.bam
zheyou73.v0.gff3
BnaC08G0419000ZY
BnaC08T0206600ZY
BnaC05T0148400ZY
BnaA09T0559500ZY
BnaA08G0248700ZY
BnaA06T0124700ZY

### MYB51

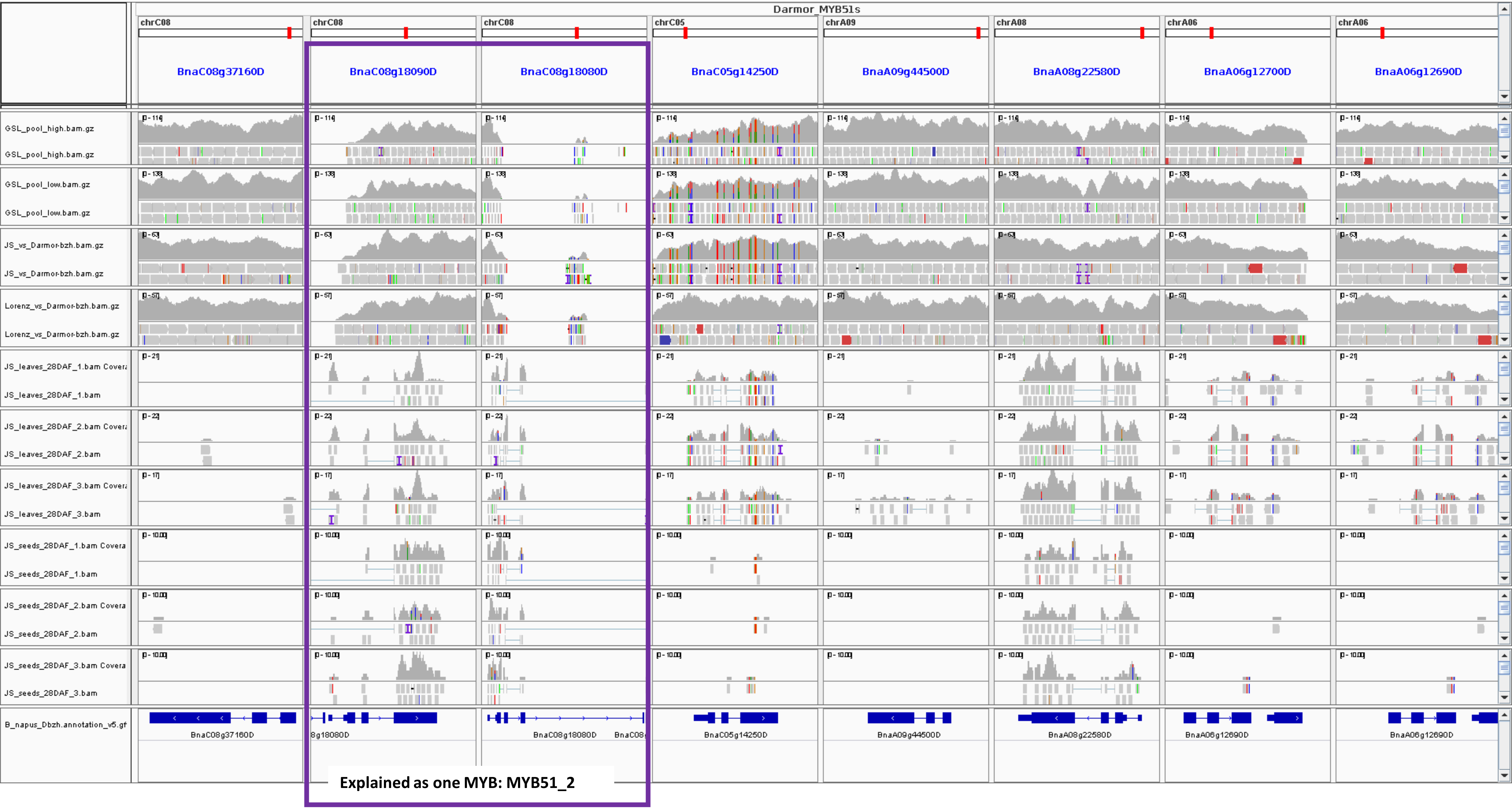
Darmor_MYB51s
chrC08
chrC08
chrC08
chrC05
chrA09
chrA08
chrA06
chrA06
BnaC08g37160D
BnaC08g18090D
BnaC08g18080D
BnaC05g14250D
BnaA09g44500D
BnaA08g22580D
BnaA06g12700D
BnaA06g12690D
GSL_pool_high.bam.gz
GSL_pool_low.bam.gz
JS_vs_Darmor-bzh.bam.gz
Lorenz_vs_Darmor-bzh.bam.gz
JS_leaves_28DAF_1.bam
JS_leaves_28DAF_2.bam
JS_leaves_28DAF_3.bam
JS_seeds_28DAF_1.bam
JS_seeds_28DAF_2.bam
JS_seeds_28DAF_3.bam
B_napus_Dbzh.annotation_v5.gf
Explained as one MYB: MYB51_2

### MYB51_2 and MYB51_3

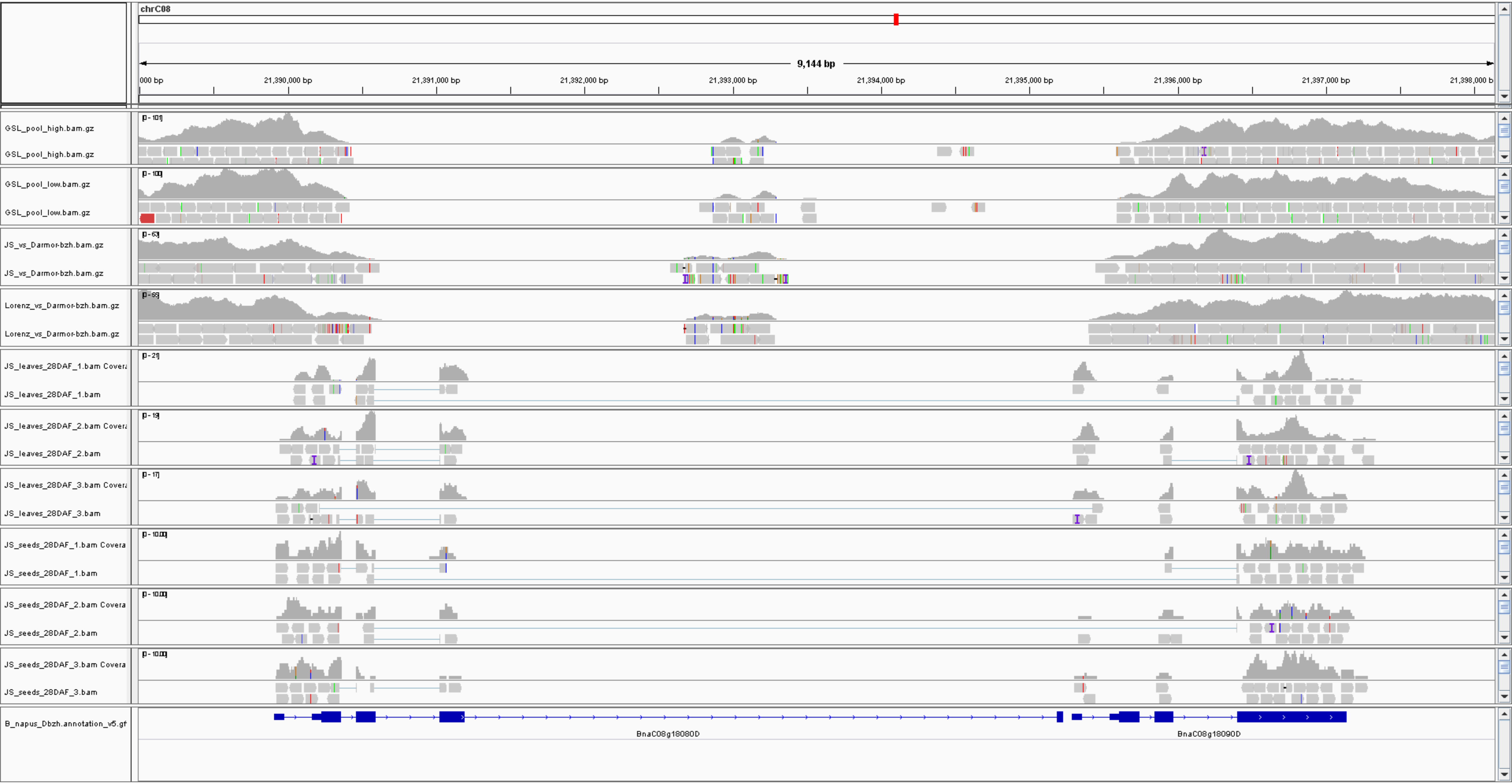


**Shows that the two annotated gene models in Darmor-bzh BnaC08g18080D and BnaC08g18090D likely explain one MYB, as supported by the RNA-Seq data**

### MYB122

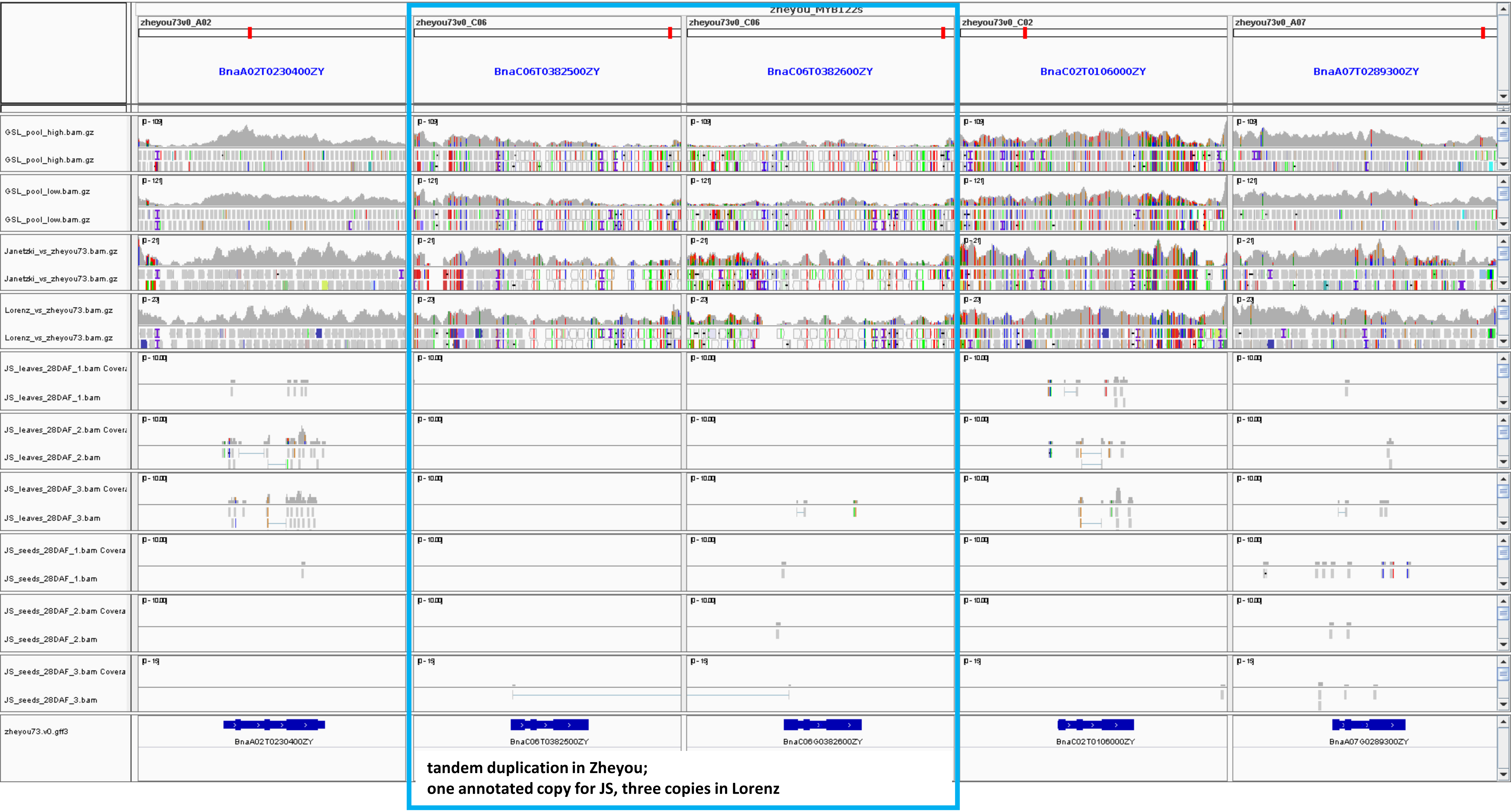
zheyou73v0_A02
zheyou73v0_C06
zheyou73v0_C06
zheyou73v0_C02
zheyou73v0_A07
BnaA02T0230400ZY
BnaC06T0382500ZY
BnaC06T0382600ZY
BnaC02T0106000ZY
BnaA07T0289300ZY
GSL_pool_high.bam.gz
GSL_pool_high.bam.gz
GSL_pool_low.bam.gz
GSL_pool_low.bam.gz
Janetzki_vs_zheyou73.bam.gz
Janetzki_vs_zheyou73.bam.gz
Lorenz_vs_zheyou73.bam.gz
Lorenz_vs_zheyou73.bam.gz
JS_leaves_28DAF_1.bam Covera
JS_leaves_28DAF_1.bam
JS_leaves_28DAF_2.bam Covera
JS_leaves_28DAF_2.bam
JS_leaves_28DAF_3.bam Covera
JS_leaves_28DAF_3.bam
JS_seeds_28DAF_1.bam Covera
JS_seeds_28DAF_1.bam
JS_seeds_28DAF_2.bam Covera
JS_seeds_28DAF_2.bam
JS_seeds_28DAF_3.bam Covera
JS_seeds_28DAF_3.bam
zheyou73.v0.gff3
BnaA02T0230400ZY
BnaC06T0382500ZY
BnaC06G0382600ZY
BnaC02T0106000ZY
BnaA07G0289300ZY
tandem duplication in Zheyou;
one annotated copy for JS, three copies in Lorenz

### MYB122

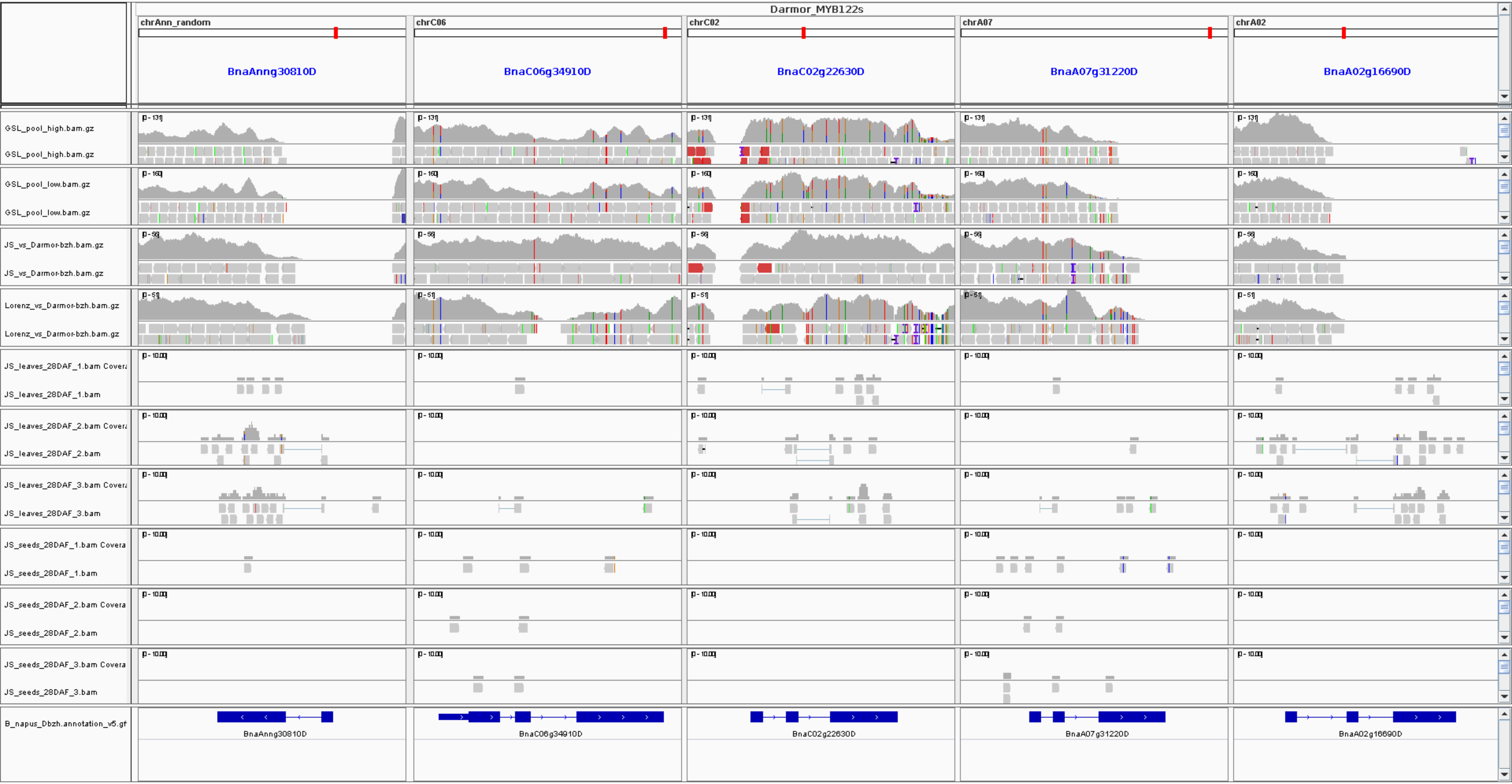
Darmor_MYB122s
chrAnn_random
chrC06
chrC02
chrA07
chrA02
BnaAnng30810D
BnaC06g34910D
BnaC02g22630D
BnaA07g31220D
BnaA02g16690D
GSL_pool_high.bam.gz
GSL_pool_low.bam.gz
JS_vs_Darmor-bzh.bam.gz
Lorenz_vs_Darmor-bzh.bam.gz
JS_leaves_28DAF_1.bam
JS_leaves_28DAF_2.bam
JS_leaves_28DAF_3.bam
JS_seeds_28DAF_1.bam
JS_seeds_28DAF_2.bam
JS_seeds_28DAF_3.bam
B_napus_Dbzh.annotation_v5.gf
