## Supplementary material for "Mapping-by-sequencing reveals genomic regions associated with seed quality parameters in *Brassica napus*": File S41

**A**

| | C09 homolog (*BnaMYB28_2*) | A09 homolog (*BnaMYB28_4*) | Phenotype |
|---|---|---|---|
| Janetzkis Schlesischer | *C09_MYB28_2_1** (ancestral allele) | absence | High GSL |
| SGDH14 | *C09_MYB28_2_1** (ancestral allele) | *A09_MYB28_4_1** (ancestral allele) | High GSL |
| Zheyou7 | *C09_MYB28_2_1** (ancestral allele) | *A09_MYB28_4_1** (ancestral allele) | High GSL |
| No2127 | *C09_MYB28_2_1** (ancestral allele) | absence | High GSL |
| Lorenz | *C09_MYB28_2_2** (insertion allele) | absence | Low GSL |
| Darmor-bzh | *C09_MYB28_2_2** (insertion allele) | absence | Low GSL |
| Express 617 | *C09_MYB28_2_2** (insertion allele) | absence | Low GSL |
| Westar | *C09_MYB28_2_2** (insertion allele) | absence | Low GSL |
| Gangan | *C09_MYB28_2_2** (insertion allele) | absence | Low GSL |
| Quinta | *C09_MYB28_2_2** (insertion allele) | absence | Low GSL |

**B**

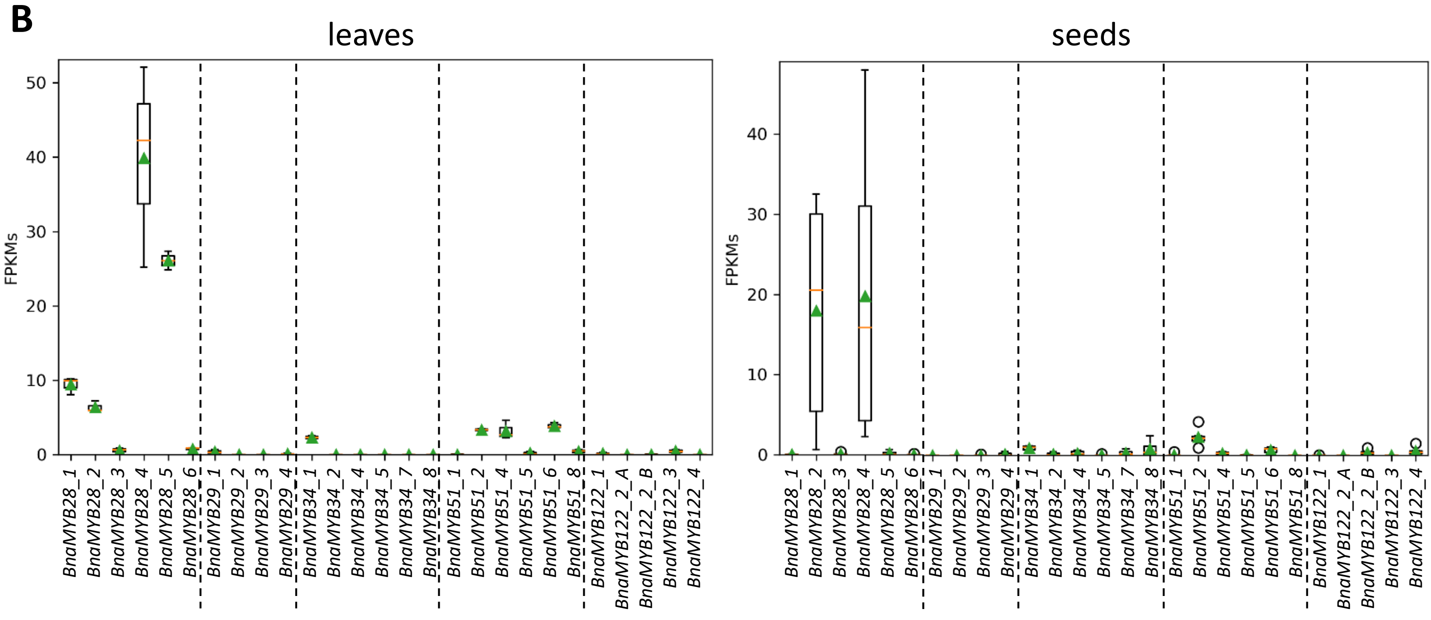


**File S41: The C09 *BnaMYB28_2* and A09 *BnaMYB28_4* alleles association with seed GSL phenotype across various *B. napus* cultivars.**

**Supplementary Text to File S41:** (A) The analysis was performed on the annotations of the assemblies derived from Song *et al.*, 2020. Moreover, the annotation of the Express 617 from Lee *et al.* 2020 was used. RNA-Seq data from seeds and leaves of Express 617 derived from Schilbert *et al.* 2021 were incorporated. SGDH14 seeds and leaves RNA-Seq data was used to construct a transcriptome assembly via Trinity v2.4.0 with default parameters and CDS sequences were extracted via TransDecoder to detect the *BnaMYB28* homeologs sequences. Seed GSL content information for each cultivars was derived from Song *et al.* 2020, Behnke *et al.* 2018 and the BnASSYST diversity panel. The recently published assembly of the low GSL cultivar Express 617 revealed the present of the insertion allele at the C09 homeolog, while the A09 homeolog is absent (File S35). Moreover, RNA-Seq data from leaves of Express 617 revealed the presence of the insertion on transcriptional level, which leads to a most likely truncated protein. Thus Express 617 revealed the same *BnaMYB28* homeolog alleles as Darmor-bzh, Lorenz, Westar, Gangan, and Quinta which are all low GSL lines as well. In addition, an in-depth investigation of four high GSL lines Janetzkis Schlesischer (P2), Zheyou7, SGDH14, and No2127 was performed. For Zheyou7 and SGDH14 both homeologs carry the ancestral alleles, leading to two functional *BnaMYB28* homeologs yielding a high seed GSL content. RNA-Seq data from seeds and leaves of SGDH14 support these hypotheses by revealing the ancestral alleles for both homeologs on transcriptional level. (B) Interestingly, as already observed from P2 seeds and leaves RNA-Seq data, the *BnaMYB28* homeolog of C09 was highly expressed compared to all other GSL MYBs. However, in contrast to P2, where the A09 *BnaMYB28* homeolog is absent, it is equally strongly expressed to the C09 homeolog in SGDH14 seeds. Thus, we assume if both homeologs are present and functional, they might contribute equally to seed GSL content. However, one functional and highly expressed *BnaMYB28* homeolog is sufficient to mark P2 as high GSL line.
